# Proximity labeling of protein complexes and cell type-specific organellar proteomes in Arabidopsis enabled by TurboID

**DOI:** 10.1101/629675

**Authors:** Andrea Mair, Shou-Ling Xu, Tess C. Branon, Alice Y. Ting, Dominique C. Bergmann

## Abstract

Defining specific protein interactions and spatially or temporally restricted local proteomes improves our understanding of all cellular processes, but obtaining such data is challenging, especially for rare proteins, cell types, or events. Proximity labeling enables discovery of protein neighborhoods defining functional complexes and/or organellar protein compositions. Recent technological improvements, namely two highly active biotin ligase variants (TurboID and miniTurboID), allowed us to address two challenging questions in plants: (1) what are *in vivo* partners of a low abundant key developmental transcription factor and (2) what is the nuclear proteome of a rare cell type? Proteins identified with FAMA-TurboID include known interactors of this stomatal transcription factor and novel proteins that could facilitate its activator and repressor functions. Directing TurboID to stomatal nuclei enabled purification of cell type- and subcellular compartment-specific proteins. Broad tests of TurboID and miniTurboID in Arabidopsis and *N. benthamiana* and versatile vectors enable customization by plant researchers.

## Introduction

All major processes of life, including growth and development and interactions among cells, organisms and the environment, rely on the activity and co-operation of hundreds of proteins. To fully understand these processes on a cellular level, we must know all players present in a cell or cell-type at a specific location and time. This requires information about transcription and chromatin state, as well as about protein abundance and protein complex compositions. A large international effort, the ‘human cell atlas’ project, is taking a first step in this direction. It aims to characterize all cell types in the human body, using recent advancements in high-throughput single-cell and multiplex techniques (Regev et al. 2017; Stuart and Satija 2019). Following this example, a call for a ‘plant cell atlas’ describing nucleic acid, protein and metabolite composition of different cell types in plants was issued at the start of this year (Rhee, Birnbaum, and Ehrhardt 2019). While several groups have produced single cell gene expression profiles (e.g. (Efroni et al. 2015; Ryu et al. 2019; Denyer et al. 2019; Nelms and Walbot 2019)) and tissue/cell-type specific profiles of active translation (e.g. (Vragovic et al. 2015; Tian et al. 2019)), we lack effective tools to obtain similarly precise information about protein distribution, abundance and the composition of protein complexes.

Today’s state-of-the-art for identification of *in-planta* protein interactors and complexes is affinity purification with protein-specific antibodies or single- and tandem affinity purification tags and subsequent mass spectrometry (MS) analysis (Struk et al. 2019; Xu et al. 2010). However, while affinity purification-mass spectrometry (AP-MS) strategies are undoubtedly useful in many instances, AP-MS is challenging for very low abundant proteins, those expressed only in rare cell types or developmental stages, and those with poor solubility like integral membrane proteins. Moreover, AP-MS tends to miss weak and transient interactions unless paired with crosslinking (Qi and Katagiri 2009; Van Leene et al. 2007; Bontinck et al. 2018). For obtaining subcellular proteomes, typical traditional approaches rely on cell fractionation protocols that enrich organelles from whole tissues, followed by protein extraction and MS analysis. Besides cross-contamination with other organelles, cell fractionation has the issue that only compartments that can be purified are accessible (Agrawal et al. 2011). The usefulness of this strategy for studying local protein compositions in individual cell types is further limited by the requirement of prior enrichment of the cell type of interest. For rare and transient cell types, acquiring a sufficient amount of ‘pure’ organelle material for MS analysis would be a major challenge.

Recent technological innovations in the form of proximity labeling (PL) techniques provide the sensitivity and specificity needed to identify protein complexes and (local) proteomes on a cell-type specific level. These techniques employ engineered enzymes to covalently attach biotin to nearby proteins which can then be affinity purified from total protein extracts using streptavidin-coupled beads without the need for crosslinking to stabilize weak and transient protein interactions or cell sorting and fractionation to enrich organelles. Because proteins do not have to be isolated in their native state, harsher extraction and more stringent wash conditions can be applied, which can reduce false positives from post-lysis interactions or from non-specific binding of proteins to the beads, and can improve solubilization of membrane proteins (for recent review see (Gingras, Abe, and Raught 2019)). An ever growing number of applications for PL in animals, including the characterization of protein complexes (e.g. the nuclear pore complex (Kim et al. 2014)), of organellar proteomes (e.g. mitochondrial matrix and intermembrane space (Rhee et al. 2013; Hung et al. 2016)) as well as for local proteomes (e.g. inside cilia (Mick et al. 2015), at synaptic clefts (Loh et al. 2016), at endoplasmic reticulum-mitochondria contact sites (Cho et al. 2017; Hung et al. 2017), demonstrate the usefulness and versatility of these techniques. Although the potential utility for PL in plants is equally tremendous, its adaptation in plants is proving challenging. So far, only three studies have described successful PL in plant systems, employing overexpression of a transcription factor (TF) in transiently transformed rice protoplasts (Lin et al. 2017) and high level expression of *Pseudomonas syringae* effector proteins in stable Arabidopsis lines (Khan et al. 2018) and transiently transformed *N. benthamiana* leaves (Conlan et al. 2018), respectively.

These plant studies all utilized BioID, which is based on an *E. coli* biotin ligase (BirA) made promiscuous by a point mutation (R118G) to yield BirA*. BirA* is either fused to a protein of interest or targeted to a desired subcellular localization to mark protein interactors and local proteomes, respectively. When biotin is supplied in the presence of ATP, BirA* binds and activates biotin and releases reactive biotinyl-AMP which can covalently bind to close-by primary amines on lysine residues (Roux et al. 2012). Based on experiments with the nuclear pore complex (Kim et al. 2014), the labeling radius of BirA* was estimated to be approximately 10 nm, which is in the size-range of an average globular protein. The actual labeling range may vary between experiments and is dependent on characteristics of the BirA* fusion protein, such as linker length and bait mobility, and on the labeling time. Typical labeling times with BirA* are between 12 and 24 hours, which limits the usefulness of BioID for studying short and time-sensitive events.

The three plant studies demonstrate that BioID can work in plants if the bait is expressed at sufficiently high levels, but the method does not perform nearly as well as it does in the animal systems for which it was designed. Several factors likely contribute to this (Bontinck et al. 2018). First, BioID is most often used in mammalian cell culture, where biotin is easily taken up from the growth medium. Structural features of plants, including cell walls and the cuticle, could impede biotin uptake, especially in older plants or certain tissues. Second, as an *E.* coli-derived enzyme, BirA*’s optimum temperature is about 37°C (Kim et al. 2016). Its activity at temperatures suitable for plant growth and treatment might be too low for efficient labeling. Finally, unlike animals, plants produce and store biotin in their cells, which hampers tight temporal control of labeling. Free biotin produced by the plant could lead to continuous low-level ‘background’ labeling prior to exogenous biotin application, which could over time lower the signal-to-noise ratio.

Recent engineering of improved versions of BirA, however, might improve PL efficiency in plants and provide the tools required to build a ‘plant cell atlas’. In a directed evolution approach, two new variants – TurboID and miniTurboID – with similar specificity as BirA* but greatly increased activity and lower temperature requirements were created (Branon et al. 2018). TurboID and miniTurboID were successfully utilized to generate different organellar proteomes in HEK cells and worked *in vivo* in a broad range of species, including Drosophila and *C. elegans* which were previously inaccessible for BioID (Branon et al. 2018). The efficacy of these new enzymes in plants and their potential to identify rare protein complexes and local proteomes in individual cell types of complex multicellular organisms, however, were not addressed.

## Highlights

- TurboID (T_ID_) and miniTurboID (miniT_ID_) work well in all tested tissues and growth stages of stably transformed Arabidopsis and in transiently transformed *N. benthamiana* leaves.
- Labeling times of under 10 minutes can give immunoblot-detectable signals, but longer incubation may be required for protein identification by mass spectrometry (MS).
- In Arabidopsis, T_ID_ activity is higher than miniT_ID_ activity, but ‘background’ labeling with endogenous biotin is also increased.
- Biotin concentrations in the range of 2.5-50 μM and 20-50 μM are suitable for enhanced labeling with T_ID_ and miniT_ID_. For most Arabidopsis tissues, submergence in the biotin solution is sufficient but some tissues and other plants may require vacuum infiltration of the biotin solution for optimal labeling.
- T_ID_ and miniT_ID_ work at temperatures compatible with normal plant growth and at elevated temperatures, but are most likely not suitable for cold stress experiments.
- Proximity labeling (PL) with the FAMA-T_ID_ fusion protein led to the identification of new putative co-activator and -repressor complex components for FAMA, a transcription factor in young guard cells.
- PL with nuclear T_ID_ produced general and young guard cell-specific proteomes with high specificity for nuclear proteins and identified guard cell specific transcription factors.
- Important considerations for the experimental design of PL experiments: choice of biotin ligase (T_ID_ vs. miniT_ID_); proper controls to distinguish specific labeling from background; optimization of labeling conditions (biotin concentration, treatment time); optimization of bead amount for affinity purification
- Important considerations for affinity purification and MS identification of biotinylated proteins: depletion of free biotin to reduce required bead amount (buffer exchange with gel filtration columns or centrifugal filters, dialysis); beads for affinity purification (avidin-, streptavidin-, or neutravidin beads vs. anti-biotin antibodies); MS sample prep (on-bead trypsin digest vs. elution and in-gel digest); MS and quantification method (label-free vs. isotopic labeling)

Here, we show that TurboID and miniTurboID enable effective PL in plants and have greatly improved activity compared to BirA*. We use TurboID to identify partners of the stomatal guard cell transcription factor FAMA and to obtain the nuclear proteome of a rare cell type in Arabidopsis seedlings – young stomatal guard cells. Our work indicates high *sensitivity* of the new PL enzymes in plants. To enable adoption by the plant research community, we provide reagents and a broadly applicable workflow for PL experiments under different experimental conditions in a variety of tissues in Arabidopsis and *N. benthamiana* and highlight critical steps in experimental design and execution.

## Results

### TurboID and miniTurboID can biotinylate plant proteins under conditions appropriate for plant growth

With increased efficiency, BioID-based PL would be a valuable tool to study protein interactions and local proteomes on a cell type-specific level in plants. We therefore made an initial diagnosis of whether the improved BirA* variants TurboID and miniTurboID (hereafter called T_ID_ and miniT_ID_; also abbreviated as TbID and mTb in animal literature) are appropriate for PL applications in plants, by testing their activity in the nucleus and cytosol of two plant model systems: transiently transformed *N. benthamiana* leaves and young seedlings of stably transformed Arabidopsis. To enable a comparison with previous experiments in the literature, we also included the original BirA* in our experiments and expressed all three versions under the ubiquitous UBIQUITIN10 (UBQ10) promoter. A YFP tag was added to confirm correct expression and localization of the T_ID_ and miniT_ID_ biotin ligases (Figure 1 – figure supplements 1 and 2).

To test biotin ligase activities, we treated leaf discs from *N. benthamiana* leaves or whole five-day old transgenic Arabidopsis seedlings, each expressing T_ID_ or miniT_ID_, with biotin and subsequently monitored biotinylation in total protein extracts using streptavidin immunoblots. For biotin treatment, we briefly vacuum infiltrated the plant tissue with the biotin solution and then incubated the tissue submerged in the solution for one hour at room temperature. Mock- or untreated plants were used as controls (Figure 1A). In T_ID_- and miniT_ID_-expressing *N. benthamiana* and Arabidopsis, biotin treatment induced strong labeling of proteins, demonstrating that the new biotin ligases work under our chosen conditions in plants. As was observed in other organisms, both T_ID_ and miniT_ID_ showed greatly increased activity compared to BirA*, which mainly achieved weak self-labeling within one hour of biotin treatment (Figure 1B-C, Figure 1 – figure supplements 3 and 4). Since plants produce and store free biotin in their cells, we were concerned about ‘background’ labeling in the absence of exogenous biotin. Although it did appear, background labeling was in most cases negligible. Direct comparison of T_ID_ and miniT_ID_ in our plant systems revealed little difference in either activity or background labeling in *N. benthamiana* (Figure 1B, Figure 1 – figure supplement 3), possibly due to the high expression levels of the constructs. In Arabidopsis, however, T_ID_ was clearly more active than miniT_ID_ but also produced more background, especially in lines with lower expression levels (Figure 1C, Figure 1 – figure supplement 4). Comparing nuclear and cytosolic constructs, we did not observe any significant differences in labeling efficiency at the resolution of immunoblots (Figure 1 – figure supplements 3 and 4).

**Figure 1:**
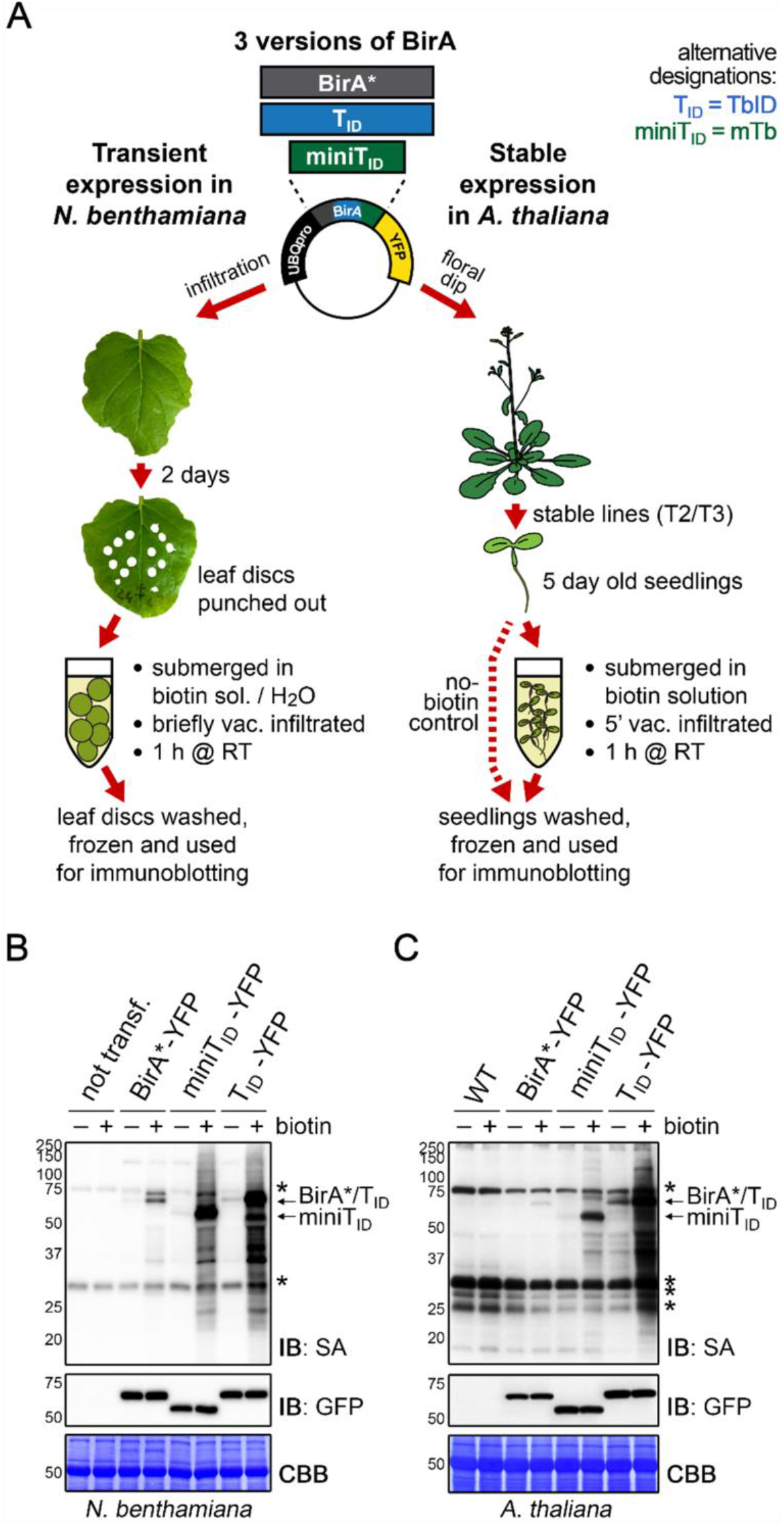
T_ID_ and miniT_ID_ exhibit robust biotinylation activity in *N. benthamiana* and Arabidopsis. **(A) Overview of the experimental setup**. UBQ10pro::BirA*/T_ID_/miniT_ID_-YFP constructs with an NLS or NES for nuclear or cytosolic localization were used for transient and stable transformation of *N. benthamiana* and *A. thaliana,* respectively. Tobacco leaf discs or whole Arabidopsis seedlings were submerged in a 250 μM biotin solution, briefly vacuum infiltrated, incubated for one hour at room temperature and frozen. Untreated controls were infiltrated with H_2_O or frozen directly. Expression and activity of the BirA versions were analyzed by immunoblotting. **(B-C) Biotin ligase activity in *N. benthamiana* (B) and Arabidopsis (C)** Streptavidin (SA) and anti-GFP immunoblots (IB) of protein extracts from tobacco leaf discs and Arabidopsis expressing the cytosolic BirA variants without (-) and with (+) biotin treatment. Untransformed tobacco leaves and Col-0 wild-type (WT) seedlings were used as controls. Each sample is a pool of 3 leaf discs or ~ 30 seedlings. Coomassie Brilliant Blue-stained membranes (CBB) are shown as a loading controls. Asterisks mark the positions of naturally biotinylated proteins. For microscopy images showing the subcellular localization of the BirA variants in *N. benthamiana* and Arabidopsis see Figure 1 – figure supplements 1 and 2. For immunoblots showing the activity and expression of both cytosolic and nuclear BirA versions in *N. benthamiana* and Arabidopsis see Figure 1 – figure supplements 3 and 4. For a schematic overview over the generation and composition of the available vectors in the ‘PL toolbox’ see Figure 1 – figure supplement 5.

From these experiments, we conclude that both T_ID_ and miniT_ID_ are well suited for use in plants. Which version is more suitable may depend on the individual question and whether high sensitivity (T_ID_) or tighter control over labeling time (miniT_ID_) is important. For this current study, we generated a versatile set of gateway-compatible entry and destination vectors that can be used to express T_ID_ or miniT_ID_ alone or as fusion with a protein of interest under a promoter of choice (Figure 1 – figure supplement 5). This ‘toolbox’ will be accessible through Addgene upon publication (vectors are listed in the materials and methods section).

### Testing boundaries with TurboID – effects of labeling time, temperature, biotin concentration and application

Achieving an optimal enzyme efficiency by using the right experimental conditions, like labeling time, temperature, biotin concentration and mode of application, can be key for using PL with low-abundant proteins in plants. We therefore tested the effect of those parameters on biotin labeling in four to five day old Arabidopsis seedlings expressing T_ID_ and miniT_iD_ under the UBQ10 promoter. In mammalian cell culture, 10 minutes of labeling with T_ID_ were sufficient to visualize biotinylated proteins by immunoblot and to perform analysis of different organellar proteomes (Branon et al. 2018). Using immunoblots, we observed similarly fast labeling in plants. T_ID_ induced labeling of proteins over background levels within 15 minutes of biotin treatment and labeling steadily increased over the next three to five hours (Figure 2A, Figure 2 – figure supplement 1). An increase in self-labeling of T_ID_ was evident even earlier, after as little as five minutes (compare Figure 4 – figure supplement 3). Time course experiments in *N. benthamiana* suggest that miniT_ID_ is equally fast, with clear labeling of proteins visible within 10 minutes (Figure 1 – figure supplement 3). This is a significant improvement over BirA*, for which labeling times of 24 hours were applied in all three published plant experiments (Khan et al. 2018; Conlan et al. 2018; Lin et al. 2017).

**Figure 2:**
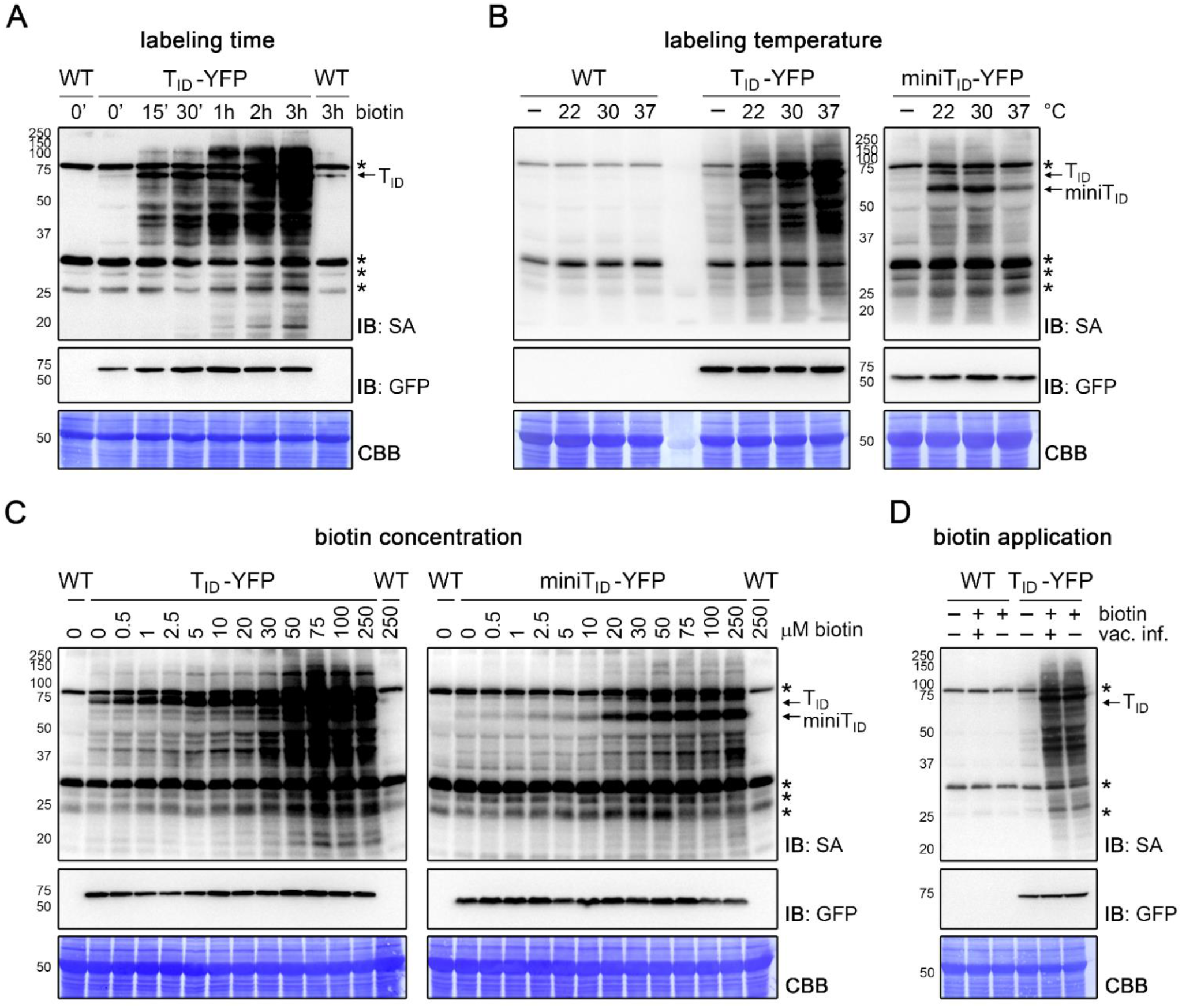
T_ID_ and miniT_ID_ work quickly and tolerate a range of experimental conditions in Arabidopsis seedlings. **(A-D)** Dependency of T_ID_ and miniT_ID_ activity on labeling time, temperature, biotin concentration and biotin application. Four to five day old seedlings were treated with biotin as described below. Activity and expression of the T_ID_/miniT_ID_-YFP constructs were analyzed by immunoblots (IB) with streptavidin-HRP (SA) and anti-GFP antibodies. Coomassie Brilliant Blue-stained membranes (CBB) are shown as a loading controls. Asterisks mark the positions of naturally biotinylated proteins. Each sample is a pool of ~ 30-50 seedlings. **(A) Labeling time**. Wild-type (WT) and UBQ10pro::T_ID_-YFP_NLS_ (T_ID_-YFP) seedlings were submerged in 250 μM biotin, briefly vacuum infiltrated and incubated for the indicated time at room temperature (22°C). A control sample was taken before treatment (0’). **(B) Temperature-dependency**. WT, UBQ10pro::T_ID_-YFP_NLS_ (T_ID_-YFP) and UBQ10pro::miniT_ID-NES_YFP (miniT_ID_-YFP) seedlings were submerged in 250 μM biotin and incubated for one hour at the indicated temperature. Control samples (-) were incubated in H_2_O at 22°C. **(C) Biotin concentration**. UBQ10pro::T_ID-NES_YFP (T_ID_-YFP) and UBQ10pro::miniT_ID-NES_YFP (miniT_ID_-YFP) seedlings were submerged in 0.5 to 250 μM biotin and incubated for one hour at room temperature. A control sample was taken before treatment (0 μM). **(D) Biotin application**. WT and UBQ10pro::T_ID_-YFP_NLS_ (T_ID_-YFP) seedlings were submerged in 250 μM biotin, briefly vacuum infiltrated (vac. inf.) or not and incubated for one hour at room temperature. A control sample was taken before treatment. For a longer time course in Arabidopsis and quantification of the immunoblots shown in (C) see Figure 2 – figure supplements 1 and 3. For a short time course and temperature dependency of T_ID_ and miniT_ID_ in *N. benthamiana* see Figure 1 – figure supplement 3 and Figure 2 – figure supplement 2.

We systematically tested the effect of different biotin treatment temperatures on T_ID_ and miniT_ID_ activity in Arabidopsis seedlings. Encouragingly, the activity of both variants was nearly as high at room temperature (22°C) as at 30°C. Moreover, T_ID_ showed only a moderate increase of activity at 37°C, while miniT_ID_ activity was actually reduced at this temperature (Figure 2B). High activity at ambient temperatures was also observed in *N. benthamiana* (Figure 2 – figure supplement 2). Increasing temperatures above plant growth conditions to improve labeling is therefore not needed.

The biotin concentration used for PL is an important consideration. Endogenous levels of biotin in plants are sufficient for low-level labeling of proteins by T_ID_, and to some extent also by miniT_ID_. While this may be useful for some applications, most applications will require strongly enhanced and time-regulated labeling through the addition of exogenous biotin. Although using excessive amounts of biotin is inconsequential for immunoblots, it poses a problem for downstream protein purification with streptavidin beads, as will be discussed later. We therefore tested biotin concentrations ranging from 0.5 to 250 μM to determine the optimal substrate concentration for T_ID_ and miniT_ID_. We found that T_ID_ has a larger dynamic range than miniT_ID_. Weak over-background labeling could already be seen with 0.5 μM biotin, which increased weakly through 20 μM, followed by a steeper increase with 30μM and was more or less saturated at 50-75 μM. MiniT_ID_ required between 2.5 and 10 μM biotin for weak activity, showed a steep increase with 20 μM (comparable to T_ID_) and was also saturated at 50-75 μM (Figure 2C, Figure 2 – figure supplement 3). T_ID_ and miniT_ID_ are therefore comparable to BirA* in their biotin requirement and concentrations of 2.5-50 μM (T_ID_) and 20-50 μM (miniT_ID_) seem to be appropriate.

In initial experiments, we vacuum infiltrated the plant material with the biotin solution to maximize biotin uptake. At least in Arabidopsis seedlings, this is not necessary. Simply submerging the plantlets in the biotin solution resulted in the same amount of labeling as vacuum infiltration followed by incubation in the biotin solution did (Figure 2D). This finding is very important since it not only simplifies handling of the experiment, but also improves isolation of labeled proteins by reducing the amount of free biotin in the tissue.

### TurboID works in a wide variety of developmental stages and tissues

For T_ID_ to be widely applicable it must be able to biotinylate proteins in many developmental stages and plant tissues. One initial concern, especially with T_ID_, was that background labeling from endogenous biotin would accumulate over time, making experiments with older tissues unfeasible. This was, however, not the case. Labeling worked well in 4 to 14 day old plate-grown seedlings without significant increase of background (Figure 3 – figure supplement 1). The same was true for separated roots and shoots of 6 to 14 day old seedlings and even for rosette leaves and flower buds of adult Arabidopsis plants grown on soil (Figure 3, Figure 3 – figure supplements 2 and 3). Background activity was low, especially in leaf tissue, and labeling worked well. Vacuum infiltration was not required for the tested plant sample types, except for unopened floral buds, where infiltration improved labeling relative to submergence in the biotin solution (Figure 3). This is likely because petals and reproductive tissues are not in direct contact with the biotin solution. Overall, our experiments suggest that T_ID_ will be applicable in a wide range of developmental stages and tissues. Since T_ID_ and miniT_ID_ behaved similar in most experiments, it is likely that the same is true for miniT_ID_.

**Figure 3:**
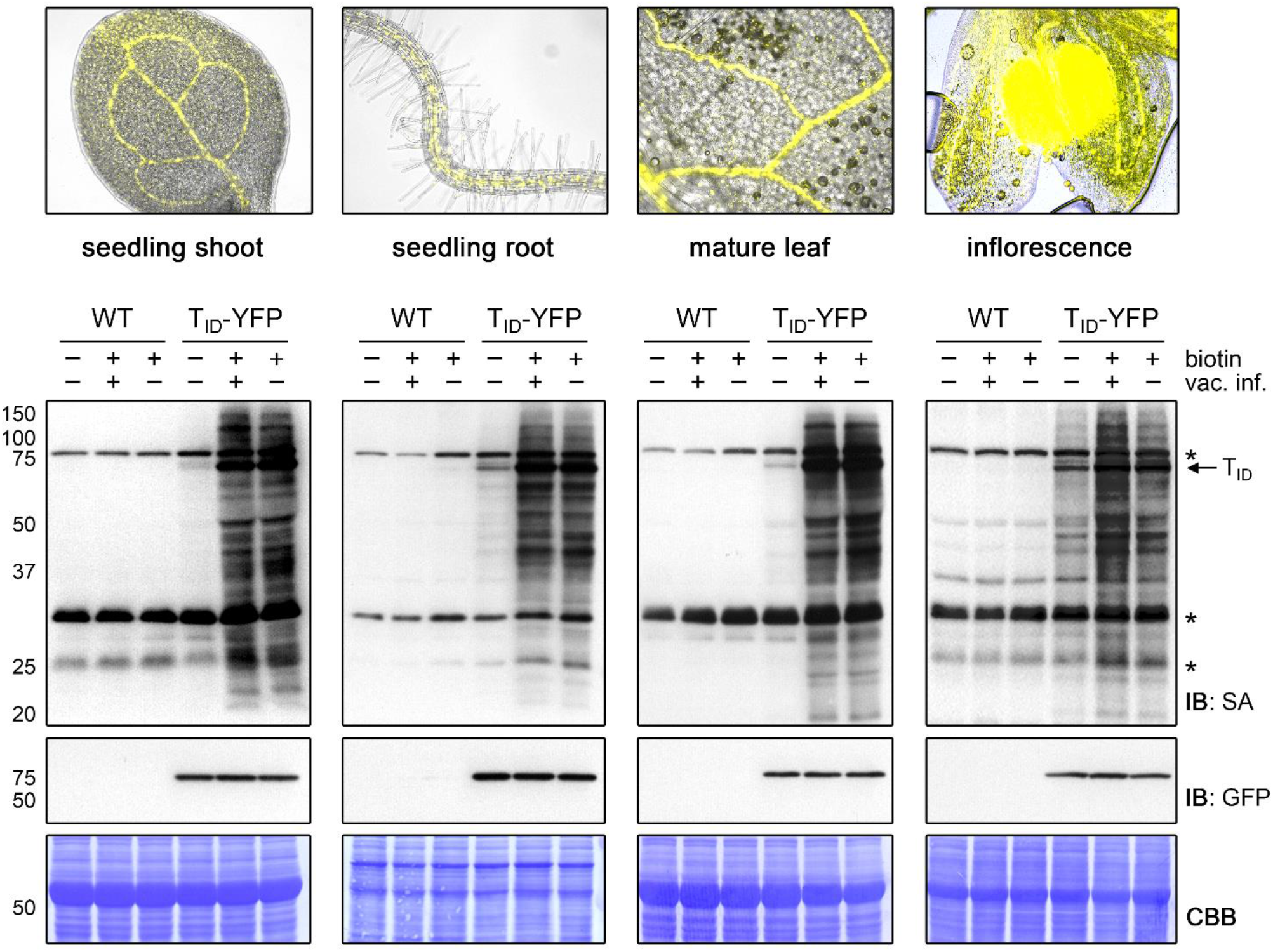
T_ID_ works in different developmental stages and organs of Arabidopsis and does not require vacuum infiltration of biotin. T_ID_ activity in shoots and roots of 10 day old plate-grown UBQ10pro::T_ID_-YFP_NLS_ (T_ID_-YFP) seedlings, and in rosette leaves and unopened flower buds of mature soil-grown plants. Col-0 wild-type (WT) was used as control. The plant material was submerged in a 250 μM biotin solution, briefly vacuum infiltrated until air spaces were filled with liquid or not vacuum infiltrated and incubated at room temperature for one hour. Control samples were taken before biotin treatment. Samples are pools of three shoots or roots, two rosette leaves or four inflorescences. Activity and expression of T_ID_-YFP were analyzed by immunoblots (IB) with streptavidin-HRP (SA) and anti-GFP antibodies. Coomassie Brilliant Blue-stained membranes (CBB) are shown as a loading controls. Asterisks mark the positions of naturally biotinylated proteins. Epifluorescence images of seedlings and mature tissues of the T_ID_-YFP line are shown on top. For immunoblots showing T_ID_ activity and background in four to 14 day old whole seedlings and shoots and roots of six to 14 day old seedlings see Figure 3 – figure supplements 1 and 2. For further microscopy images of the T_ID_-YFP line see Figure 3 – figure supplement 3.

### Testing TurboID’s potential to identify partners of a very low-abundant TF and to explore the nuclear proteome of a rare and transient cell type

After confirming general applicability of T_I_D for PL in plants, we wanted to test its performance for the identification of rare protein complexes and the characterization of cell type-specific organellar proteomes in a real experiment. For this purpose, we chose a cell type-specific transcription factor (FAMA) and a subcellular compartment of a rare cell type (nuclei of FAMA-expressing stomatal cells) for a case study. FAMA is a nuclear basic helix-loop-helix (bHLH) TF that is expressed in young stomatal guard cells (GCs) in the epidermis of developing aerial tissues (Ohashi-Ito and Bergmann 2006). The low abundance of FAMA and FAMA-expressing cells renders identification of interaction partners and cell type-specific nuclear proteins by traditional methods challenging and makes it well suited for a proof-of-concept experiment. Potential FAMA interaction partners were previously identified in yeast-2-hybrid (Y2H) and bimolecular fluorescence complementation studies (Chen et al. 2016; Kanaoka et al. 2008; Lee, Lucas, and Sack 2014; Li, Yang, and Chen 2018; Matos et al. 2014; Ohashi-Ito and Bergmann 2006), but only few have been confirmed by *in vivo* functional data.

For our study, we generated plants expressing T_ID_ and a fluorescent tag for visualization under the FAMA promoter, either as a FAMA-protein fusion or alone with a nuclear localization signal (NLS) (Figure 4A, Figure 4 – figure supplement 1). By comparing proteins labeled in the FAMApro::FAMA-T_ID_-Venus (FAMA-T_ID_) and FAMApro::T_ID_-YFP_NLS_ (_FAMA_nucT_ID_) plants to each other and to wild-type (WT) plants and UBQ10pro::T_ID_-YFP_NLS_ (_UBQ_nucT_ID_) plants, we can test the ability of the system to identify (1) proteins in close proximity to FAMA (FAMA complexes), (2) the nuclear protein composition during the FAMA-cell stage, (3) the nuclear proteome in general, and (4) possible FAMA-stage specific nuclear proteins. The FAMA-T_ID_-Venus construct is functional, since it complements the seedling-lethal phenotype of the *fama-1* mutant (Figure 4 – figure supplement 2).

Determining suitable incubation times is crucial since too short an incubation can yield insufficient protein amounts for identification but excessive incubation could label the whole subcellular compartment. We therefore performed labeling time-courses with the FAMA-T_ID_ line, using immunoblots as a readout. FAMA auto-labeling could be observed after as little as five minutes but clear labeling of other proteins required approximately 15 to 30 minutes. Longer incubation led to further increase in labeling up to three hours, both in the form of stronger discrete bands and of diffuse labeling, but stayed more or less the same thereafter (Figure 4B, Figure 4 – figure supplement 3). Based on these observations, we chose 0.5 and 3 hour biotin treatments, after which we would expect abundant and most FAMA interactors to be labeled, respectively, for the ‘FAMA interactome’ experiment (Figure 4C). For the ‘nuclear proteome’ experiment (Figure 4D), only the longer 3 hour time point was used, since over-labeling of the compartment was not a concern. Accordingly, the FAMA-T_ID_, _FAMA_nucT_ID_ and WT lines were treated for 0, 0.5 and 3 hours, and the _UBQ_nucT_ID_ line only for 3 hours. As plant material, we chose seedlings five days post germination, which corresponds to a peak time in FAMA promoter activity, as determined empirically by microscopy. We used three biological replicates per sample. To make the datasets as comparable as possible, all steps preceding data analysis were done together for the two experiments, as described in the next section.

**Figure 4:**
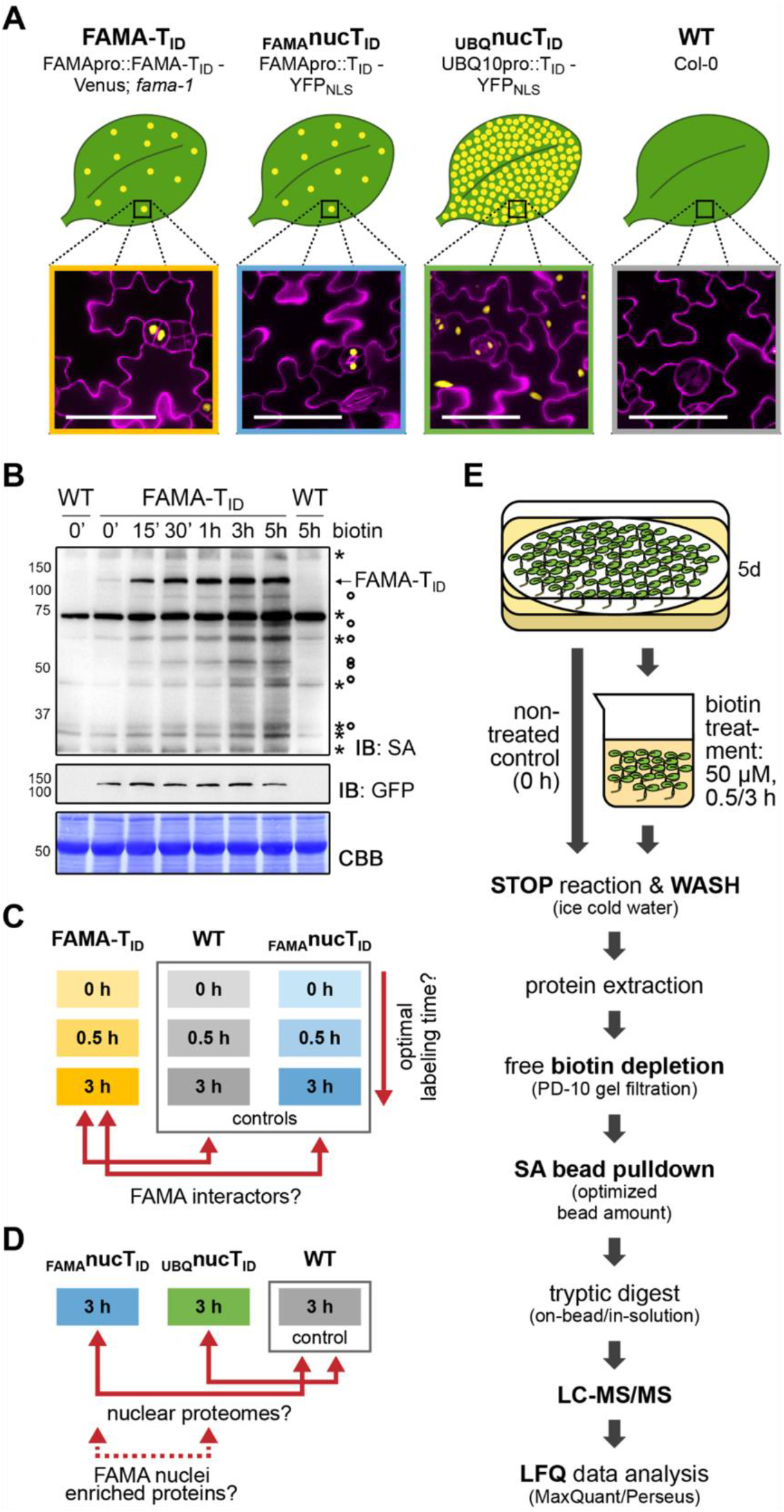
Testing T_ID_’s potential to label protein interactors and subcellular proteomes in a rare cell type in Arabidopsis. **(A) Plant lines generated for the ‘FAMA interactome’ and ‘nuclear proteome’ experiments**. Line names and genotypes are given on the top, schematic expression of the T_ID_ fusion proteins (yellow dots) in a leaf and confocal microscopy images of the epidermis of five day old seedlings are shown below (T_ID_ fusion protein = yellow; propidium iodide-stained cell wall = purple; scale bar = 50 μM). FAMA-T_ID_ and _FAMA_nucT_ID_ constructs are expressed in young guard cells, while the _UBQ_nucT_ID_ construct is expressed ubiquitously. **(B) Time course to optimize time points for the experiments.** Five day old wild-type (WT) and FAMA-T_ID_ seedlings were submerged in 250 μM biotin, briefly vacuum infiltrated and incubated for the indicated time at room temperature. Control samples were taken before treatment (0’). Samples are pools of ~ 30 seedlings. Activity and expression of FAMA-T_ID_ were analyzed by immunoblots (IB) with streptavidin-HRP (SA) and anti-GFP antibodies. The Coomassie Brilliant Blue-stained membrane (CBB) is shown as loading control. Asterisks and circles mark the positions of naturally biotinylated proteins and putative FAMA-T_ID_ targets, respectively. **(C-D) Scheme of samples and comparisons** used in the ‘FAMA interactome’ (C) and ‘nuclear proteome’ (D) experiments. **(E) Simplified workflow of the experimental procedure** from biotin labeling to protein identification by liquid chromatography coupled to mass spectrometry (LC-MS/MS). Three biological replicates were used. Abbreviations: SA, streptavidin; LFQ, label free quantification. For larger extracts of the confocal microscopy images of the plant lines used in the PL experiments shown in (A) see Figure 4 – figure supplement 1. For complementation of the *fama-1* phenotype by the FAMApro::FAMA-T_ID_-Venus construct see Figure 4 – figure supplement 2. For another labeling time course with the FAMA-T_ID_ line using shorter labeling times with and without vacuum infiltration of biotin see Figure 4 – figure supplement 3. For immunoblots showing successful labeling and purification of proteins for the ‘FAMA interactome’ and ‘nuclear proteome’ experiments see Figure 4 – figure supplements 4 and 5. For immunoblots demonstrating the importance of the biotin depletion step and a comparison of different biotin depletion strategies see Figure 4 – figure supplements 6 and 7.

### From labeling to identification of biotinylated proteins – identifying critical steps

Through empirical testing of experimental conditions using the _UBQ_nucT_ID_ line, we identified steps and choices that have a big impact on success of protein purification and identification after PL with T_ID_. These include sample choice to maximize bait abundance, removal of free biotin, optimizing the amount of streptavidin beads for affinity purification (AP) and choosing among MS sample prep procedures. Below, we describe our experimental procedure (Figure 4E) and highlight key choices.

We first labeled five day old seedlings, by submerging them in a 50 μM biotin solution for 0, 0.5 or 3 hours, quickly washed them with ice cold water to stop the labeling reaction and to remove excess biotin and isolated total proteins for AP of biotinylated proteins. The protein extracts were then passed through PD-10 gel filtration columns to reduce the amount of free biotin in the sample before proceeding with AP using magnetic streptavidin beads. Successful labeling and purification was confirmed by immunoblots (Figure 4 – figure supplements 4 and 5). The inclusion of a biotin-depletion step was found to be critical as free biotin in the protein extracts competes with biotinylated proteins for binding of the streptavidin beads (Figure 4 – figure supplement 6). While for mammalian cell culture or rice protoplasts thorough washing of the cells seems to suffice for removal of free biotin, this is not the case for intact plant tissue (see also (Conlan et al. 2018; Khan et al. 2018)). Especially when large amounts of starting material and moderate amounts of biotin are used, little to none of the biotinylated proteins may be bound by the beads. To maximize the amount of purified proteins it is further advisable to determine the appropriate amount of beads required for each experiment. We used 200 μl beads for approximately 16 mg total protein per sample. This amount was chosen based on tests with different bead-to-extract ratios (Figure 4 – figure supplement 7) and was sufficient to bind most biotinylated proteins in our protein extracts, although the beads were slightly oversaturated by the highly labeled _UBQ_nucT_ID_ samples (Figure 4 – figure supplement 5D).

Following AP, we performed liquid chromatography coupled to tandem mass spectrometry (LC-MS/MS) analysis to identify and quantify the captured proteins. Tryptic digest for LC-MS/MS analysis was done on-beads, since test experiments revealed that elution from the beads using two different methods (Cheah and Yamada 2017; Schopp and Bethune 2018) and subsequent in-gel digestion of biotinylated proteins yielded significantly lower protein amounts and less protein identifications (data not shown). This apparent sample loss is caused by the strong biotin-streptavidin interaction, which allows for stringent washing conditions but also prevents efficient elution of biotinylated proteins from the beads. Notably, highly biotinylated proteins, which likely comprise the most interesting candidates, will interact with more than one streptavidin molecule and will be especially hard to elute. After MS analysis, we identified and quantified the proteins by label-free quantification and filtered for significantly enriched proteins. This part was done separately for the ‘FAMA interactome’ and ‘nuclear proteome’ experiments and is described in the following sections.

### Proximity labeling is superior to AP-MS for identification of candidate interactors of FAMA

FAMA acts as both an activator and repressor for hundreds of genes (Hachez et al. 2011), suggesting a need for coordinated action with other TFs, co-activators and – repressors (Matos et al. 2014). Identifying such proteins through classical affinity purification-mass spectrometry approaches is hampered by the low overall abundance of FAMA. Apart from INDUCER OF CBF EXPRESSION 1 (ICE1), which is a known heterodimerization partner of FAMA (Kanaoka et al. 2008), we failed to identify any transcriptional (co-)regulators by AP-MS with FAMA-CFP, despite the use of crosslinking agents and large amounts of plant material (15 g of four day old seedlings per sample). Moreover, less than 20% of the AP-MS-derived ‘candidates’ were predicted to be nuclear, and one quarter were chloroplast proteins (Figure 5 – figure supplement 1, source file Figure 5 – supplemental table 5). We therefore wanted to see if PL would improve the identification of biologically relevant FAMA interactors.

For the ‘FAMA interactome’ experiment we compared proteins purified from plants expressing the FAMA-T_ID_ fusion (FAMA-T_ID_) with proteins from WT and with proteins from plants expressing nuclear T_ID_ (_FAMA_nucT_ID_) after 0, 0.5 and 3 hours of biotin treatment. In total, we identified 2,511 proteins with high confidence (quantified in all three replicates of at least one sample). Principal component analysis (PCA) showed a clear separation of the samples by genotype and time point (Figure 5 – figure supplement 2). Despite this clear separation, the majority of proteins were common to all samples, including the untreated WT control (source file Figure 5 – supplemental table 1), indicating that a large proportion of identified proteins bound to the beads non– specifically, and underlining the importance of appropriate controls and a data filtering pipeline like the one described below.

**Table 1:**
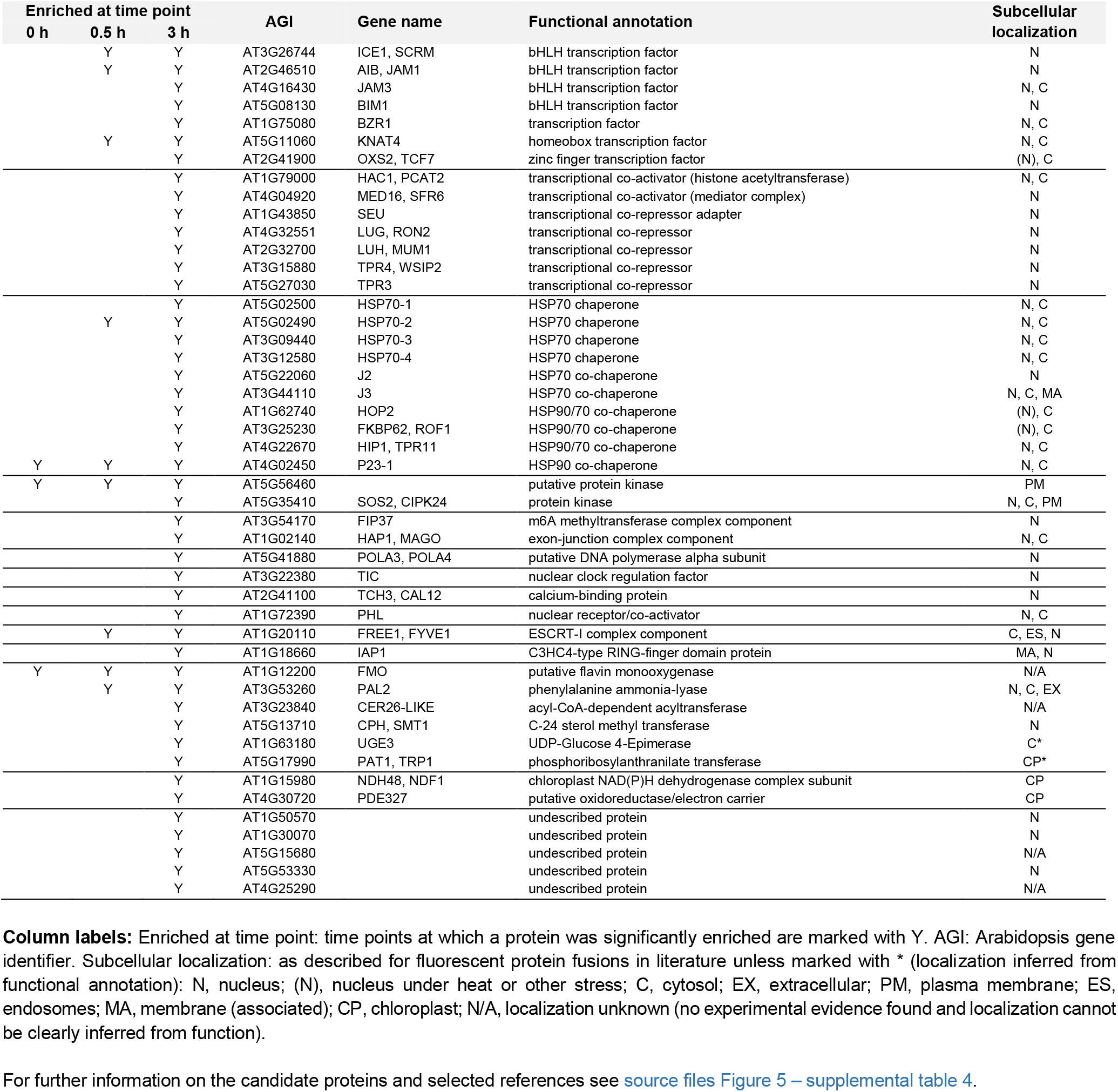
FAMA complex candidates from Figure 5

To narrow down the number of candidates and identify proteins enriched in the FAMA-T_ID_ plants, we applied several filtering steps (Figure 5). First, we identified proteins that were significantly enriched in the FAMA-T_ID_ samples compared to WT, considering only proteins that were found in all three replicates of FAMA-T_ID_. This resulted in a list of 73, 85 and 239 proteins (including FAMA) at the 0, 0.5 and 3 hour time points, respectively (Figure 5, source file – Figure 5 – supplemental table 2). Since T_ID_ is highly active and endogenous levels of biotin are sufficient for low-level labeling, there is a risk that proteins are labeled stochastically and that, over time, the whole nuclear proteome would be labeled. Notably, more than half of the proteins enriched in the FAMA-T_ID_ plants at any of the time points were also enriched in the _FAMA_nucT_ID_ plants (Figure 5, Figure 5 – figure supplement 3, source file – Figure 5 – supplemental table 2). We therefore filtered out proteins that were not significantly enriched in the FAMA-T_ID_ versus the _FAMA_nucT_ID_ samples, reducing the dataset to 6, 15 and 57 proteins (including FAMA) (Figure 5, source file – Figure 5 – supplemental table 2). Finally, we removed proteins that were not significantly enriched after biotin treatment compared to the untreated samples, since these proteins are likely genotype-specific contaminations.

**Figure 5:**
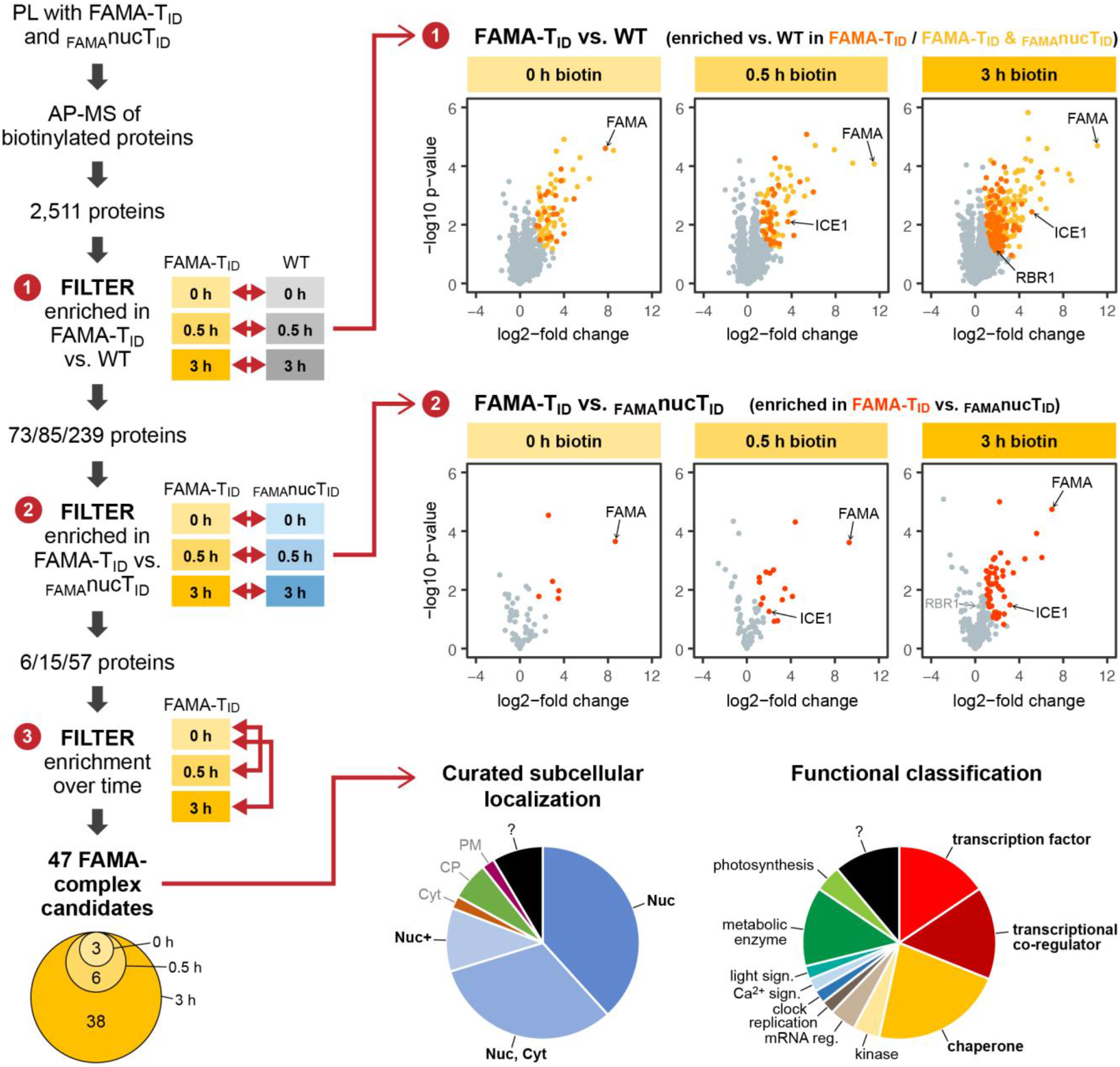
‘FAMA interactome’ experiment – PL with T_ID_ reveals potential FAMA interactors involved in transcriptional regulation Workflow (left) and results (right) of the experimental setup and data filtering process. Biotinylated proteins from seedlings expressing FAMA-T_ID_ or nuclear T_ID_ in FAMA-stage cells (_FAMA_nucT_ID_) and from wild-type (WT) after 0, 0.5 and 3 hours of biotin treatment were affinity purified (AP) with streptavidin beads and analyzed by mass spectrometry (MS). Proteins that were identified in all three biological replicates of at least one genotype and time point were used to filter for FAMA-complex candidates by three consecutive filtering steps: First, proteins enriched in FAMA-T_ID_ compared to WT were determined for each time point using unpaired 2-sided t-tests with a permutation-based FDR for multiple sample correction, considering only proteins that were identified in all three replicates of FAMA-T_ID_ at this time point (cutoff: FDR = 0.05, S0 = 0.5; ➊). Significant proteins were then filtered for enrichment compared to _FAMA_nucT_ID_ (same t-test parameters; ➋) and for enrichment in biotin-treated versus untreated FAMA-T_ID_ samples (t-test: p < 0.05; ➌). The Venn diagram on the bottom left shows the distribution of the 47 candidates between time points (see Table 1 for candidate list). Scatter plots on the right show log2-fold changes and −log10 p-values from t-test comparisons between FAMA-T_ID_ and WT (top) and FAMA-T_ID_ and _FAMA_nucT_ID_ (center). Proteins significantly enriched in FAMA-T_ID_ are shown in yellow, orange and red. All filtering steps and statistical analyses were done in Perseus. Subcellular localization and functional distribution of the candidate proteins is shown as pie charts on the bottom right. Data were manually curated from literature. Abbreviations: Nuc, nucleus; Cyt, cytosol, CP, chloroplast, PM, plasma membrane, +, and other. For hierarchical clustering and PCA of samples used in this experiment, see Figure 5 – figure supplement 2. For scatterplots and heatmaps of proteins enriched in FAMA-T_ID_ and _FAMA_nucT_ID_ compared to WT and for enrichment of proteins over time in all three genotypes, see Figure 5 – figure supplements 3 and 4. For tables summarizing all identified and enriched proteins, see source file Figure 5 – supplemental tables 1-4. For an overview over the workflow and proteins identified by affinity purification of FAMA-interacting proteins using a classical AP-MS strategy with GFP-Trap beads, see Figure 5 – figure supplement 1 and source file Figure 5 – supplemental table 5.

This left us with 47 ‘high confidence’ candidates (Figure 5, Table 1), 35 of which were previously demonstrated to be in the nucleus using fluorescent protein fusions or were found in MS-based nuclear proteome studies (Figure 5, Table 1, source file – Figure 5 – supplemental tables 2 and 4). Notably, more than half of the candidates have a role in regulation of transcription or are chaperones which could assist in FAMA’s role as a TF or in protein folding and stabilization, respectively (Figure 5, Table 1, source file – Figure 5 – supplemental table 4). Moreover, several of these proteins have previously been shown to interact with each other, which suggests that they could be part of the same FAMA complexes. This is a huge improvement compared to our AP-MS experiment, which could only confirm FAMA’s interaction with its obligate heterodimerization partner ICE1. The transcriptional regulators we found with PL can be roughly divided into two categories: TFs and transcriptional co-regulators. Among the TFs we again found ICE1 (as well as peptides shared between ICE1 and its orthologue SCRM2). We also found three other bHLH TFs (AIB/JAM1, JAM3, and BIM1) and the non-canonical bHLH-type TF BZR1. AIB and JAM3 play partially redundant roles in negative regulation of jasmonic acid (JA) signaling (Sasaki-Sekimoto et al. 2013; Fonseca et al. 2014), while BIM1 and BZR1 mediate brassinosteroid (BR) signaling (Yin et al. 2002; Wang et al. 2002). Both JA and BR signaling play roles in stomatal function or development (Acharya and Assmann 2009; Gudesblat et al. 2012; Kim et al. 2012).

Among the transcriptional co-regulators we found two significantly enriched transcriptional co-activators: MED16, which is part of the mediator complex that links TFs to RNA Pol II (Kidd et al. 2011), and HAC1, which is a histone acetyl transferase (HAT) (Deng et al. 2007). Combined with previous data showing a link between FAMA and RNA Pol II (Chen et al. 2016), this suggests that FAMA activates genes both directly by recruiting RNA Pol II and by opening up the chromatin for other transcriptional regulators. Among transcriptional co-repressors were TOPLESS (TPL)-related proteins TPR3 and TPR4 and LEUNIG (LUG) and LEUNIG HOMOLOG (LUH), which recruit histone deacetylases (HDACs) to TFs (Long et al. 2006). Additionally, we identified the linker protein SEUSS (SEU), which mediates interaction of LUG and LUH with TFs (Liu and Karmarkar 2008; Sitaraman, Bui, and Liu 2008). The identification of all three members of the SEU/LUG/LUH co-repressor complex is a strong indication of a functional complex with FAMA in the plant. Relaxing our filtering criteria to include proteins that are enriched in the FAMA-T_ID_ vs _FAMA_nucT_ID_ samples but were not significant under our stringent cutoff, we find several more components of transcriptional co-regulator complexes, including three more MED proteins, another HAT, two more TPL-related proteins and TPL itself.

RBR1, a cell cycle regulator and a known interactor of FAMA (Lee, Lucas, and Sack 2014; Matos et al. 2014), is also among the FAMA-T_ID_ enriched proteins but, due to a modest fold change, did not pass our last filters (Figure 5, source file – Figure 5 – supplemental table 2). This suggests that by setting a stringent cutoff on enrichment between FAMA-T_ID_ and _FAMA_nucT_ID_, we might lose some true interactors. This might be especially true of ubiquitously expressed proteins with many partners like RBR1, where FAMA-RBR1 interactions are likely to represent only a small fraction of all complexes.

Overall, this experiment demonstrates the usefulness of PL to identify potential interaction partners of rare proteins. We identified several good FAMA-complex candidates which could support FAMA in its role as a key TF and provide a possible mechanism for FAMA to induce fate-determining and lasting transcriptional changes in developing GCs. Some of the FAMA-complex candidates identified through PL are also slightly enriched in FAMA AP-MS samples compared to their controls. However, the enrichment is not enough to call any of them, except ICE1, significant in the AP-MS experiment. PL therefore not only gave us higher specificity for nuclear proteins than the AP-MS did, but it is potentially more sensitive as well. It is worth noting, that most FAMA interaction candidates were identified at the 3 hour time point and that longer biotin treatment greatly improved identification of biotinylated proteins (Figure 5, Figure 5 – figure supplement 4, source file – Figure 5 – supplemental table 3).

### Proximity labeling can be used to analyze the nuclear proteome in rare FAMA-expressing cells during GC development

The second question our PL experiment should answer was whether T_ID_ could be used to take a snapshot of the nuclear proteome of FAMA-expressing cells. Traditional tools to study organellar proteomes are not well-suited for such an endeavor, since they require isolation of the cell-type and organelle of interest and therefore lack the required sensitivity. Branon et al (Branon et al. 2018), showed that T_ID_ can be used to efficiently and specifically purify proteins from different subcellular compartments without prior cell fractionation. Their work was done using a homogeneous population of cultured mammalian cells, however, so it remained to be shown whether it would be possible to isolate an organellar proteome from an individual cell type, especially a rare or transient one, in a complex multicellular organism.

To identify nuclear proteins in FAMA-expressing young GCs and compare them to the global nuclear proteome at this growth stage, we purified proteins from seedlings expressing nuclear T_ID_ under the FAMA (_FAMA_nucT_ID_) and UBQ10 (_UBQ_nucT_ID_) promoter and from WT after three hours of biotin treatment. PCA and hierarchical clustering showed a clear separation of the three genotypes (Figure 6 – figure supplement 1). In total, we identified 3,176 proteins with high confidence (source file – Figure 6 – supplemental table 1). 1,583 proteins were significantly enriched in _UBQ_nucT_ID_ compared to WT (Figure 6, Figure 6 – figure supplement 2, source file Figure 6 – supplemental table 2), which is comparable to the number of proteins identified with nuclear T_ID_ in human HEK cells (Branon et al. 2018). These proteins comprise our ‘global’ nuclear protein dataset. Despite the relative rareness of FAMA-expressing cells, the _FAMA_nucT_ID_ dataset yielded 451 enriched proteins (Figure 6, Figure 6 – figure supplement 2, source file Figure 6 – supplemental table 3). Notably, most of them overlap with our global nuclear protein dataset (Figure 6), as would be expected since the _UBQ_nucT_ID_ dataset also contains FAMAstage cells.

**Figure 6:**
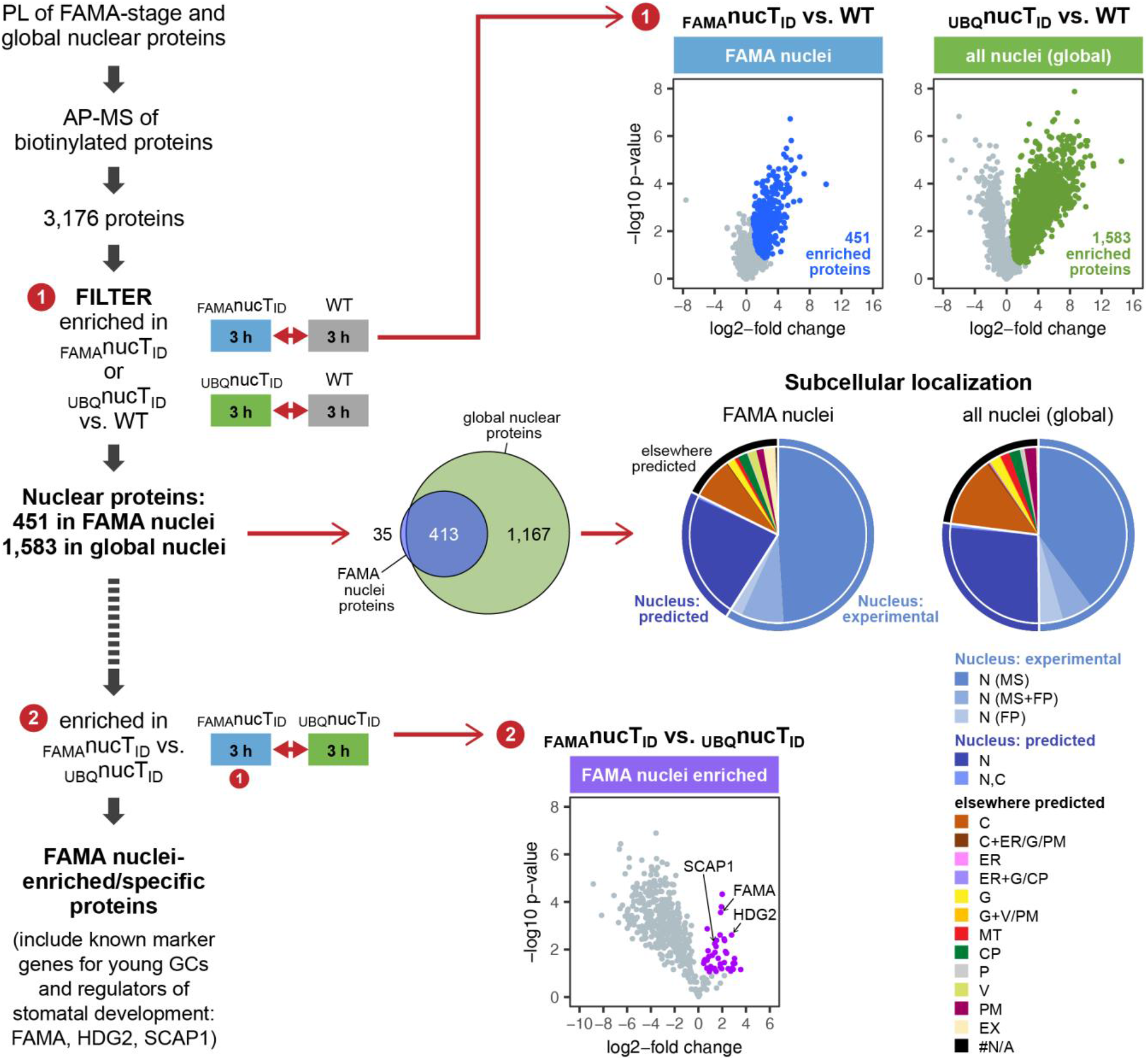
‘Nuclear proteome’ experiment – PL with nuclear T_ID_ results in identification of FAMA- and global nuclear proteomes with high organellar specificity. **Workflow (left) and results (right) of the experimental setup and data filtering process**. Biotinylated proteins from seedlings expressing nuclear T_ID_ under the FAMA (_FAMA_nucT_ID_) and UBQ10 (_UBQ_nucT_ID_) promoters and from wild-type (WT) after 3 hours of biotin treatment were affinity purified (AP) with streptavidin beads and analyzed by mass spectrometry (MS). Proteins that were identified in all three replicates of at least one genotype were used to filter for the FAMA cell- and global nuclear proteomes. Proteins enriched in _FAMA_nucT_ID_ or _UBQ_nucT_ID_ compared to WT were determined by unpaired 2-sided t-tests with a permutation-based FDR for multiple sample correction, considering only proteins that were identified in all three replicates of the respective nucT_ID_ line (cutoff: FDR = 0.05, S0 = 0.5; ➊). Scatter plots of the log2-fold changes and −log10 p-values from the t-test comparisons are on the top right. Significantly enriched proteins are shown in blue and green. Overlap of the enriched proteins and subcellular distribution are shown as Venn diagrams and pie charts on the center right. Proteins previously found in the nucleus in MS- or fluorescent protein (FP) fusion-based experiments, and proteins predicted to be in the nucleus (SUBA4 consensus prediction) are depicted in different shades of blue (N, nucleus; C, cytosol; CP, plastid; MT, mitochondrion; ER, endoplasmic reticulum; G, golgi; P, peroxisome; V, vacuole; PM, plasma membrane; EX, extracellular; #N/A, not available). Proteins that are highly enriched in or even specific to FAMA nuclei were identified through further filtering by comparing the abundance of the 451 FAMA nuclear proteins between _FAMA_nucT_ID_ and _UBQ_nucT_ID_ using a 2-sided t-test with an FDR of 0.01 as cutoff (➋). A scatterplot of the t-test results with significantly enriched proteins in purple is shown at the center bottom. For hierarchical clustering and PCA of samples used in this experiment, see Figure 6 – figure supplement 1. For scatterplots and heatmaps of proteins enriched in _FAMA_nucT_ID_ and _UBQ_nucT_ID_, see Figure 6 – figure supplements 2. For a visualization of enriched GO terms, see Figure 6 – figure supplement 3. For tables summarizing all identified and enriched proteins, published nuclear proteins and guard cell proteins, the localization data used for the pie charts in Figure 6, enriched GO terms of the nuclear proteomes and examples of identified proteins from different nuclear compartments, see source file Figure 6 – supplemental tables 1-8.

To estimate how ‘pure’ our nuclear proteomes are, we curated published nuclear and subnuclear compartment proteomes (Bae et al. 2003; Bigeard et al. 2014; Calikowski, Meulia, and Meier 2003; Chaki et al. 2015; Palm et al. 2016; Pendle et al. 2005; Sakamoto and Takagi 2013; Goto et al. 2019) and searched the Arabidopsis protein subcellular localization database SUBA (version 4, (Hooper et al. 2017), http://suba.live/) for proteins that were observed in the nucleus as fluorescent-protein fusions. This resulted in a combined list of 4,681 ‘experimentally determined nuclear proteins’; 4,021 from MS and 975 from localization studies (source file – Figure 6 – supplemental table 4). More than three quarters of the proteins enriched in our _UBQ_nucT_ID_ and _FAMA_nucT_ID_ datasets are either experimentally verified nuclear proteins or are predicted to be localized in the nucleus (Figure 6, source file Figure 6 – supplemental tables 2, 3 and 5). This suggests that most identified proteins are indeed nuclear proteins. Of the remaining proteins, most are predicted to be in the cytosol and could have been labeled by T_ID_ right after translation and before nuclear import of the biotin ligase or by a small mis-localized fraction of T_ID_. Importantly, chloroplast proteins, which are a major source of contamination in plant MS experiments, make up only about 3 % of our identified proteins (source file Figure 6 – supplemental table 5). In contrast, about 12 % and 6 % of the proteins identified in the two most recent Arabidopsis nuclear proteome studies (Palm et al. 2016; Goto et al. 2019), are predicted to be in the chloroplast (SUBAcon prediction, SUBA4). Gene ontology (GO) analysis is also consistent with nuclear enrichment in both nuclear T_ID_ datasets (Figure 6 – figure supplement 3, source file Figure 6 – supplemental tables 2,3 and 7). Importantly, our nuclear T_ID_ successfully labeled all major sub-nuclear compartments and domains, including the nuclear pore complex, the nuclear envelope, the nuclear lamina, the nucleolus and other small speckles, as well as DNA- and chromatin-associated proteins (subdomain markers from (Petrovska, Sebela, and Dolezel 2015; Tamura and Hara-Nishimura 2013), see source file Figure 6 – supplemental table 8 for examples).

After the general assessment of data quality and nuclear specificity, we asked whether proteins enriched in nuclei of FAMA-expressing cells, obtained by comparing the _FAMA_nucT_ID_ to the _UBQ_nucT_ID_ dataset (Figure 6, Figure 6 – figure supplement 2, source file Figure 6 – supplemental table 3), include known GC nuclear proteins. Indeed, looking at these proteins we find several known stomatal lineage- and GC-associated TFs, including FAMA itself, HOMEODOMAIN GLABROUS 2 (HDG2) and STOMATAL CARPENTER 1 (SCAP1) (Ohashi-Ito and Bergmann 2006; Peterson et al. 2013; Negi et al. 2013). Additionally, there were 10 proteins among the 42 highly _FAMA_nucT_ID_ enriched proteins, that were previously identified as part of the GC proteome by Zhao and colleagues using MS analysis of 300 million GC protoplasts (Zhao et al. 2010; Zhao et al. 2008) (source file Figure 6 – supplemental tables 3 and 6). It is worth noting that neither FAMA nor SCAP1 are in the Zhao GC protoplast proteome, presumably due to their relatively low expression and the use of material from more mature plants. These results suggest that PL can find lowly expressed proteins, that our knowledge of the GC proteome is not complete, and that additional important regulators of GC development and function might be uncovered by looking at stage-specific proteomes.

Overall, this experiment demonstrates the usefulness of T_ID_ as a tool for studying subcellular proteomes on a whole-plant as well as on a cell type-specific level. This will allow us to address questions that were previously inaccessible and thus has the potential to greatly improve our understanding of cellular processes in a cell-type specific context.

## Discussion

### Suitability of TurboID and miniTurboID for application in plants and performance of TurboID in FAMA-complex identification and nuclear proteome analysis

Our experiments presented in this study demonstrate that the new biotin ligase versions T_ID_ and miniT_ID_ drastically improve the sensitivity of PL in plants, compared to previously used BirA*, and tolerate a range of experimental conditions. We observed rapid labeling of proteins by T_ID_ and miniT_ID_ in different species, tissues and at different growth stages from seedlings to mature plants, using a simple biotin treatment protocol at room temperature. This greatly broadens the range of possible PL applications for plants and will allow to address hitherto inaccessible or hard to address questions in the future. To test the usefulness of T_ID_ for identification of protein interactors and organellar proteomes, we aimed to identify FAMA protein complexes and the nuclear proteome in FAMA-expressing cells. We deliberately picked a rare protein and cell type to observe its performance under conditions that make the use of traditional methods challenging or unfeasible.

In spite of FAMA’s low abundance, PL with the FAMA fusion protein in our ‘FAMA interactome’ experiment worked very well and outperformed our FAMA AP-MS experiments both in sensitivity and in specificity for nuclear proteins. Unlike in the AP-MS experiments, we identified a number of new proteins with gene regulatory functions that could support FAMA in its role as a master regulator of GC differentiation. Beyond the known dimerization partner ICE1, which was found in both types of experiments, PL identified four additional bHLH TFs with the potential to form alterative FAMA heterodimers as well as three non-bHLH TFs. We further identified several epigenetic regulators, which could fulfill roles predicted for FAMA complexes based on previous genetic and transcriptomic experiments (Adrian et al. 2015; Hachez et al. 2011; Matos et al. 2014) and fill a gap in our current model of gene regulation during GC formation. It was satisfying to see that when FAMA complex candidates were known to form functional complexes, we often identified multiple components of the complex. One example is the SEU-LUG/LUH co-repressor complex of which only SEU is known to interact directly with TFs (Liu and Karmarkar 2008). MED14, which was found as potential FAMA complex component using relaxed criteria, could also be part of this interaction chain. LUG is known to act by recruiting HDACs to promote epigenetic gene repression, but can also interact with mediator complex subunits like MED14 to interfere with their interaction with transcriptional activators (Liu and Karmarkar 2008). MED14, if indeed a FAMA complex component, could therefore either interact with the FAMA heterodimer or with the repressors LUG or LUH. It is likely that our candidates are part of different activating and repressive FAMA complexes. Which proteins are in which complex and which are direct FAMA interactors will need to be tested with independent methods.

Interestingly, we did not confirm all of FAMA’s previously postulated interaction partners in our PL experiment. This has both biological and technical causes. It is possible that some previously observed interactions are too rare or conditional (e.g. MED8 may require pathogen exposure) while others probably only happen under artificial conditions like in Y2H assays (e.g. bHLH71 and bHLH93, which seem not to be required for stomatal development (Ohashi-Ito and Bergmann 2006)). Enrichment of NRPB3 in FAMA-T_ID_ samples, on the other hand, could not be assessed because of non-specific binding of the protein to the beads. Other technical aspects that can complicate or prevent identification of interaction partners are removal of low-frequency interactions through stringent data filtering or a general lack of labeling. Because only proteins with exposed, deprotonated lysine residues can be labeled, sterically or chemically inaccessible proteins will not be detected.

PL of FAMA-expressing nuclei and comparison with a general nuclear proteome in our ‘nuclear proteome’ experiment showed that T_ID_ is suitable to capture sub-cellular proteomes at a cell type-specific level, even when the cell type is rare. We identified 1,618 proteins in both nuclear datasets combined, which is comparable to the most recent nuclear proteome obtained from cultured Arabidopsis cells (Goto et al. 2019). Judging from experimentally determined and predicted localization of the identified proteins and functional annotation with GO analysis, we obtained high specificity for nuclear proteins and identified proteins from all major nuclear subdomains. The biggest ‘contamination’ stems from the cytosol, which could be caused by a fraction of T_ID_ in the cytosol (e.g. right after translation) or from activated biotin diffusing out of the nucleus. Chloroplast-predicted proteins made up only a small fraction. Among FAMA-nuclei enriched proteins, we identified several known nuclear markers of young GCs, confirming our ability to detect cell type-specific proteins, as well as proteins that have not yet been linked to GC development or function but could be interesting to investigate further. One of them is SHL (SHORT LIFE), which is a histone reader that can bind both H3K27me3 and H3K4me3 histone marks and has been implicated in seed dormancy and flower repression (Mussig et al. 2000; Qian et al. 2018). Interaction of FAMA with RBR1 and with newly identified co-repressors and co-activators from this study, strongly suggests that chromatin marks are important to lock GC in their terminally differentiated state (Matos et al. 2014; Lee, Lucas, and Sack 2014) and SHL could be involved in this process.

### Considerations for a successful PL experiment

When designing a PL experiment, several things should be considered in order to achieve the best possible result. First, a choice has to be made which biotin ligase to use. Whether T_ID_ or miniT_ID_ is more suitable will depend on the research question. T_ID_ is more active, which is an advantage for low abundant proteins, for cases where over-labeling is not a concern or when labeling times should be kept as short as possible. MiniT_ID_ is less active in the presence of endogenous levels of biotin and will give less background labeling, which could be beneficial in tissues with higher than average endogenous biotin levels (Shellhammer and Meinke 1990) or when a more restricted and controlled labeling time is desired. Additional factors that should be considered when choosing a biotin ligase are whether one version works better in a specific subcellular compartment (as was the case in human HEK cells (Branon et al. 2018)) and whether the larger T_ID_ interferes with the activity or correct subcellular targeting of the protein of interest (POI). We added a fluorophore to all our biotin ligase constructs. This is very useful to confirm correct expression and subcellular localization of the T_ID_ or miniT_ID_ fusion protein, but may in some cases affect the activity of the ligase or the tagged protein. Interference with activity and targeting can depend on the position of the biotin ligase relative to the POI (N- or C-terminal tag). Another decision at the construct-design phase is the choice of linker length. For our experiments we added a short flexible linker to T_ID_. For identification of large protein complexes, increasing the linker length may improve labeling of more distal proteins.

Another crucial consideration are controls. Extensive and well-chosen controls are essential to distinguish between true candidates and proteins that either bind non-specifically to the beads or that are stochastically labeled because they are localized in the same subcellular compart as the POI. The former class of contaminants can be identified by including a non-transformed control (e.g. WT). The best control for the latter will be situation-dependent. For identifying interaction partners of a POI, one could use free T_ID_ or miniT_ID_ targeted to the same subcellular localization as the POI, as we have done in our ‘FAMA interactome’ experiment. Alternatively, one or more unrelated proteins that are in the same (sub)compartment but do not interact with the POI can be used. This strategy might improve identification of proteins that are highly abundant or ubiquitously expressed but of which only a small fraction interacts with the bait. Such proteins may be lost during data filtering if a whole-compartment control is used, as we observed for RBR1 in our ‘FAMA interactome’ experiment. In either case, the control construct should have approximately the same expression level as the POI. For sub-organellar proteomes (e.g a specific region at the plasma membrane) or for organellar proteomes in a specific cell-type, it is useful to label the whole compartment or the compartment in all cell-types for comparison.

Before doing a large-scale PL experiment, different experimental conditions should be tested to find a biotin concentration/treatment time/plant amount/bead amount combination that is suitable for the question and budget. Increasing the plant amount, labeling time or biotin concentration can improve protein coverage and can help to get more complete compartmental proteomes, but will also affect the amount of beads required. Excessive labeling is only advisable in closed compartments and when labeling of all proteins is the goal. Immunoblots are a useful tool to test different combinations of labeling concentration and time as well as for determining the correct bead amount for the pulldown so as not to over-saturate the beads. One should keep in mind, though, that immunoblots are more sensitive than MS and longer labeling times may be required for identification of labeled proteins by MS. Moreover, the signal intensity does not necessarily reflect the amount of labeled protein because highly biotinylated proteins have multiple binding sites for streptavidin, which leads to signal amplification, and labeling strength will also depend on the number of sterically/chemically available sites for biotinylation.

If biotinylation is induced by addition of exogenous biotin, a crucial step before AP of biotinylated target proteins is the depletion of free biotin in the sample to reduce the amount of beads required and therefore the per sample cost. We tested the effectiveness of two different approaches: gel filtration with PD-10 desalting columns (also used by (Conlan et al. 2018)) and repeated concentration and dilution with Amicon Ultra centrifugal filters (used by (Khan et al. 2018)). The latter method has the potential to remove more biotin and to be more suitable for large amounts of plant material. In our hands, though, using Amicon centrifugal filters led to considerable sample loss, presumably due to binding of the membrane, and was very slow. PD-10 columns, in contrast, did not lead to a notable loss of biotinylated proteins (Figure 4 – figure supplement 7). Surprisingly, consecutive filtering of protein extracts with two PD-10 columns did not improve the bead requirement. An alternative to these two methods is dialysis, which is suitable for larger volumes but is very time consuming. If the target is in an easy-to-isolate and sufficiently abundant organelle, cell fractionation prior to AP might also be considered to remove unbound biotin.

For AP, different kinds of avidin, streptavidin or neutravidin beads are available. Their strong interaction with biotin allows for efficient pulldowns and stringent wash conditions, but makes elution of the bound proteins difficult, especially if they are biotinylated at multiple sites. Should elution of the bound proteins be important for downstream processing or the identity of the biotinylated peptides be of major interest, the use of biotin antibodies might be preferable (Udeshi et al. 2017).

Finally, some consideration should also be given to the MS strategy. For example, digesting the proteins on the streptavidin beads instead of eluting them can increase the peptide yield. For data analysis, label-free quantification produces a more quantitative comparison of samples than comparison of peptide counts. Isotopic labeling, which allows samples to be analyzed together, can further improve quantitative comparison.

### Potential applications and challenges for PL in plants

We can see many potential applications for PL in plants, extending beyond two presented in this work. One that has the potential to be widely used is *in vivo* confirmation of suspected protein interactions or complex formation. The strategy is comparable to currently used co-immunoprecipitation (Co-IP) experiments but has the benefit that weak and transient, as well as other hard-to-purify interactions, are easier to detect. For this approach, proteins closely-related to the bait can be used as controls for interaction specificity.

One of the applications we demonstrate in this study is *de novo* identification of protein interaction partners and complex components. Extending from that, PL can be used to observe changes in complex composition in response to internal or external cues (e.g. stress treatment). Currently used techniques like peptide arrays, two-hybrid screens in yeast or plant protoplasts and AP-MS have the disadvantage that they are either artificial or work poorly for low abundant and membrane proteins and tend to miss weak and transient interactions. PL could overcome some of their deficiencies. One should keep in mind, however, that rather than identifying proteins bound directly to a bait protein, PL will mark proteins in its vicinity. Labeling is generally strongest for direct interactors, but labeling radius will depend on properties of the bait such as size, mobility and linker length as well as on the duration of labeling. To define a protein interaction network it will be useful to use several different baits (Gingras, Abe, and Raught 2019).

Another application of PL is characterization of subcellular proteomes, such as whole organellar proteomes, as we have demonstrated for the nucleus. Going forward, more detailed characterization of different organellar proteomes as well as sub-organellar proteomes and local protein composition, for example at membrane contact sites, can and should be addressed. This can be done on a whole plant level, but also at organ- and even cell type-specific level. Importantly, PL enables investigation of previously inaccessible compartments and of rare and transient cell types as we have demonstrated for FAMA-expressing GCs. Differences between individual cell-types or treatments can be investigated as well. Labeling times will, among other things, depend on the 2D or 3D mobility and distribution of the bait. For whole-compartment labeling, a combination of several baits may increase efficiency and protein coverage. One drawback of PL compared to traditional biochemical methods is that it requires the generation of transgenic plants, which limits its use to plants that can currently be transformed.

PL can also be used in combination with microscopy, to visualize the subcellular localization of biotinylated proteins and reveal labeling patterns of individual bait proteins. This can be utilized to confirm that labeling is restricted to the desired compartment, but it can also be used to fine-map the subcellular localization of a protein of interest or to obtain information about its topology, as was demonstrated for an ER transmembrane protein in human cells (Lee et al. 2016).

Extended uses of PL techniques that will require some modification of T_ID_ and miniT_ID_ or the PL protocol before they can be applied include interaction-dependent labeling of protein complexes (split-BioID), identification of RNAs associated with biotinylated proteins and identification of proteins associated with specific DNA or RNA sequences (for a recent review of PL methods describing these applications see (Trinkle-Mulcahy 2019)).

While there are is a plethora of questions that can be addressed with PL, there are also limitations to what will be possible. For example, although T_ID_ and miniT_ID_ are much faster than BirA*, controlled short labeling pulses (as are possible with APEX-based PL techniques) will be hard to achieve. T_ID_ and miniT_ID_ are always active and use endogenous biotin to continuously label proteins, preventing a sharp labeling start. In addition, exogenous biotin needs time to enter the plant tissue to initiate labeling. In mammalian cell culture, 10 minutes of labeling might be sufficient, but in a whole multicellular organism it may take much longer, depending on the experimental setup and how complete labeling should be. In our ‘nuclear proteome’ experiment, for example, three hours of biotin treatment were not sufficient to reach labeling saturation and only very few proteins were enriched in _FAMA_nucT_ID_ samples by 30 minutes biotin treatment (Figure 5 – figure supplement 4, source file Figure 5 – supplemental table 3). Development of strategies to reduce background labeling, for example by (conditional) reduction of endogenous biotin levels, could improve labeling time control in the future. Another limitation stems from temperature sensitivity. Although T_ID_ and miniT_ID_ work well at room temperature and elevated temperatures, they are inactive at 4°C (Branon et al. 2018) and thus likely incompatible with cold treatment as might be done for cold adaptation and cold stress experiments. Further, it is likely that some compartments will be harder to work in than others. Insufficient ATP and biotin availability and adverse pH or redox conditions could reduce T_ID_ and miniT_ID_ activity. For example, although the final step of biotin synthesis happens in mitochondria (Alban 2011), free biotin in mitochondria is undetectable (Baldet et al. 1993). It is possible that active biotin export from mitochondria will be a challenge for PL.

Going forward, it will be interesting to see how T_ID_ and miniT_ID_ perform for different applications and which challenges arise from the use in other plants, tissues and organelles. Our experiments in Arabidopsis and *N. benthamiana* suggest that PL will be widely applicable in plants and will provide a valuable tool for the plant community.

## Materials and methods

### Generation of the ‘PL toolbox’ vectors

#### Cloning of gateway-compatible entry vectors containing different BirA variants

BirA* (R118G), TurboID (T_ID_) or miniTurboID (miniT_ID_) were amplified from V5-hBirA(R118G)-NES_pCDNA3, V5-hBirA-Turbo-NES_pCDNA3 or V5-hBirA-miniTurbo-NES_pCDNA3 (Branon et al. 2018) using primers BirA-fw and BirA-rv or BirA-fw and BirA-NES-rv, thereby retaining the N-terminal V5 tag, either removing or retaining the C-terminal NES and adding a GGGGSGGG linker and an AscI restriction site to both ends. The T_ID_ and miniT_ID_ PCR products were then cloned into a pENTR vector containing YFP between the attL1 and L2 sites to generate YFP-T_ID_ and YFP-miniT_ID_ fusions or into a pDONR-P2R-P3 vector containing mVenus-STOP between the attR2 and L3 sites to generate T_ID_-mVenus-STOP and miniT_ID_-mVenus-STOP fusions by restriction cloning with AscI. The AscI site in the pDONR-P2R-P3 vector was first introduced by mutagenesis PCR using primers pDONR-mut-Ascl-fw and pDONR-mut-Ascl-rv. These vectors can be used for 3-way gateway recombination with a promoter and gene of choice as described below. For 2-way recombination with a promoter of choice, T_ID_ and miniT_ID_ (with and without NES) were further amplified from the pDONR-P2R-P3 plasmids described above using primers BirA-YFPnls-fw and BirA-YFPnls-rv to remove the N-terminal linker and to add a Notl and Ncol site at the N-and C-terminus, respectively. BirA* was directly amplified from the PCR products with BirA-fw and BirA-rv or BirA-NES-rv in the same way. The resulting PCR products were either cloned into a pENTR vector containing YFP followed by a stop codon (versions with NES) or a YFP followed by an NLS and a stop codon (versions without NES) between the attL1 and L2 sites using Notl and Ncol to generate BirA-_NES_YFP and BirA-YFP_NLS_ fusions (BirA = Bira*, T_ID_ or miniT_ID_). For a schematic overview over the cloning process and the composition of the vectors, see Figure 1 – figure supplement 5. For a list of vectors for 3- and 2-way recombination see the ‘PL toolbox’ vectors table at the end of the methods section.

#### Cloning of binary vectors for proximity labeling

The UBQ10pro::BirA-YFP_NLS_ (BirA = BirA*, T_ID_ or miniT_ID_) and FAMApro::T_ID_-YFP_NLS_ constructs were generated by LR recombination of the R4pGWB601 backbone (Nakamura et al. 2010) with either a pENTR5’/TOPO containing the 2 kb UBQ10 promoter or a pJET containing 2.4 kb of the FAMA (AT3G24140) upstream sequence (positions −2420 to −1, flanked by attL4 and R1, cloned from pENTR/D-TOPO (Ohashi-Ito and Bergmann 2006) by changing attL1 and L2 to L4 and R2) and the pENTR-BirA-YFP_NLS_ plasmids described above. The UBQ10pro::_NES_YFP-BirA constructs were generated by LR recombination of the R4pGWB601 backbone with a pENTR5’/TOPO containing the 2kb UBQ10 promoter and the pENTR-BirA-_NES_YFP plasmids described above. For a schematic overview over the cloning process and composition of the vectors, see Figure 1 – figure supplement 5. For a list of vectors, see ‘PL toolbox’ vectors table at the end of the methods section. The FAMApro::FAMA-T_ID_-Venus construct was generated by LR recombination of the pB7m34GW,0 backbone (Karimi, De Meyer, and Hilson 2005) with the pJET containing 2.4kb of the FAMA upstream sequence, with pENTR containing the genomic sequence of FAMA without stop codon flanked by attL1 and L2 (amplified from genomic DNA with primers gFAMA-fw and gFAMA-rv and recombined with pENTR/D-TOPO) and with pDONR-P2R-P3-T_ID_-mVenus (described above).

### Transformation and biotin activity assays in *N. benthamiana*

*N. benthamiana* ecotype NB-1 was transformed with UBQ10pro::BirA-YFP_NLS_ and UBQ10pro::BirA-_NES_YFP (BirA = BirA, T_ID_ or miniT_ID_) by infiltrating young leaves with a suspensions of Agrobacteria (strain GV3101) carrying one of the binary vectors. Agrobacteria were grown from an overnight culture for two hours, supplemented with 150 μM Acetosyringone, grown for another four hours, pelleted and resuspended in 5 % sucrose to an OD600 of 2. For more stable expression, Agrobacteria carrying a 35S::p19 plasmid (tomato bushy stunt virus (TBSV) protein p19) were coinfiltrated at a ratio of 1:1 for the temperature-dependency experiment. Two days after infiltration, expression was confirmed by epifluorescence microscopy and 5 mm wide leaf discs were harvested. Two to three discs were combined per sample. They were submerged in a 50 or 250 μM biotin solution, quickly vacuum infiltrated until the air spaces were filled with liquid and incubated at the indicated temperature for one hour. Control samples were not treated or were infiltrated with H_2_O. After biotin treatment, leaf discs were dried and flash-frozen for later immunoblotting. All experiments were done in duplicates with leaf discs for each of the two replicates taken from different plants if possible. Only one replicate is shown. Activity of different BirA variants was compared in four independent experiments, with similar results, temperature dependency of T_ID_ and miniT_ID_ was tested in two and one experiment, respectively.

### Arabidopsis lines used in this study

*Arabidopsis thaliana* Col-0 was used as wild-type (WT). The *fama-1* mutant line is SALK_100073 (Ohashi-lto and Bergmann 2006). Plant lines for testing the activity of BirA*, T_ID_ and miniT_ID_ (UBQ10pro::BiraA*-YFP_NLS_, UBQ10pro::T_ID_-YFP_NLS_, UBQ10pro::miniT_ID_-YFP_NLS_, UBQ10pro::BirA*-_NES_YFP, UBQ10pro::T_ID-NES_YFP, UBQ10pro::miniT_ID-NES_YFP) and for the ‘FAMA interactome’ and nuclear proteome’ experiments (FAMApro::FAMA-T_ID_-mVenus in *fama-1,* FAMApro::T_ID_-YFP_NLS_) were generated by floral dip of WT or *fama-1* +/- plants with the plasmids described above using agrobacterium strain GV3101. Selection was done by genotyping PCR *(fama-1)* and segregation analysis. We did not observe any obvious decrease in viability or developmental delay in our transgenic Arabidopsis plants. All lines had a single insertion event and were either heterozygous T2 or homozygous T3 or T4 lines. While screening for biotin ligase lines with the UBQ10 promoter, we observed that most regenerants had very weak YFP signal, especially the nuclear constructs. This was not observed with any of the cell-type specific promoters we tested. Lines used for the FAMA-CFP AP-MS experiments were previously described in other studies: FAMApro::FAMA-CFP (Weimer et al. 2018), SPCHpro::GFP_NLS_ and MUTEpro::GFP_NLS_ (Adrian et al. 2015).

### Plant growth conditions and biotin assays in Arabidopsis

Seeds were surface sterilized with ethanol or bleach and stratified for two to three days. For biotin treatment in whole Arabidopsis seedlings or roots and shoots, seedlings were grown on ½ Murashige and Skoog (MS, Caisson labs) plates containing 0.5% sucrose for four to 14 days under long-day conditions (16 h light/8h dark, 22°C). For treatment of rosette leaves and flowers, seedlings were transferred to soil and grown in a long-day chamber (22°C) until the first flowers emerged, at which point medium sized rosette leaves (growing but almost fully expanded) and inflorescences with unopened flower buds were harvested. All samples were pools from several individual plants. Biotin treatment was done by submerging the plant material in a biotin solution (0.5 – 250 μM biotin in water) and either vacuum infiltrating the tissue briefly until the air spaces were filled with liquid (approximately five minutes) or not, followed by incubation at room temperature, 30°C or 37°C for up to five hours. Controls were either treated with H_2_O or not treated. Following treatment, the plant material was dried and flash-frozen for later immunoblotting. To confirm reproducibility of the experiment, most experiments were done in duplicates (only one replicate shown) or repeated more than once with similar results. Comparison of all different BirA variants in Arabidopsis was done once with two independent lines for each NLS and NES constructs. Difference in activity and background between T_ID_ and miniT_ID_ matches other comparison done with varying temperature and biotin concentration. Comparison of T_ID_ and miniT_ID_ at different temperatures was done three times. Comparison of the biotin ligase activity with different biotin concentrations was done twice for T_ID_ and once for miniT_ID_. Three time courses with up to three or five hours of biotin treatment were done with the UBQ10pro::T_ID_-YFP_NLS_ line and time courses with the FAMA-T_ID_ line were done in duplicates. Experiments testing the effect of biotin application with and without vacuum infiltration in different tissues were done in duplicates. Activity of T_ID_ in 4-14 day old seedlings was tested in two independent experiments. Activity of T_ID_ in roots and shoots of 6-14 day old seedlings was tested once.

### Optimization of streptavidin (SA) pulldown conditions

#### Saturation of SA beads by free biotin

To test the impact of free biotin from biotin treatment on the affinity purification (AP) efficiency with SA-coupled beads, WT and UBQ10pro::T_ID_-YFP_NLS_ seedlings were submerged in H_2_O or 50 μM biotin for one hour, washed twice and frozen. AP was done in essence as described in (Schopp et al. 2017). Briefly, 1 ml of finely ground plant material was resuspended in an equal volume of ice cold extraction buffer (50 mM Tris pH 7.5, 500 mM NaCl, 0.4 % SDS, 5 mM EDTA, 1 mM DTT, 1 mM PMSF and 1x complete proteasome inhibitor), sonicated in an ice bath four times for 30 seconds on high setting using a Bioruptor UCD-200 (Diagenode) with 1.5-minute breaks on ice and supplemented with Triton-X-100 to reach a final concentration of 2 %. 2.3 ml of ice cold 50 mM Tris pH 7.5 were added to dilute the extraction buffer to 150 mM NaCl and samples were centrifuged at top speed for 15 minutes. The supernatant was mixed with 50 μl Dynabeads MyOne Streptavidin T1 (Invitrogen) that were pre-washed with equilibration buffer (50 mM Tris pH 7.5, 150 mM NaCl, 0.05 % Triton-X-100, 1 mM DTT) and incubated over night at 4°C on a rotor wheel. The beads were washed twice each with wash buffer 1 (2 % SDS), wash buffer 2 (50 mM Hepes pH 7.5, 500 mM NaCl, 1 mM EDTA, 1% Triton-X-100, 0.5% Na-deoxycholate), wash buffer 3 (10 mM Tris pH 8, 250 mM LiCl, 1 mM EDTA, 0.5% NP-40, 0.5 % Na-deoxycholate) and wash buffer 4 (50 mM Tris pH 7.5, 50 mM NaCl, 0.1 % NP-40). Protein were eluted by boiling the beads for 15 minutes at 98°C in elution buffer (10 mM Tris pH 7.5, 2% SDS, 5% beta mercaptoethanol, 2 mM biotin) and used for immunoblotting and SDS-PAGE. Two experiments with similar results were done.

#### Testing of biotin depletion strategies and optimization of the SA bead amount

To compare different biotin depletion strategies and determine the required bead amount for the PL experiments, five day old UBQ10pro::T_ID_-YFP_NLS_ seedlings were submerged in a 50 μM biotin solution for three hours, washed tree times with ice cold water, frozen and later used for biotin depletion and AP experiments. Proteins were extracted as described later in ‘Affinity purification of biotinylated proteins’. For biotin depletion with one or two PD-10 gel filtration columns (GE-Healthcare), the columns were equilibrated according to the manufacturer’s instructions with extraction buffer without PMSF and complete protease inhibitor (50 mM Tris pH 7.5, 150 mM NaCl, 0.1 % SDS, 1 % Triton-X-100, 0.5 % Na-deoxycholate, 1 mM EGTA, 1 mM DTT). 2.5 ml protein extract were loaded ono the column. For the trial with one PD-10 column, the gravity protocol was used, eluting with 3.5 ml equilibration buffer. For the trial with two PD-10 columns, the proteins were eluted from the first column with 2.5 ml equilibration buffer using the spin protocol. The flow through was applied to a second PD-10 column and eluted with 3.5 ml equilibration buffer using the gravity protocol. For biotin depletion with an Amicon Ultra-4 Centrifugal filter (Millipore Sigma), 625 μl protein extract were concentrated three times to 10 – 20% of the starting volume by centrifuging in a swinging bucket rotor at 4,000g and 4°C and diluted with extraction buffer to the initial volume. To test bead requirements, a volume equivalent to 1/5 of the amount used in the ‘FAMA interactome’ and ‘nuclear proteome’ PL experiments was incubated with five to 30 μl Dynabeads MyOne Streptavidin C1 (Invitrogen) over night at 4°C on a rotor wheel. The beads were either washed once with 1 M KCl and with 100 mM Na_2_CO_3_ and twice with extraction buffer (1x PD-10) or just twice with extraction buffer (Amicon Ultra, 2x PD-10) before elution of bound proteins by boiling for 10 minutes at 95°C in 4x Leammli buffer supplemented with 20 mM DTT and 2 mM biotin. Each biotin depletion strategy was tested once.

### ‘FAMA interactome’ and ‘nuclear proteome’ PL experiments

#### Seedling growth and biotin treatment

Approximately 120μl of seeds per sample and line were surface sterilized with ethanol and bleach and stratified for two days. The seeds were then spread on filter paper (Whatman Shark Skin Filter Paper, GE Healthcare 10347509) that was placed on ½ MS plates containing 0.5 % sucrose and grown in a growth chamber under long-day conditions for five days. For biotin treatment, seedlings were carefully removed from the filter paper, transferred into beakers, covered with 40 ml of a 50 μM biotin solution and incubated in the growth chamber for 0.5 or 3 hours. The biotin solution was removed and seedlings were quickly rinsed with ice cold water and washed tree times for three minutes in approximately 200 ml of ice cold water to stop the labeling reaction (Branon et al. 2018) and to remove excess biotin. The seedlings were then dried with paper towels, a sample was taken for immunoblots and the remaining seedlings were split into aliquots of about 6 to 7 g fresh weight for the three biological replicates, frozen in liquid nitrogen, ground to a fine powder and stored at −80°C until further use. Untreated 0 h samples were frozen directly after harvest. Treatment was timed to keep harvesting times as close together as manageable. Small samples for immunoblots were taken after treatment. The number of replicates was chosen to provide statistical power but to keep the experimental cost and sample handling in a feasible range.

#### Affinity purification of biotinylated proteins

For the affinity purification with streptavidin beads, 3 ml of densely packed ground plant material were resuspended in 2 ml extraction buffer (50 mM Tris pH 7.5, 150 mM NaCl, 0.1 % SDS, 1 % Triton-X-100, 0.5 % Na-deoxycholate, 1 mM EGTA, 1 mM DTT, 1x complete, 1 mM PMSF) and incubated on a rotor wheel at 4°C for 10 minutes. 1μl Lysonase (Millipore) was added to digest cell walls and DNA/RNA and the suspension was incubated on the rotor wheel at 4°C for another 15 minutes. The extracts were then distributed into 1.5 ml reaction tubes and sonicated in an ice bath four times for 30 seconds on high setting using a Bioruptor UCD-200 (Diagenode) with 1.5-minute breaks on ice. The suspension was centrifuged for 15 minutes at 4°C and 15,000 g to remove cell debris and the clear supernatant was applied to a PD-10 desalting column (GE healthcare) to remove excess free biotin using the gravity protocol according to the manufacturer’s instructions. Briefly, the column was equilibrated with five volumes of ice cold equilibration buffer (extraction buffer without complete and PMSF), 2.5 ml of the protein extract were loaded and proteins were eluted with 3.5 ml equilibration buffer. The protein concentration of the protein extract was then measured by Bradford (BioRad protein assay). The protein extract was diluted 1:5 to avoid interference of buffer components with the Bradford assay. A volume of each protein extract corresponding to 16 mg protein was transferred into a new 5 ml LoBind tube (Eppendorf) containing Dynabeads MyOne Streptavidin C1 (Invitrogen) from 200 μl bead slurry that were pre-washed with extraction buffer. Complete protease inhibitor and PMSF were added to reach final concentrations of 1x complete and 1 mM PMSF, respectively, and the samples were incubated on a rotor wheel at 4°C overnight (16 h). The next day, the beads were separated from the protein extract on a magnetic rack and washed as described in (Branon et al. 2018) with 1 ml each of the following solutions by incubating on the rotor wheel for 8 minutes and removing the wash solution: 2x with cold extraction buffer (beads were transferred into a new tube the first time), 1x with cold 1 M KCl, 1x with cold 100 mM Na_2_CO_3_, 1x with 2M Urea in 10 mM Tris pH 8 at room temperature and 2x with cold extraction buffer without complete and PMSF. 2 % of the beads were boiled in 50 μl 4x Laemmli buffer supplemented with 20 mM DTT and 2 mM biotin at 95°C for 5 minutes for immunoblots. The rest of the beads was spun down to remove the remaining wash buffer and stored at −80°C until further processing. Sample prep was done in three batches with one replicate of each sample in each batch. Samples from replicate 2 (batch 2) were used for immunoblots to confirm the success of the procedure.

#### MS sample preparation

For on-beads tryptic digest, the frozen streptavidin beads were thawed and washed twice each with 1 ml 50 mM Tris pH 7.5 (transferred to new tube with first wash) and 1 ml 2 M Urea in 50 mM Tris pH 7.5. The buffer was removed and replaced by 80 μl Trypsin buffer (50 mM Tris pH 7.5, 1 M Urea, 1m M DTT, 0.4 μg Trypsin). The beads were then incubated for three hours at 25°C with shaking, and the supernatant was transferred into a fresh tube. The beads were washed twice with 60 μl 1 M Urea in 50 mM Tris pH 7.5 and all supernatants were combined (final volume 200 μl). The combined eluates were first reduced by adding DTT to a final concentration of 4 mM and incubating at 25°C for 30 minutes with shaking and then alkylated by adding Iodoacetamide to a final concentration of 10 mM and incubating at 25°C for 45 minutes with shaking. Finally, another 0.5 μg Trypsin were added and the digest completed by overnight (14.5 h) incubation at 25°C with shaking. The digest was acidified by adding formic acid to a final concentration of ~ 1 % and desalted using OMIX C18 pipette tips (10 – 100 μL, Agilent). C18 desalting tips were first activated by twice aspiring and discarding 200 μl buffer B2 (0.1 % formic acid, 50 % acetonitrile) and equilibrated by four times aspiring and discarding 200 μl buffer A2 (0.1 % formic acid). Peptides were bound by aspiring and dispensing the sample eight times. Then, the tip was washed by 10 times aspiring and discarding 200 μl buffer A2 and the peptides were eluted by aspiring and dispensing 200 μl buffer B2 in a new tube for eight times. The desalted peptides were dried in a speed vac and stored at −80°C until further processing.

#### LC-MS/MS

For LC-MS/MS analysis, peptides were resuspended in 0.1 % formic acid. Samples were analyzed on a Q-Exactive HF hybrid quadrupole-Orbitrap mass spectrometer (Thermo Fisher), equipped with an Easy LC 1200 UPLC liquid chromatography system (Thermo Fisher). Peptides were separated using analytical column ES803 (Thermo Fisher). The flow rate was 300 nl/min and a 120-min gradient was used. Peptides were eluted by a gradient from 3 to 28 % solvent B (80 % acetonitrile, 0.1 % formic acid) over 100 minutes and from 28 to 44 % solvent B over 20 minutes, followed by short wash at 90 % solvent B. Precursor scan was from mass-to-charge ratio (m/z) 375 to 1,600 and top 20 most intense multiply charged precursors were selection for fragmentation. Peptides were fragmented with higher-energy collision dissociation (HCD) with normalized collision energy (NCE) 27.

#### MS data analysis – protein identification and label-free quantification

Protein identification and LFQ were done in MaxQuant (version 1.6.2.6) (Tyanova, Temu, and Cox 2016). Settings were default settings with minor modifications. Methionine oxidation and N-terminal acetylation were set as variable modifications and Carbamidomethylcysteine as fixed modification. Maximum number of modifications per peptide was five. Trypsin/P with a maximum of two missed cleavages was set as digest. Peptides were searched against the latest TAIR10 protein database containing a total of 35,386 entries (TAIR10_pep_20101214, updated 201108-23, www.arabidopsis.org) plus a list of likely contaminants containing Trypsin, human Keratin, streptavidin, YFP, T_ID_-mVenus and T_ID_-YFP and against the contaminants list of MaxQuant. Minimum allowed peptide length was seven. FTMS and TOF MS/MS tolerance were 20 ppm and 40 ppm, respectively, and the peptide FDR and protein FDR were 0.01. Unique and razor peptides were used for protein quantification. LFQ minimum ratio count was set to two and fast LFQ was active with a minimum and average number of neighbors of three and six, respectively. Match between runs and second peptides were checked. The datasets for the ‘FAMA interactome’ and ‘nuclear proteome’ experiments were searched separately. The mass spectrometry proteomics data (raw data and MaxQuant analysis) have been deposited to the ProteomeXchange Consortium (http://proteomecentral.proteomexchange.org) via the PRIDE partner repository (Vizcaino et al. 2013) and will be made accessible upon publication.

#### MS data analysis – identification of enriched proteins

Filtering and statistical analysis were done with Perseus (version 1.6.2.3) (Tyanova et al. 2016). The ‘proteinGroups.txt’ output file from MaxQuant was imported into Perseus using the LFQ intensities as Main category. The datamatrix was filtered to remove proteins marked as ‘only identified by site’, ‘reverse’ and ‘potential contaminant’. LFQ values were log2 transformed and proteins that were not identified/quantified in all three replicates of at least one time point of one genotype (low confidence) were removed. The clustering analyses shown in Figure 5 – figure supplement 2 and Figure 6 – figure supplement 1 were done at this point, using the built-in ‘Hierarchical clustering’ function with the standard settings (Euclidean distance with average linkage, no constraints and preprocessing of k-means). Next, missing values were imputed for statistical analysis using the ‘Replace missing values from normal distribution’ function with the following settings: width = 0.3, down shift = 1.8 and mode = total matrix and principal component analysis was done. Filtering steps to identify enriched proteins differed between the two experiments and are described below.

#### ‘FAMA interactome’ experiment

To identify proteins enriched in T_ID_-expressing samples versus the WT control, unpaired 2-sided Students t-tests were performed comparing the FAMA-T_ID_ or _FAMA_nucT_ID_ with the corresponding WT samples at each time point. The integrated modified permutation-based FDR (“Significance Analysis of Microarrays” (SAM) method) was used for multiple sample correction with an FDR of 0.05, an S0 of 0.5 and 250 randomizations to determine the cutoff. As an additional filter, only proteins that were quantified in all three replicates of the FAMA-T_ID_ or _FAMA_nucT_ID_ samples at this time point were tested (see source file Figure 5 – supplemental table 2 for number of proteins used in each test). Significantly enriched proteins were used for hierarchical clustering with standard settings, using Z-transformed non-imputed LFQ values. To further identify proteins that are significantly enriched in FAMA-T_ID_ versus _FAMA_nucT_ID_, corresponding t-tests were performed on the reduced dataset using the same parameters as for the first t-tests. Finally, to filter out proteins that were not enriched in at least one of the two treatments compared to untreated FAMA-T_ID_, unpaired 2-sided Students t-tests were performed on the remaining proteins comparing the samples at the 0.5 and 3 h time points to the 0 h time point using a p-value of 0.5 as cutoff.

#### Protein enrichment over time

To identify proteins enriched after short or long biotin treatment, unpaired 2-sided Students t-tests were performed comparing the 0.5 and 3 h treatment samples of WT, FAMA-T_ID_ and _FAMA_nucT_ID_ to the corresponding 0 h samples. An FDR of 0.05 and an S0 of 0.5 with 250 randomizations were chosen as cutoff for multiple sample correction. Only proteins that were quantified in all three replicates of at least one of the two groups were tested (see source file Figure 5 – supplemental table 3 for number of proteins used in each test).

#### ‘Nuclear proteome’ experiment

To identify proteins enriched in nuclear T_ID_-expressing samples versus the WT control, unpaired 2-sided Students t-tests were performed comparing the _UBQ_nucT_ID_ or _FAMA_nucT_ID_ with WT samples. An FDR of 0.05 and an S0 of 0.5 with 250 randomizations were chosen as cutoff for multiple sample correction. The FDR/S0 combination was chosen to get a minimum number of ‘false negatives’ (statistically enriched proteins in WT) in the _FAMA_nucT_ID_ versus WT comparison. Due to oversaturation of the streptavidin beads with the _UBQ_nucT_ID_ samples, there is an over proportional amount of proteins that more abundant in WT compared to _UBQ_nucT_ID_. Nevertheless, the same cutoff was used for both comparisons. As an additional filter, only proteins that were quantified in all three replicates of the _UBQ_nucT_ID_ or _FAMA_nucT_ID_ sample were tested (see source file Figure 6 – supplemental tables 2 and 3 for number of proteins used in each test). To further identify proteins that are highly enriched in FAMA nuclei compared to all nuclei, we looked for proteins that were equally or even more abundant in the _FAMA_nucT_ID_ than the _UBQ_nucT_ID_ samples, by performing unpaired 2-sided t-tests between _FAMA_nucT_ID_ and _UBQ_nucT_ID_ on the reduced dataset. As cutoff, an FDR of 0.01 and S0 of 0 (only the q-value determines significance) were used. Significantly enriched proteins were used for hierarchical clustering with standard settings, using Z-transformed non-imputed LFQ values.

#### Plots and heatmaps

Scatterplots and PCA plots were made in RStudio (version 1.1.463 (RStudio Team (2016)), R version 3.5.1 (R Core Team (2018)) using t-test and PCA results exported from Perseus. Hierarchical clustering was done in Perseus and the plots were exported as pdf.

#### Classification of the subcellular localization of nuclear proteins form PL

Proteins identified by PL with nuclear T_ID_ were divided into three broad categories: (1) previously identified in the nucleus experimentally, (2) not found experimentally but predicted to be in the nucleus and (3) predicted to be elsewhere. To determine proteins in the first category, a list of proteins identified in published nuclear- or subnuclear proteomics studies and of proteins detected in the nucleus in localization studies with fluorescent proteins (from SUBA4 (Hooper et al. 2017); http://suba.live/; date of retrieval: February 5 2019) was compiled (see source file Figure 6 – supplemental table 4). Localization predictions for proteins not identified experimentally in the nucleus was done using the SUBAcon prediction algorithm (Hooper et al. 2014) on SUBA4 (date of retrieval: February 5 2019). SUBAcon provides a consensus localization based on 22 different prediction algorithms and available experimental data.

#### Gene ontology analysis

Proteins that were significantly enriched in _FAMA_nucT_ID_ and _UBQ_nucT_ID_ compared to WT (FAMA- and global nuclear proteins) were used for GO analysis with AgriGO v2 (Tian et al. 2017) with the following settings: Singular enrichment analysis (SAE) with TAIR10_17 as background, statistical method: fisher, multitest adjustment: Yekutieli (FDR), significance level: 0.05, minimum number of entries: five. Significantly enriched GO terms from AgriGO v2 (see Source file 6 – supplemental table 7) were visualized using REViGO (Supek et al. 2011) with the following settings: list size medium, associated numbers are p-values, GO term size: Arabidopsis thaliana, semantic similarity measure: SimRel. The R script provided by REViGO was used to draw the TreeMap plot in RStudio. Very small labels were removed and label size was adjusted in Adobe Illustrator.

### Microscopy

Brightfield and epifluorescence images of *N. benthamiana* leaves and Arabidopsis seedlings, leaves and flowers were taken with a Leica DM6B microscope using a Leica CRT6 LED light source. Confocal microscopy images of Arabidopsis seedlings expressing different biotin ligase constructs were taken with a Leica SP5 microscope. For confocal microscopy, cell walls were stained with propidium iodide (Molecular Probes) by incubating in a 0.1 mg/ml solution for three to five minutes. Images were processed in FIJI (ImageJ) (Schindelin et al. 2012). Several independent Arabidopsis lines and transiently transformed *N. benthamiana* leaves were analyzed. Images shown in figures and figure supplements are representative.

### Immunoblots

Samples for immunoblots were prepared by resuspending frozen and ground plant material from biotin treatment assays with 1x Leammli buffer (60 mM Tris pH 6.8, 2 % SDS, 10 % glycerol, 2.5 % beta-mercaptoethanol, 0.025 % bromphenol blue) or mixing protein extracts 1:1 with 2x Leammli buffer and boiling the samples for five minutes at 95°C. Proteins bound to SA or GFP-T rap beads were eluted from the beads as described in the respective methods sections. Proteins were separated by SDS-PAGE and blotted onto Immobilon-P PVDF membrane (0.45 μm, Millipore) using a Trans-Blot Semi-Dry transfer Cell (BioRad). The following antibodies were used: Streptavidin-HRP (S911, Thermo Fisher Scientific), Rat monoclonal anti-GFP antibody (3H9, Chromotek), AffiniPure Donkey Anti-Rat IgG (712-035-153, Jackson Immuno Research Laboratories). Blots were probed with primary antibodies overnight at 4°C or up to one hour at room temperature and with the secondary antibody for one hour at room temperature and incubated with ECL Western blotting substrates according to the manufacturer’s instructions. Signals were detected on X-ray films or on a ChemiDoc MP Imaging System (BioRad).

### FAMA-CFP AP-MS experiments

#### Seedling growth and crosslinking

For the two FAMA-CFP AP-MS experiments, FAMApro::FAMA-CFP complementing the homozygous *fama-1* mutant and two lines expressing nuclear GFP under an early and late stomatal lineage specific promoter (SPCHpro::GFP_NLS_ and MUTEpro::GFP_NLS_) were used. Large amounts of seedlings were grown for four days in liquid culture in ½ MS + 0.5 % sucrose (AP-MS experiment 1) or on filter paper that was placed on ½ MS + 0.5 % sucrose plates (AP-MS experiment 2). Seedlings were then treated with the proteasome inhibitor MG-132 by submerging them in liquid ½ MS + 0.5 % sucrose and 10 μM MG-132 for 2.5 (AP-MS 1) or 2 (AP-MS 2) hours to increase the abundance of FAMA and other unstable proteins and crosslinked with formaldehyde. Crosslinking was done by removing excess liquid, submerging the seedlings in crosslinking buffer (25 mM Hepes pH 7.5, 0.5 mM EDTA, 1 mM PMSF, 10 mM MgCl_2_, 75 mM NaCl) containing 0.25 % (AP-MS 1) or 0.125 % (AP-MS 2) formaldehyde and vacuum infiltrating for 15 minutes on ice. Crosslinking was stopped by adding glycine to a final concentration of 120 mM and vacuum infiltrating for another 10 minutes. Seedlings were washed with 800 ml ice cold H_2_O and frozen in liquid nitrogen. Seedlings were treated in two batches per experiment.

#### Affinity purification

For AP, the frozen plant material was ground to a fine powder. Two biological replicates per genotype were used for AP-MS 1 and four replicates of the FAMA-CFP line and two of each of the control lines were used for AP-MS 2. AP-MS 1 was done to test overall performance of the experimental setup before doing a larger scale experiment. Therefore, only a small number of replicates was used. To increase statistical power in AP-MS 2, the replicate number was doubled. Approximately 15 g plant material (from one or two of the treatment batches) per sample were resuspended in 25 ml extraction buffer (25 mM Tris pH 7.5, 10 mM MgCl_2_, 0.5 mM EGTA, 75 mM NaCl, 0.5/1 % Triton-X-100, 1 mM NaF, 0.5 mM Na_3_VO_4_, 15 mM beta glycerophosphate, 1 mM DTT, 1mM PMSF, 1x complete proteasome inhibitor) and incubated on a rotor wheel at 4°C for 30 to 45 minutes to thaw. The extracts were sonicated in 15 ml tubes in an ice bath using Bioruptor UCD-200 (Diagenode) on high setting. For experiment 1, samples were sonicated twice for five minutes with a 30 second on/off interval and a five-minute break on ice. For experiment 2, the sonication cycles were reduced to 2.5 minutes with a 30/60 seconds on/off interval. After sonication, 15 μl Lysonase (Millipore) were added and the extracts were incubated for another 30 minutes on a rotor wheel at 4°C before centrifuging at 12,000 g and 4°C for 10 minutes. The supernatants were filtered through two layers of Miracloth (Millipore) and incubated with 25 μl of GFP-Trap_MA beads (ChromoTek), pre-washed with extraction buffer, for five (AP-MS 1) or three (AP-MS 2) hours at 4°C on a rotor wheel. The beads were then washed six times with extraction buffer containing 100 mM NaCl (1x 15 ml, 5x 1ml), resuspended in 50 μl 2x Leammli buffer and boiled for five minutes at 95°C. Samples were stored at −80°C until further processing.

#### MS sample preparation

Affinity purified proteins were separated by SDS-PAGE using Mini-PROTEAN TGX gels (BioRad) and stained with the Novex colloidal blue staining kit (Invitrogen). About 1 cm long sample areas were excised from the gel, and used for in-gel Tryptic digestion. Peptides were desalted using C18 ZipTips (Millipore). LC-MS/MS was done as described for the PL experiments.

#### MS data analysis

Protein identification and label-free quantification (LFQ) were done for both experiments together. RAW files from LC-MS/MS were searched in MaxQuant (version 1.6.2.6) (Tyanova, Temu, and Cox 2016) using the same settings as described for the PL experiments with one difference: LFQ minimum ratio count was set to 1 instead of 2. The mass spectrometry proteomics data have been deposited to the ProteomeXchange Consortium (http://proteomecentral.proteomexchange.org) via the PRIDE partner repository (Vizcaino et al. 2013) and will be made accessible upon publication.

Filtering and statistical analysis were done with Perseus (version 1.6.2.3) (Tyanova et al. 2016). The ‘proteinGroups.txt’ output file from MaxQuant was imported into Perseus using the LFQ intensities as Main category. The data matrix was filtered to remove proteins marked as ‘only identified by site’, ‘reverse’ and ‘potential contaminant’. LFQ values were log2 transformed and samples from AP-MS 1 and AP-MS 2 were separated for individual analysis. Proteins that were not identified/quantified in at least two (AP-MS 1) or three (AP-MS-2) FAMA-CFP or control samples (low confidence) were removed. Missing values were imputed for statistical analysis using the ‘Replace missing values from normal distribution’ function with the following settings: width = 0.3, down shift = 1.8 and mode = total matrix. Significantly enriched proteins were identified by unpaired 2-sided Students t-tests comparing the FAMA-CFP with the control samples and using the integrated modified permutation-based FDR with 250 randomizations for multiple sample correction. Only proteins that were identified in two (AP-MS 1) or at least three (AP-MS 2) replicates of the FAMA-CFP samples were used for the test. FDR/S0 combinations were chosen to get a minimum number of ‘false negatives’ (statistically enriched proteins in controls). For AP-MS 1, the FDR was 0.2 and S0 was 0.5 and for AP-MS 2, the FDR was 0.01 and S0 was 0.5. Scatterplots with log2-fold changes and −log 10 p-values form t-tests were done in RStudio.

### Primers

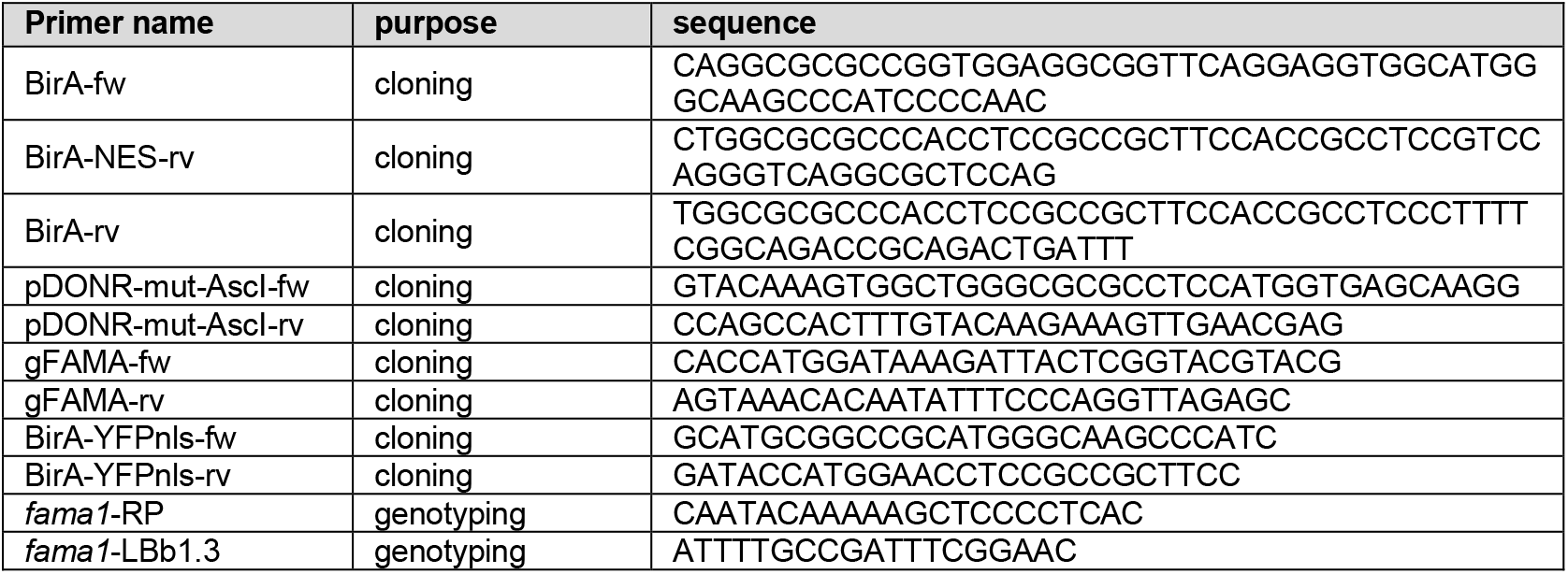

### ‘PL toolbox’ vectors generated for common use

'PL toolbox' vectors generated for common use
| Entry vectors for creating protein of interest-T<sub>ID</sub>/miniT<sub>ID</sub> + YFP/mVenus fusions |  |  |  |
| --- | --- | --- | --- |
| Plasmid name | Gateway sites | application | Addgene ID |
| pENTR_L1-YFP-Turbo-NES-L2 | attL1-attL2 | For N-terminal YFP-T<sub>ID</sub> fusion to non-nuclear proteins | TBD |
| pENTR_L1-YFP-Turbo-L2 | attL1-attL2 | For N-terminal YFP-T<sub>ID</sub> fusion to nuclear proteins | TBD |
| pENTR_L1-YFP-miniTurbo-NES-L2 | attL1-attL2 | For N-terminal YFP-miniT<sub>ID</sub> fusion to non-nuclear proteins | TBD |
| pENTR_L1-YFP-miniTurbo-L2 | attL1-attL2 | For N-terminal YFP-miniT<sub>ID</sub> fusion to nuclear proteins | TBD |
| pDONR-P2R-P3_R2-Turbo-NES-mVenus-STOP-L3 | attR2-attL3 | For C-terminal T<sub>ID</sub>-mVenus fusion to non-nuclear proteins | TBD |
| pDONR-P2R-P3_R2-Turbo-mVenus-STOP-L3 | attR2-attL3 | For C-terminal T<sub>ID</sub>-mVenus fusion to nuclear proteins | TBD |
| pDONR-P2R-P3_R2-miniTurbo-NES-mVenus-STOP-L3 | attR2-attL3 | For C-terminal miniT<sub>ID</sub>-mVenus fusion to non-nuclear proteins | TBD |
| pDONR-P2R-P3_R2-miniTurbo-mVenus-STOP-L3 | attR2-attL3 | For C-terminal miniT<sub>ID</sub>-mVenus fusion to nuclear proteins | TBD |
| Entry vectors for expressing BirA*/T<sub>ID</sub>/miniT<sub>ID</sub> + YFP under a promoter of choice |  |  |  |
| Plasmid name | Gateway sites | application | Addgene ID |

| pENTR_L1-BirA(R118G)-NES-YFP-STOP-L2 | attL1-attL2 | For expressing cytosolic BirA* under a promoter of choice | *TBD* |
| --- | --- | --- | --- |
| pENTR_L1-Turbo-NES-YFP-STOP-L2 | attL1-attL2 | For expressing cytosolic T<sub>ID</sub> under a promoter of choice | *TBD* |
| pENTR_L1-miniTurbo-NES-YFP-STOP-L2 | attL1-attL2 | For expressing cytosolic miniT<sub>ID</sub> under a promoter of choice | *TBD* |
| pENTR_L1-BirA(R118G)-YFP-NLS-STOP-L2 | attL1-attL2 | For expressing nuclear BirA* under a promoter of choice | *TBD* |
| pENTR_L1-Turbo-YFP-NLS-STOP-L2 | attL1-attL2 | For expressing nuclear T<sub>ID</sub> under a promoter of choice | *TBD* |
| pENTR_L1-miniTurbo-YFP-NLS-STOP-L2 | attL1-attL2 | For expressing nuclear miniT<sub>ID</sub> under a promoter of choice | *TBD* |
| **Binary plant transformation vectors for ubiquitous expression of BirA\*/T<sub>ID</sub>/miniT<sub>ID</sub> + YFP/mVenus** |  |  |  |
| Plasmid name | Resistance in plants | application | Addgene ID |
| R4pGWB601_UBQ10p::BirA(R118G)-NES-YFP | BASTA | For expressing cytosolic BirA* under the UBQ10 promoter | *TBD* |
| R4pGWB601_UBQ10p::BirA(R118G)-YFP-NLS | BASTA | For expressing nuclear BirA* under the UBQ10 promoter | *TBD* |
| R4pGWB601_UBQ10p::Turbo-NES-YFP | BASTA | For expressing cytosolic T<sub>ID</sub> under the UBQ10 promoter | *TBD* |
| R4pGWB601_UBQ10p::Turbo-YFP-NLS | BASTA | For expressing nuclear T<sub>ID</sub> under the UBQ10 promoter | *TBD* |
| R4pGWB601_UBQ10p::miniTurbo-NES-YFP | BASTA | For expressing cytosolic miniT<sub>ID</sub> under the UBQ10 promoter | *TBD* |
| R4pGWB601_UBQ10p::miniTurbo-YFP-NLS | BASTA | For expressing nuclear miniT<sub>ID</sub> under the UBQ10 promoter | *TBD* |

## Supporting information

Supplemental Figures linked to six main text figures

## Acknowledgements

We thank Annika Weimer for suggestions for proximity labeling construct design and testing of biotin ligase activity and other previous and current members of the Bergmann lab for creating tools that enabled this work and for critical discussion. Work in this publication has been funded with support from the following sources: AM was supported in part by a Schrödinger fellowship from the Austrian Science Fund FWF (grant number J4019-B29) and TCB was supported by Dow Graduate Research and Lester Wolfe Fellowships. DCB is an investigator of the Howard Hughes Medical institute. Further support came from the NIH grant R01-CA186568 (to AYT) and from the Carnegie endowment fund to the Carnegie mass spectrometry facility.

## Author Contributions

AM designed and performed all experiments, designed figures and wrote the manuscript. SLX generated and did the initial analysis of mass spectrometry data, TCB and AYT provided reagents and made comments on the manuscript. DCB designed experiments, wrote the MS and provided funding.

## Supplementary files

Supplementary File 1: non-cropped immunoblots

## Recurring abbreviations

AP: affinity purification
AP-MS: affinity purification-mass spectrometry
bHLH: basic helix-loop-helix
BirA*: promiscuous E. coli biotin ligase
FDR: false discovery rate
GC: guard cell
GO: gene ontology
HAT: histone acetyl transferase
HDAC: histone deacetylase
LC-MS/MS: liquid chromatography coupled to tandem mass spectrometry
MS: mass spectrometry
miniT_ID_: miniTurboID
NES: nuclear export signal
NLS: nuclear localization signal
PCA: principal component analysis
PL: proximity labeling
POI: protein of interest
SA: streptavidin
TF: transcription factor
T_ID_: TurboID
WT: wild-type
Y2H: yeast-2-hybrid

