## Supplemental Figures linked to six main text figures for "Proximity labeling of protein complexes and cell type-specific organellar proteomes in Arabidopsis enabled by TurboID"

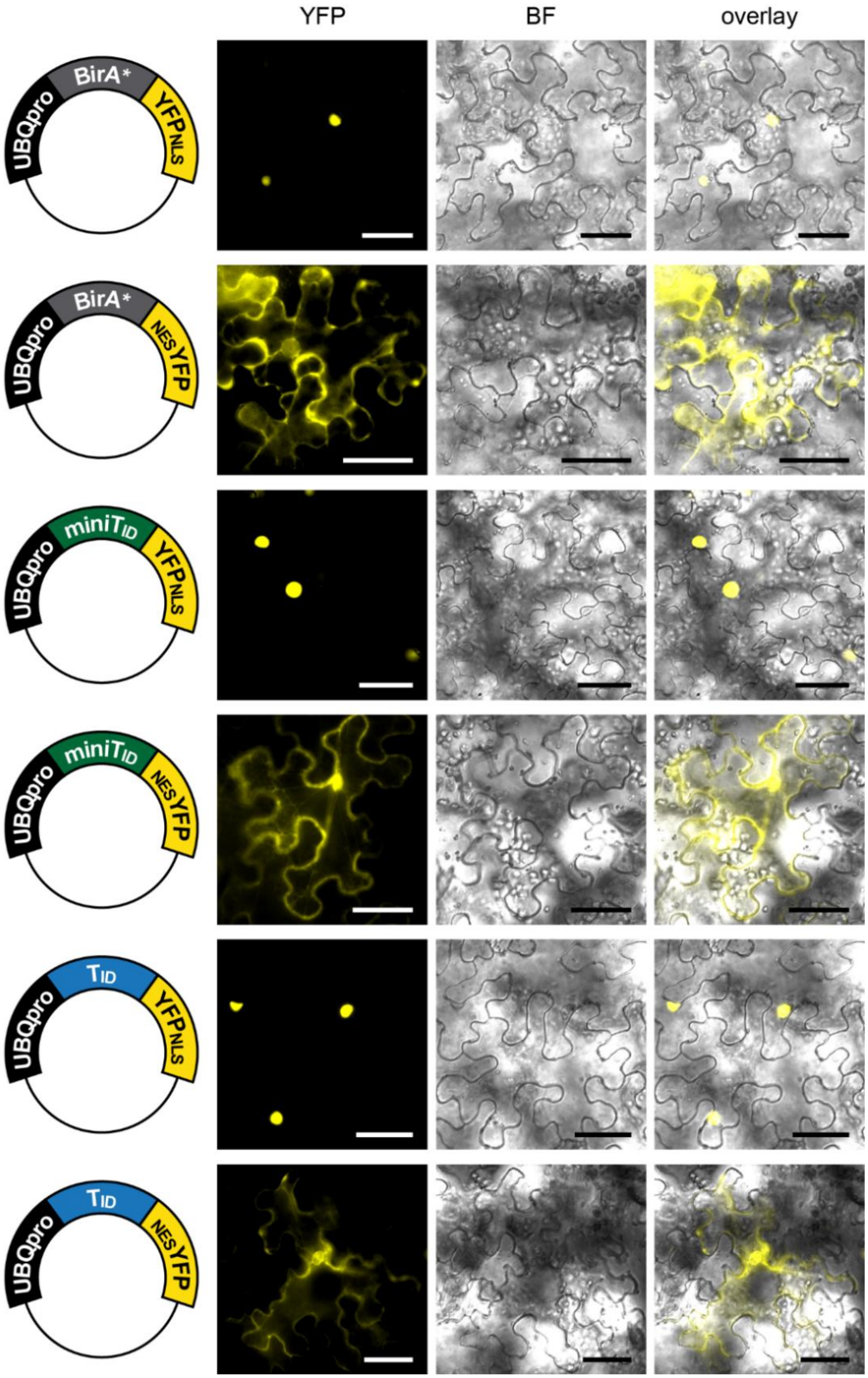


**Figure 1 – figure supplement 1: Subcellular localization of biotin ligase constructs in transiently transformed *N. benthamiana* leaves**

Epifluorescence images of *N. benthamiana* leaves transformed with UBQ10pro::BirA-YFP$_{NLS}$ or UBQ10pro::BirA-$_{NES}$YFP expression vectors (BirA = BirA*, miniT$_{ID}$ or T$_{ID}$) two days after transformation. Shown are the YFP and brightfield (BF) channels and an overlay. Scale bar = 50 µm. Constructs are indicated on the left. Nuclear targeted BirA-YFP (NLS) constructs were largely nuclear and cytosol targeted BirA-YFP versions (NES) constructs largely cytosolic.

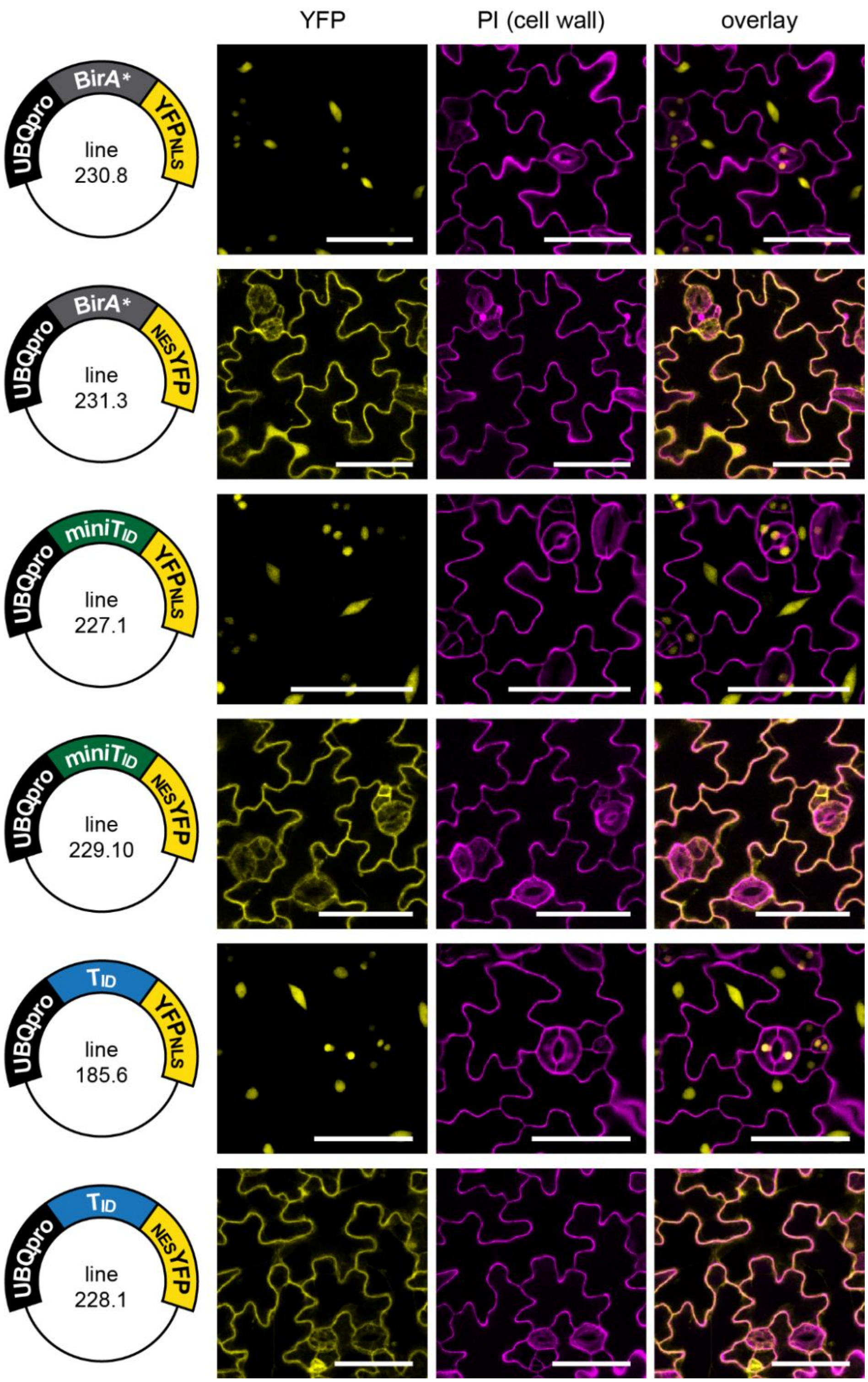


**Figure 1 – figure supplement 2: Subcellular localization of biotin ligase constructs in stable Arabidopsis lines**

Confocal microscopy images of the cotyledon epidermis of five day old Arabidopsis seedlings transformed with UBQ10pro::BirA-YFP$_{NLS}$ or UBQ10pro::BirA-$_{NES}$YFP expression vectors (BirA = BirA*, miniT$_{ID}$ or T$_{ID}$). Shown are the YFP channel (yellow), the cell walls stained with propidium iodide (purple) and an overlay. Scale bar = 50 µm. Constructs and lines used for images are indicated on the left. Nuclear targeted BirA-YFP (NLS) constructs were mostly nuclear, and cytosol targeted (NES) constructs were excluded from the nucleus.

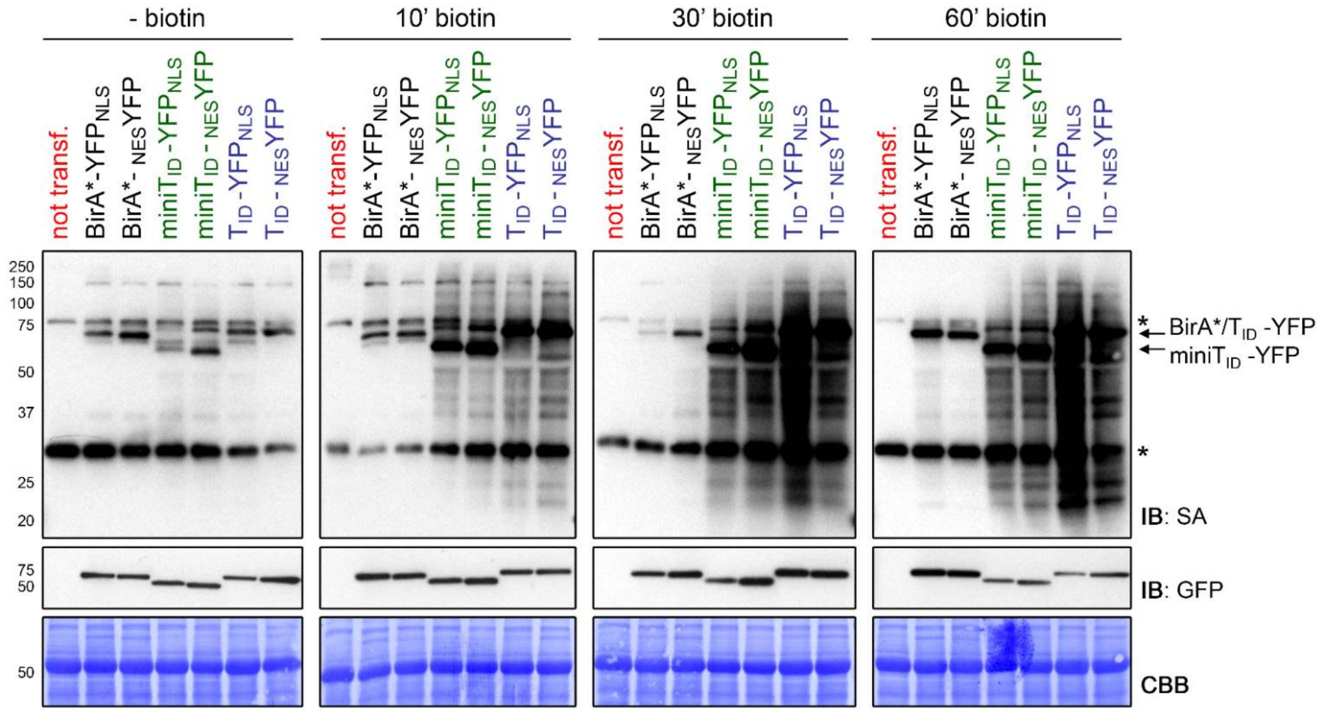


**Figure 1 – figure supplement 3: $T_{ID}$ and mini$T_{ID}$ are highly active in the cytosol and nucleus of transiently transformed *N. benthamiana* leaves**

*N. benthamiana* leaves were transformed with UBQ10pro::BirA-$YFP_{NLS}$ or UBQ10pro::BirA-$_{NES}$YFP expression vectors (BirA = BirA*, mini$T_{ID}$ or $T_{ID}$). Two days after infiltration, leaf discs expressing either of the six constructs or non-transformed leaves (not transf.) were briefly vacuum infiltrated with 50 µM biotin and incubated for 10, 30 or 60 minutes. Untreated samples were used as controls to visualize the background activity of the biotin ligases with endogenous biotin. Each sample is a pool of 2 leaf discs. Activity and expression of the BirA-YFP variants were analyzed by immunoblots (IB) with streptavidin-HRP (SA) and anti-GFP antibodies. Coomassie Brilliant Blue-stained membranes (CBB) are shown as loading controls. Asterisks mark the positions of naturally biotinylated proteins.

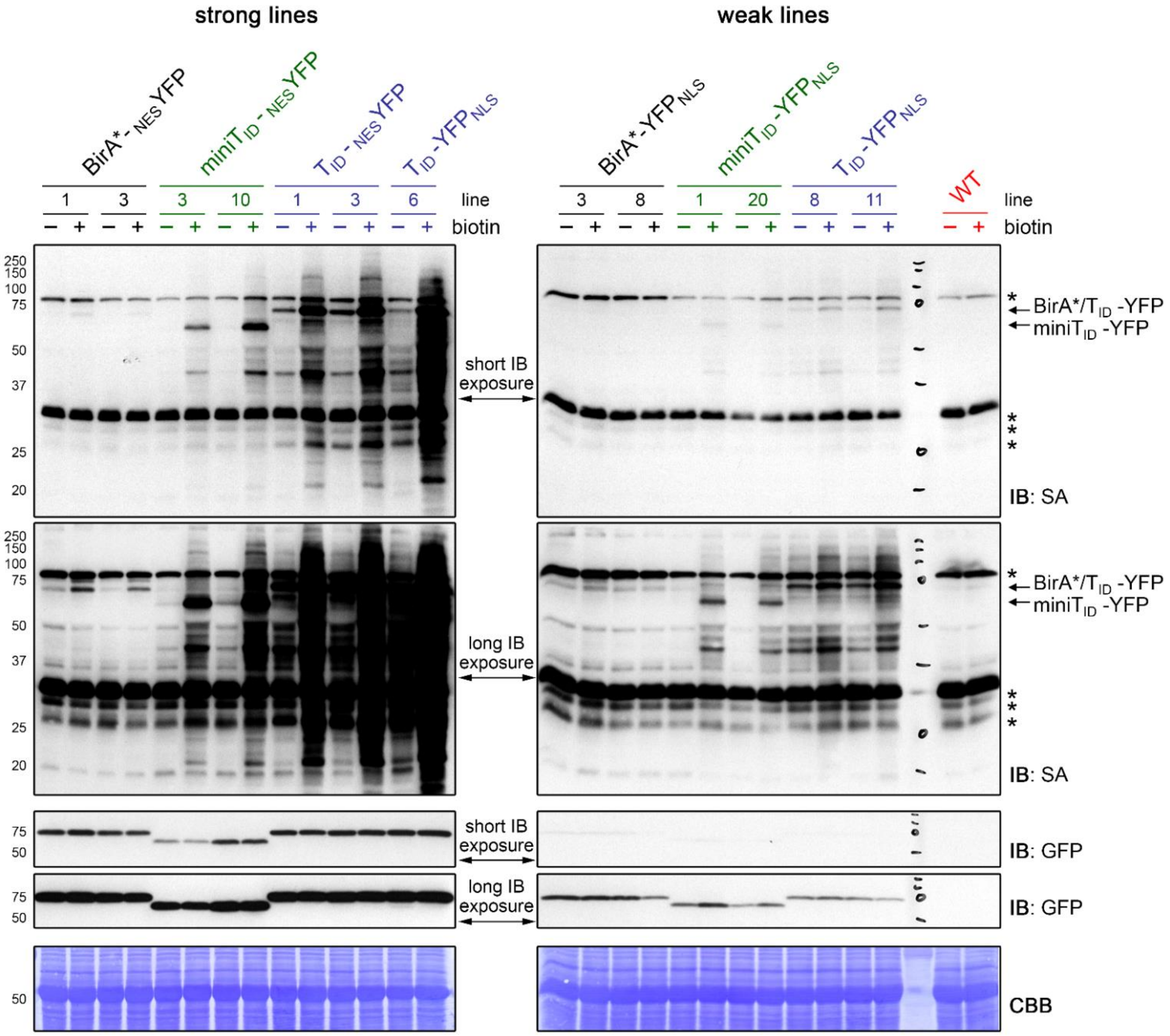


**Figure 1 – figure supplement 4: $T_{ID}$ is more active than mini$T_{ID}$, but also produces more background labeling in Arabidopsis**

Five day old Arabidopsis seedlings expressing UBQ10pro::BirA-$YFP_{NLS}$ or UBQ10pro::BirA-$_{NES}$YFP constructs (BirA = BirA*, mini$T_{ID}$ or $T_{ID}$) were submerged in 250 µM biotin, briefly vacuum infiltrated and incubated at room temperature for one hour (+). Non-treated seedlings were used as controls (-) to visualize the background activity of the biotin ligases with endogenous biotin. Activity and expression of the BirA-YFP variants were analyzed by immunoblots (IB) with streptavidin-HRP (SA) and anti-GFP antibodies. Coomassie Brilliant Blue-stained membranes (CBB) are shown as loading controls. Asterisks mark the positions of naturally biotinylated proteins. Each sample is a pool of ~ 30 seedlings. At least two lines were tested per construct; strong cytosolic lines and a strong nuclear $T_{ID}$ line are on the left, weak nuclear lines and WT are on the right. Blots were made and exposed in parallel. Different exposure times are shown to enable comparison of weak and strong lines. Short exposure times were five seconds, long exposures times were 30 seconds (SA blot) and 70 seconds (anti-GFP blot). Mini$T_{ID}$ is less active than $T_{ID}$, but has less background labeling with endogenous biotin, especially in low-expressing lines. WT samples and samples from lines 3, 10 and 3 of the BirA*-$_{NES}$YFP, mini$T_{ID}$-$_{NES}$YFP and $T_{ID}$-$_{NES}$YFP plants were also used for the blots shown in **Figure 1C**.

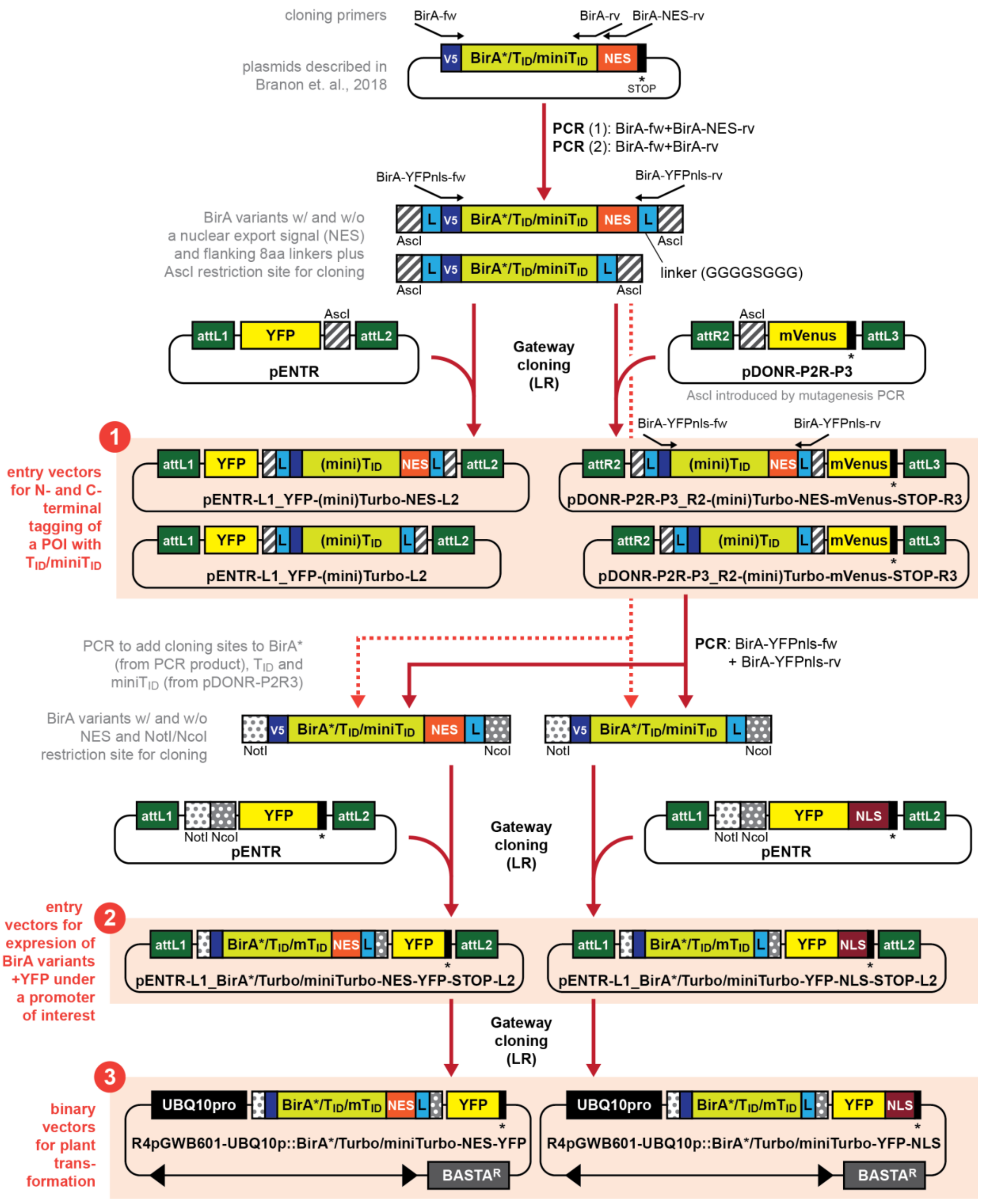


**Figure 1 – figure supplement 5: Generating a toolbox of gateway-compatible vectors for PL in plants**

Schematic overview over cloning steps involved in the generation of the PL toolbox and over the available vectors (highlighted as orange boxes). Analogous constructs for different biotin ligase variants are summarized (indicated by BirA*/$T_{ID}$/m$T_{ID}$ or $T_{ID}$/m$T_{ID}$). Primers and restriction sites used for PCR and restriction cloning are indicated on the top or bottom of the constructs. V5 = V5 tag, L = 8 amino acid linker (GGGGSGGG), NES = nuclear export signal, NLS = nuclear localization signal, POI = protein of interest. **Available plasmids** include: ❶ 8 entry vectors for N- and C-terminal tagging of proteins of interest with $T_{ID}$ or mini$T_{ID}$ plus a fluorophore, ❷ 6 entry vectors for expressing any of the three BirA variants under a promoter of interest with a NES or NLS and a fluorophore, and ❸ 6 binary plant transformation vectors for expressing any of the three BirA variants with an NES or NLS under the UBQ10 promoter. A detailed description and primer sequences are included in the materials and methods section. A list of available plasmids with Addgene catalogue number can also be found in the methods section.

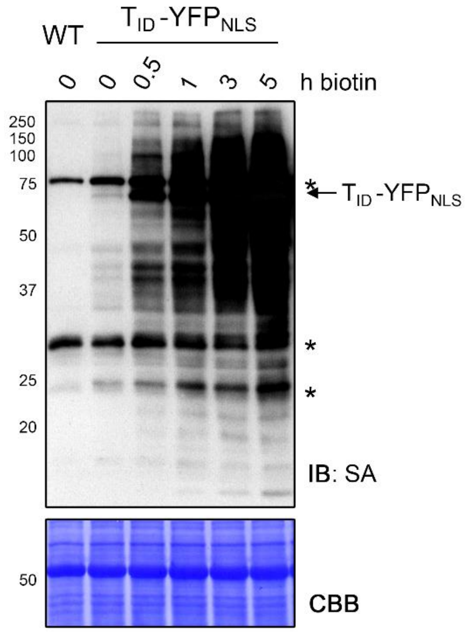


**Figure 2 – figure supplement 1: Biotinylation by $T_{ID}$ in Arabidopsis increases over time**

Labeling time course with $T_{ID}$ in four day old Arabidopsis seedlings expressing the UBQ10pro::$T_{ID}$-$YFP_{NLS}$ construct ($T_{ID}$-$YFP_{NLS}$). Seedlings were submerged in 50 µM biotin, briefly vacuum infiltrated and incubated for the indicated time at room temperature. A control sample and a wild-type (WT) sample were taken before biotin treatment. Activity of $T_{ID}$ over time was analyzed by immunoblot (IB) with streptavidin-HRP (SA). The Coomassie Brilliant Blue-stained membrane (CBB) is shown as loading control. Asterisks mark the positions of naturally biotinylated proteins. Each sample is a pool of seedlings. The increase of labeling between three and five hours of biotin treatment demonstrates that labeling of nuclear proteins is not saturated after three hours.

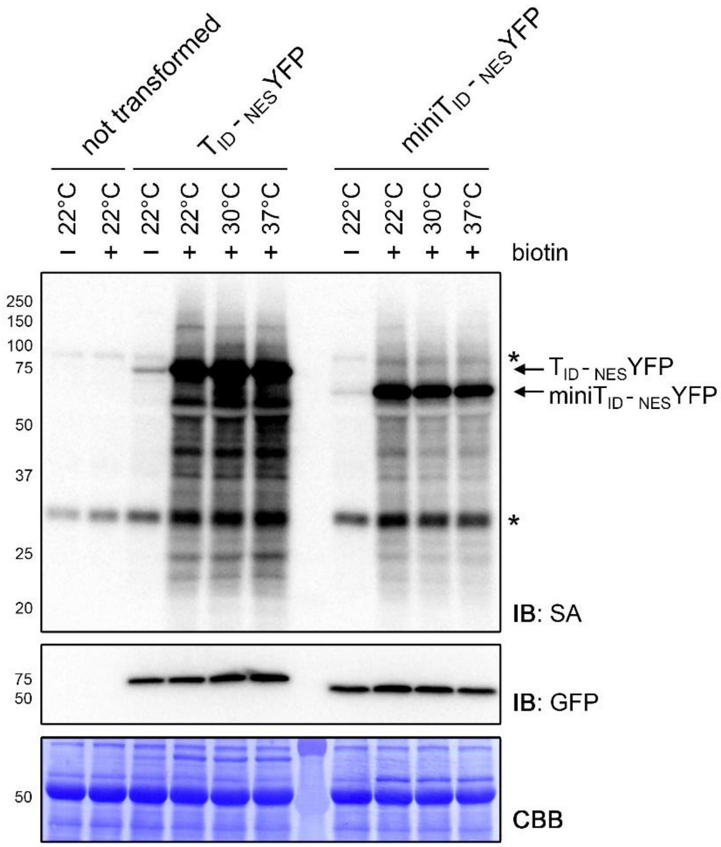


**Figure 2 – figure supplement 2: $T_{ID}$ and mini$T_{ID}$ are active from 22°C to 37°C in *N. benthamiana***

Streptavidin-HRP (SA) and anti-GFP immunoblots (IB) showing the activity and expression of $T_{ID}$ and mini$T_{ID}$ at room temperature (22°C), 30°C and 37°C. *N. benthamiana* leaves were transformed with UBQ10pro::$T_{ID}$-$_{NES}$YFP ($T_{ID}$-$_{NES}$YFP) or UBQ10pro::mini$T_{ID}$-$_{NES}$YFP (mini$T_{ID}$-$_{NES}$YFP) expression vectors. Two days after infiltration, leaf discs expressing the $T_{ID}$ and mini$T_{ID}$ constructs and non-transformed leaves were briefly vacuum infiltrated with $H_2O$ (-) or 250 µM biotin (+) and incubated at the indicated temperature for one hour. Each sample is a pool of tree leaf discs. The Coomassie Brilliant Blue-stained membrane (CBB) is shown as loading control. Asterisks mark the positions of naturally biotinylated proteins.

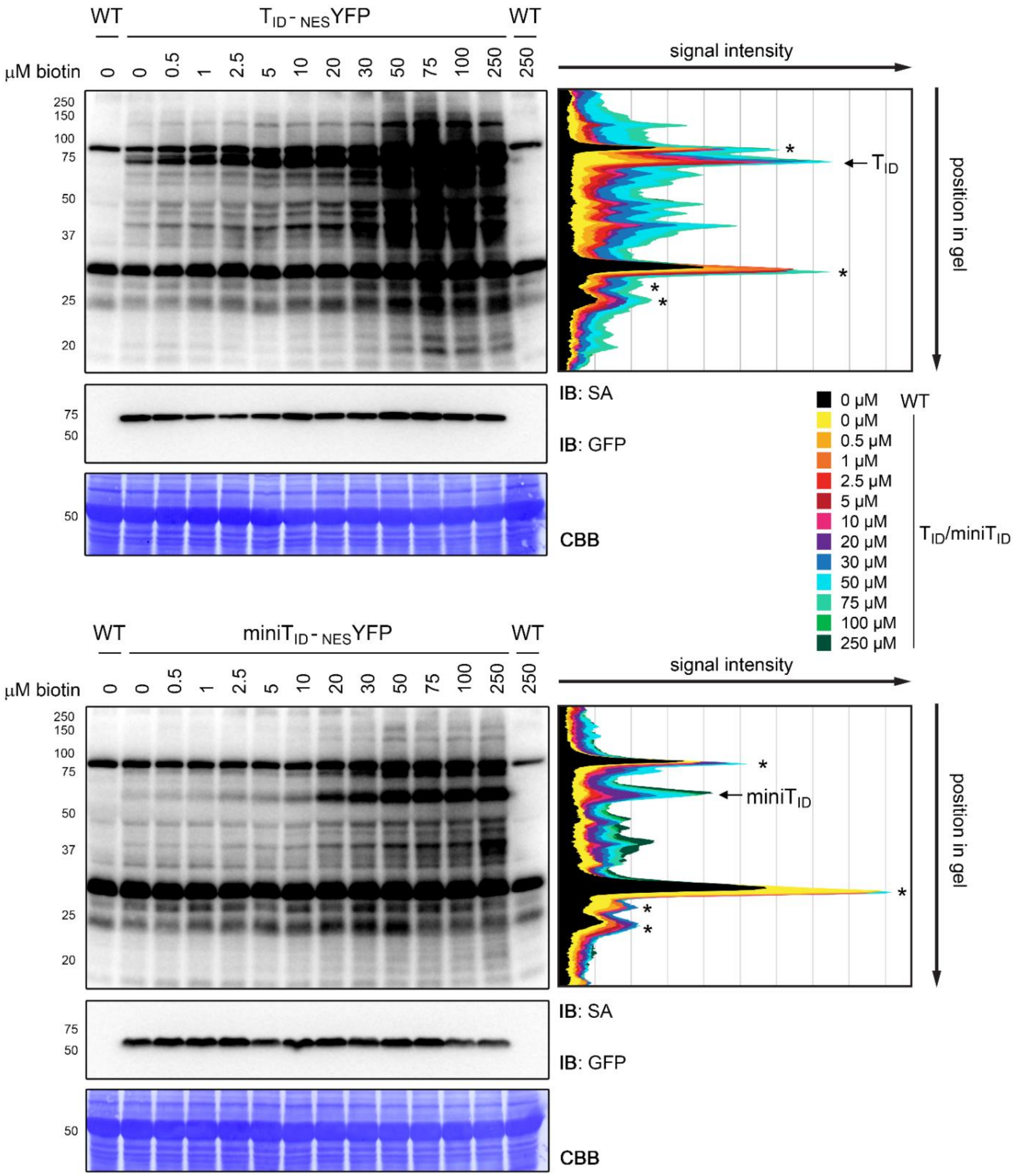


**Figure 2 – figure supplement 3: Quantification of $T_{ID}$ and miniT$_{ID}$ activity in Arabidopsis at different biotin concentrations**

Quantification of the streptavidin-HRP (SA) immunoblots (IB) shown in **Figure 2C**. Five day old seedlings transformed with UBQ10pro::$T_{ID}$-$_{NES}$YFP ($T_{ID}$-$_{NES}$YFP) and UBQ10pro::miniT$_{ID}$-$_{NES}$YFP (miniT$_{ID}$-$_{NES}$YFP) were submerged in 0.5 to 250 µM biotin and incubated for one hour at room temperature. A control sample was taken before treatment (0 µM). The anti-GFP blot shows expression of the YFP fusion proteins. Quantification was done in FIJI (ImageJ) by drawing five parallel vertical lines through each of the lanes at different positions and reading out the signal intensity along the line. The graphs on the right show the average signal intensity of the five lines along the length of the gel. The peaks associated with $T_{ID}$-YFP and miniT$_{ID}$-YFP are indicated and the positions of naturally biotinylated proteins are marked by asterisks.

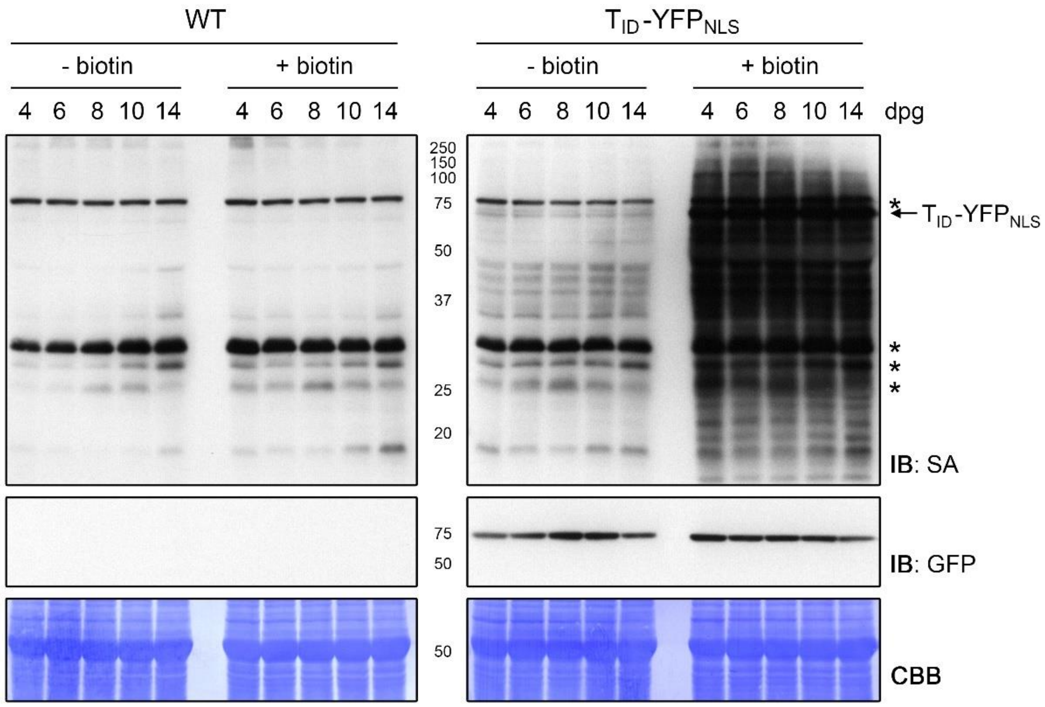


**Figure 3 – figure supplement 1: Activity and background labeling of $T_{ID}$ are similar in seedlings ranging from 4 to 14 days of age**

Plate-grown Arabidopsis wild-type (WT) and UBQ10pro::$T_{ID}$-$YFP_{NLS}$ ($T_{ID}$-$YFP_{NLS}$) seedlings of the indicated age (dpg = days post germination) were submerged in a 250 µM biotin solution, briefly vacuum infiltrated and incubated for one hour at room temperature (+ biotin). Non-treated control samples (- biotin) were taken at the same time. Each sample is a pool of seedlings. Activity and expression of $T_{ID}$-$YFP_{NLS}$ were analyzed by immunoblots (IB) with streptavidin-HRP (SA) and anti-GFP antibodies. Coomassie Brilliant Blue-stained membranes (CBB) are shown as a loading controls. Asterisks mark the positions of naturally biotinylated proteins.

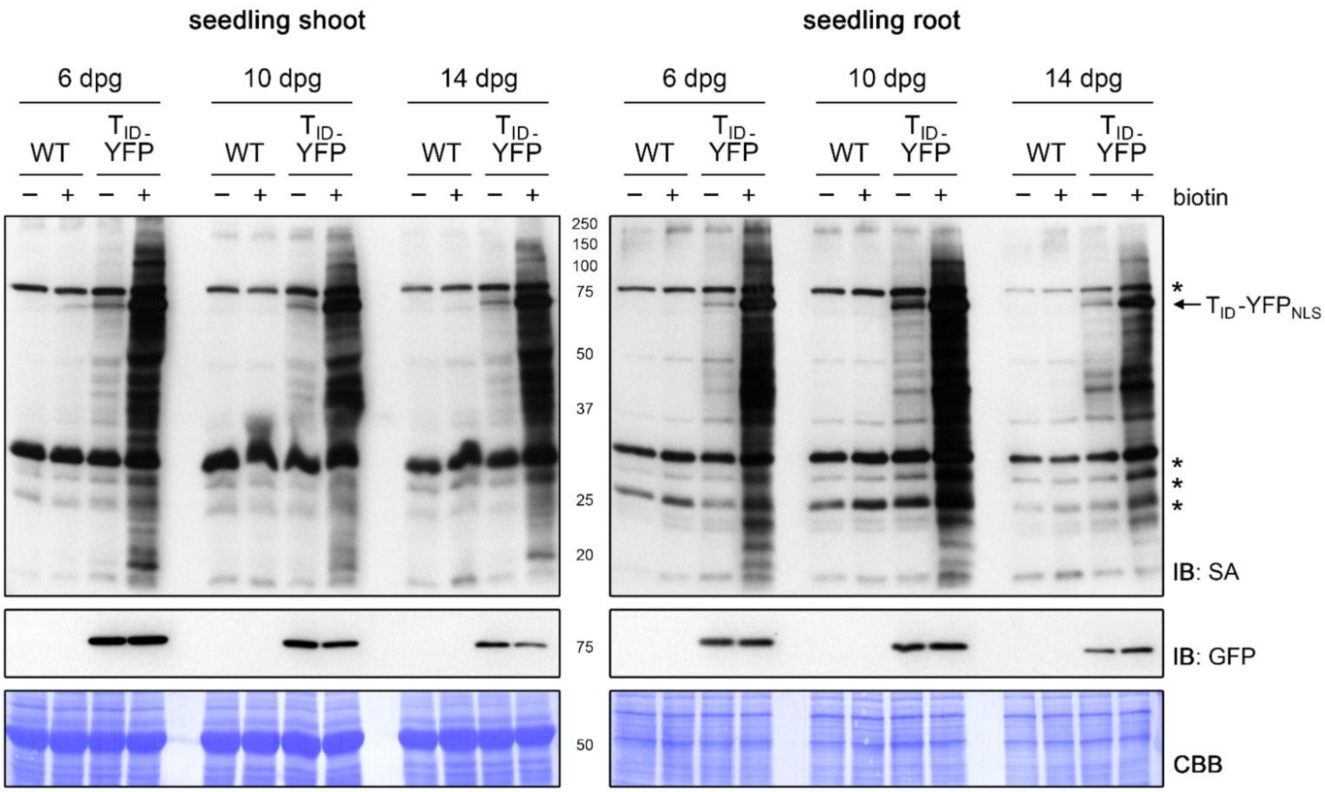


**Figure 3 – figure supplement 2: Activity and background labeling of $T_{ID}$ are similar in roots and shoots of 6 to 14 day old seedlings**

Six, 10 and 14 days post germination (dpg), plate-grown Arabidopsis wild-type (WT) and UBQ10pro::$T_{ID}$-$YFP_{NLS}$ ($T_{ID}$-$YFP_{NLS}$) seedlings were divided into a shoot and root section. Shoots and roots were submerged separately in a 250 µM biotin solution, briefly vacuum infiltrated and incubated for one hour at room temperature (+). Non-treated control samples (-) were taken at the same time. Each sample is a pool of seedlings. Activity and expression of $T_{ID}$-$YFP_{NLS}$ were analyzed by immunoblots (IB) with streptavidin-HRP (SA) and anti-GFP antibodies, respectively. Coomassie Brilliant Blue-stained membranes (CBB) are shown as a loading control. Asterisks mark the positions of naturally biotinylated proteins.

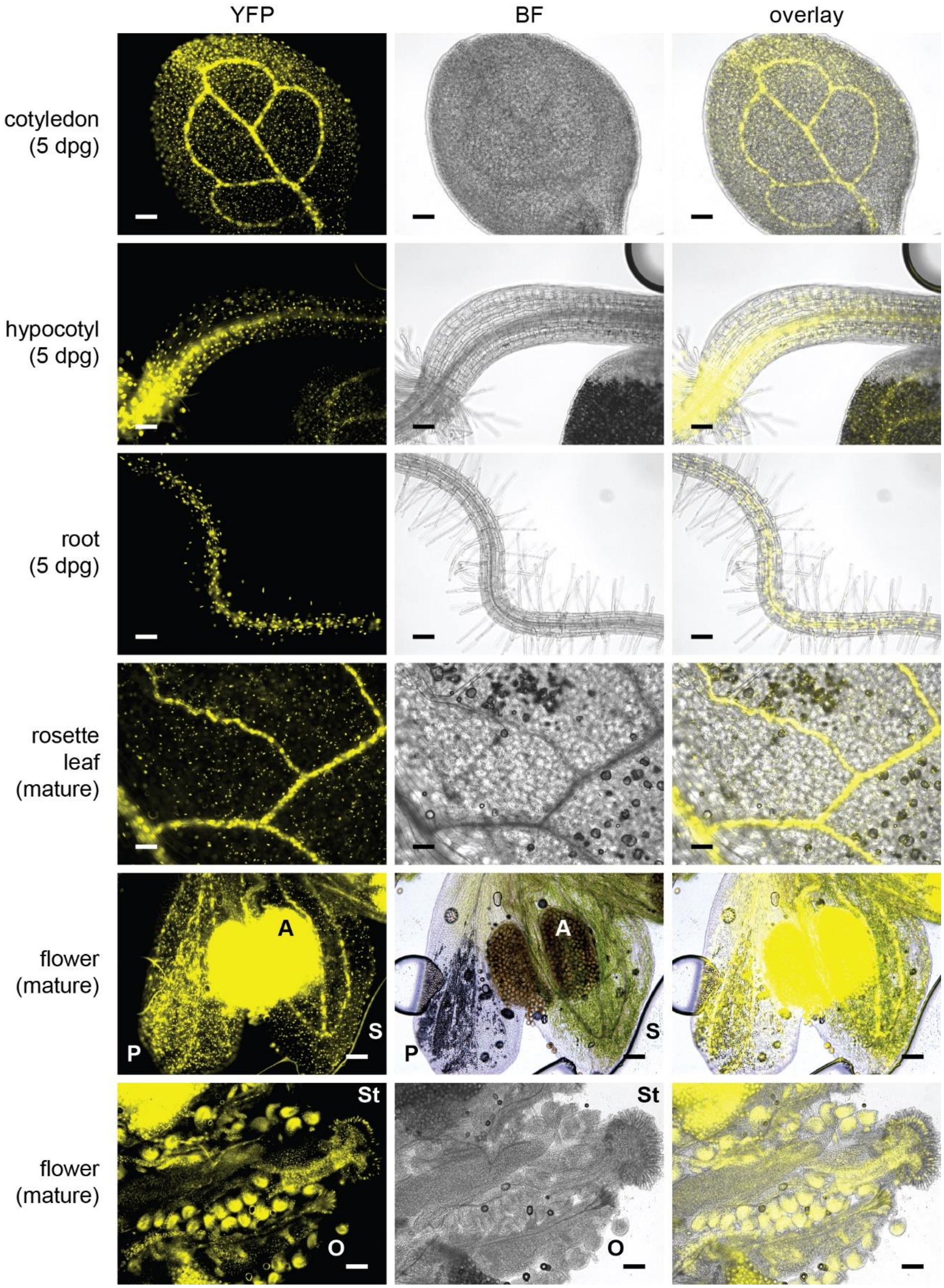


**Figure 3 – figure supplement 3: UBQ10pro::$T_{ID}$-$YFP_{NLS}$ is expressed throughout the whole plant**

Epifluorescence microscopy images of plate-grown Arabidopsis seedlings and soil-grown mature tissues from the strong $T_{ID}$-$YFP_{NLS}$ line used in **Figures 2** and **3** and the PL experiments in **Figures 4** to **6**. Shown are the YFP and brightfield (BF) channel and an overlay. Scale bar = 100 µm. Abbreviations: P, petal; S, sepal; A, anther; St, stigma; O, ovary. The overlay images of the 5 day old cotyledon and root and of mature flowers are also used in **Figure 3**.

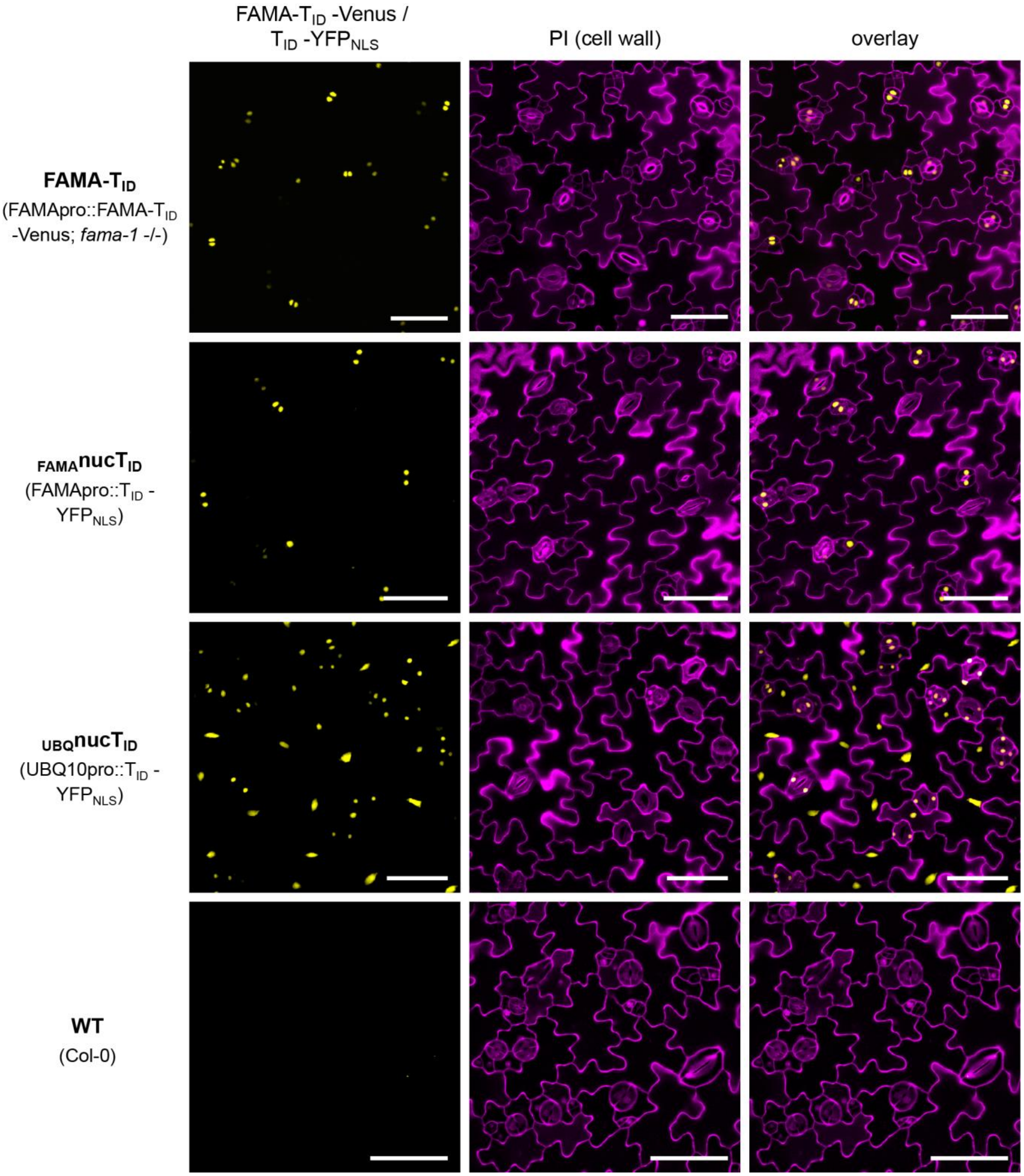


**Figure 4 – figure supplement 1: Expression of the $T_{ID}$ constructs in lines used for the 'FAMA interactome' and 'nuclear proteome' PL experiments**

Confocal microscopy images of the cotyledon epidermis of five day old Arabidopsis seedlings. Shown are the YFP channel (yellow), the cell walls stained with propidium iodide (purple) and an overlay. Scale bar = 50 µm. Excerpts of the overlay images were used in **Figure 4A**. FAMA-$T_{ID}$-Venus and $T_{ID}$-$YFP_{NLS}$ controlled by the FAMA promoter are expressed in developing guard cells (GCs). Expression is highest in dividing and young GCs and decreases as GCs mature. We did not see expression in developing myrosin cells (Li and Sack 2014), but cannot rule out expression at growth stages that were not used in our experiments. $T_{ID}$-$YFP_{NLS}$ controlled by the UBQ10 promoter is expressed ubiquitously (see also Figure 3 – figure supplement 3).

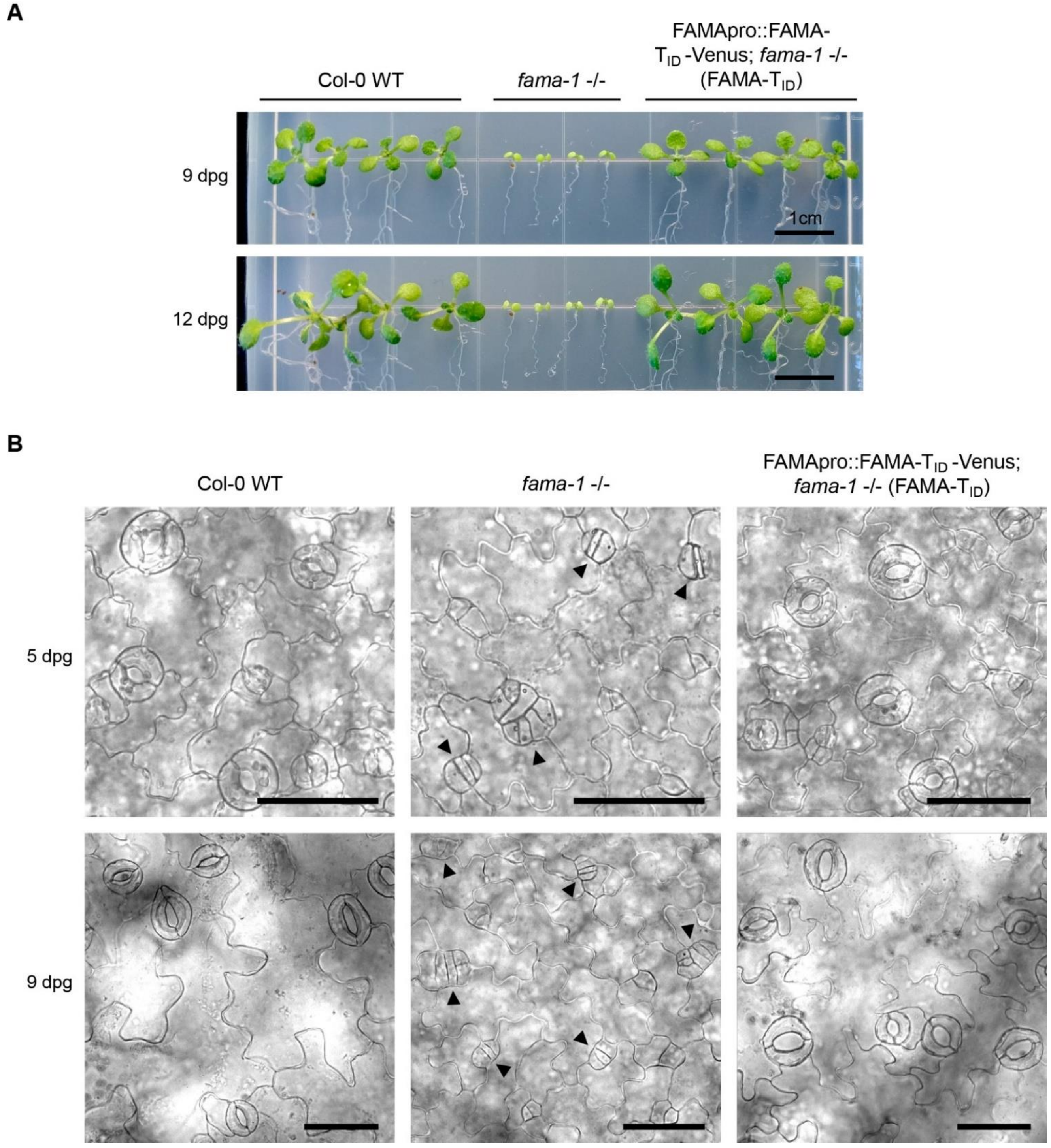


**Figure 4 – figure supplement 2: The FAMApro::FAMA-T$_{ID}$-Venus construct rescues the *fama-1* mutant phenotype**

**(A) FAMA-T$_{ID}$-Venus rescues the seedling-lethal growth phenotype of *fama-1*.** Col-0 wild-type (WT), *fama-1 -/-* and the FAMA-T$_{ID}$ rescue line (FAMApro::FAMA-T$_{ID}$-Venus in *fama-1 -/-* background) were grown on ½ MS plates supplemented with 0.5% sucrose. Pictures of the seedlings were taken after 9 and 12 days. **(B) FAMA-T$_{ID}$-Venus rescues the stomatal development phenotype of *fama-1*.** Brightfield images of the cotyledon epidermis of seedlings five and 9 days post germination (dpg). Scale bar = 50 µm. In the *fama-1* mutant, guard cell precursors continue dividing and fail to differentiate into stomata with a central pore, resulting in clusters of guard cell precursors (black arrowheads).

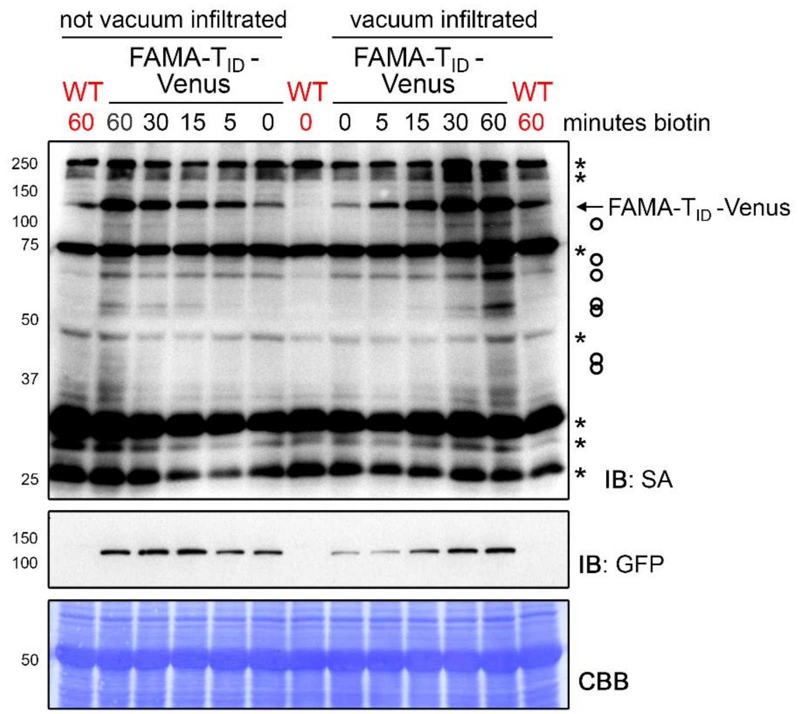


**Figure 4 – figure supplement 3: FAMA-$T_{ID}$ self-labeling increases within five minutes of biotin treatment regardless of vacuum infiltration**

Time course to determine time points for the PL experiments with FAMA-$T_{ID}$. Five day old wild-type (WT) and FAMApro::FAMA-$T_{ID}$-Venus (FAMA-$T_{ID}$-Venus) seedlings were submerged in 250 µM biotin, briefly vacuum infiltrated or not and incubated for the indicated time at room temperature. Control samples were taken before treatment (0 minutes). Activity and expression of $T_{ID}$ were analyzed by immunoblots (IB) with streptavidin-HRP (SA) and anti-GFP antibodies. The Coomassie Brilliant Blue-stained membrane (CBB) is shown as loading control. Asterisks and circles mark the positions of naturally biotinylated proteins and putative FAMA-$T_{ID}$ targets, respectively. Each sample is a pool of ~ 20 seedlings.

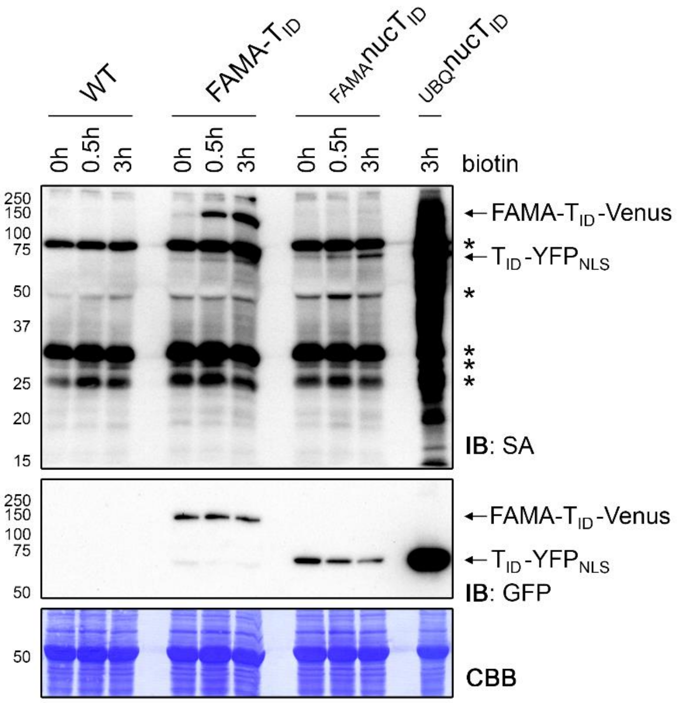


**Figure 4 – figure supplement 4: Confirming successful labeling of proteins in the PL experiment**

For the PL experiments shown in **Figures 4-6**, five day old wild-type (WT), FAMA-$T_{ID}$, $_{FAMA}$nuc$T_{ID}$ and $_{UBQ}$nuc$T_{ID}$ seedlings were submerged in a 50 μM biotin solution for 0, 0.5 and 3 hours and washed repeatedly with ice cold water. Activity and expression of the $T_{ID}$ constructs at the tree time points were analyzed by immunoblots (IB) with streptavidin-HRP (SA) and anti-GFP antibodies. The Coomassie Brilliant Blue-stained membrane (CBB) is shown as loading control. Asterisks mark the positions of naturally biotinylated proteins. See Figure 4 – figure supplement 5 for affinity purification of the biotinylated proteins.

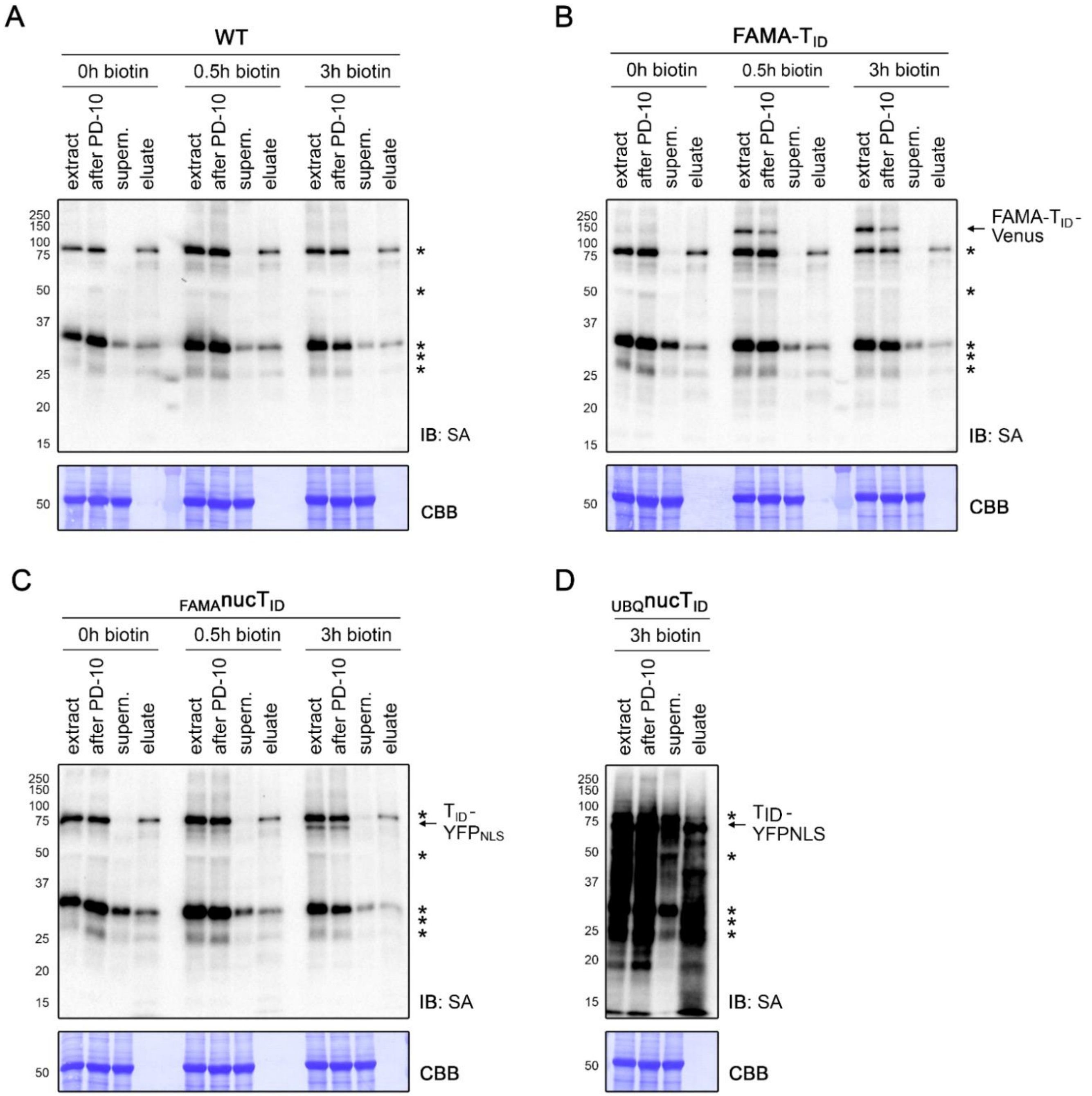


**Figure 4 - figure supplement 5: Affinity purification of biotinylated proteins in the PL experiment**

Five day old wild-type (WT), FAMA-$T_{ID}$, ${}_{FAMA}nucT_{ID}$ and ${}_{UBQ}nucT_{ID}$ seedlings were treated with biotin for 0, 0.5, and 3 hours (see Figure 4 – figure supplement 4) and used for affinity purification (AP) of biotinylated proteins. Samples were taken at different steps of the AP procedure and used for immunoblots (IB) with streptavidin-HRP (SA) to confirm successful biotin depletion and binding of biotinylated proteins to the beads **(A-D)**. One of three biological replicates is shown for each sample. The positions of the $T_{ID}$ fusion proteins are indicated on the right and naturally biotinylated proteins are marked by asterisks. Protein extracts before (extract) and after biotin depletion (after PD-10) confirm that no significant sample loss occurred at this step. The protein extracts after AP (supern.) were depleted of biotinylated proteins, indicating binding of these proteins to the beads. Small aliquots of the beads were used for elution of bound proteins. In most cases, only the naturally biotinylated proteins could be eluted, presumably because $T_{ID}$ targets are biotinylated on multiple sites and bind the beads more strongly.

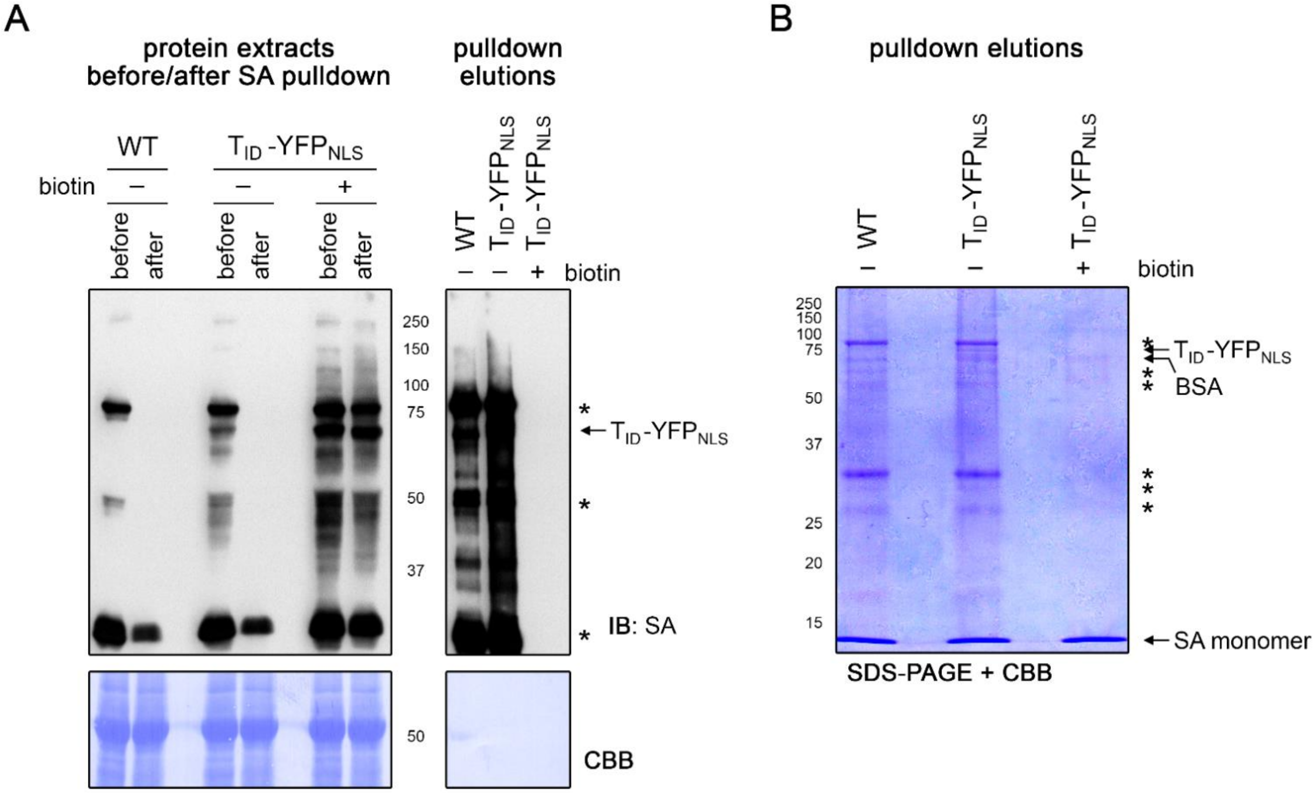


**Figure 4 – figure supplement 6: Free biotin from biotin treatment out-competes biotinylated proteins for streptavidin bead binding**

Four day old wild-type (WT) and UBQ10pro::$T_{ID}$-$YFP_{NLS}$ ($T_{ID}$-$YFP_{NLS}$) seedlings were submerged in $H_2O$ (-) or 50 µM biotin for one hour and used for affinity purification of biotinylated proteins with streptavidin (SA) beads. Protein extracts before and after overnight incubation with the beads and proteins eluted from the beads were used for immunoblotting (IB) with streptavidin-HRP (SA) **(A)** and for SDS-PAGE **(B)** to analyze binding of biotinylated proteins to the beads. Biotin treatment led to an increase of labeling in the $T_{ID}$-$YFP_{NLS}$ line, but also inhibited AP of biotinylated proteins as can be seen both in the protein extracts after incubation with the beads and in the eluates. Coomassie Brilliant Blue-stained membranes (CBB) are shown as loading controls for the IB. Asterisks mark the positions of naturally biotinylated proteins. The positions of $T_{ID}$-$YFP_{NLS}$, as well as of BSA (beads used in this experiment are blocked with BSA) and SA monomers that are eluted from the beads, are indicated.

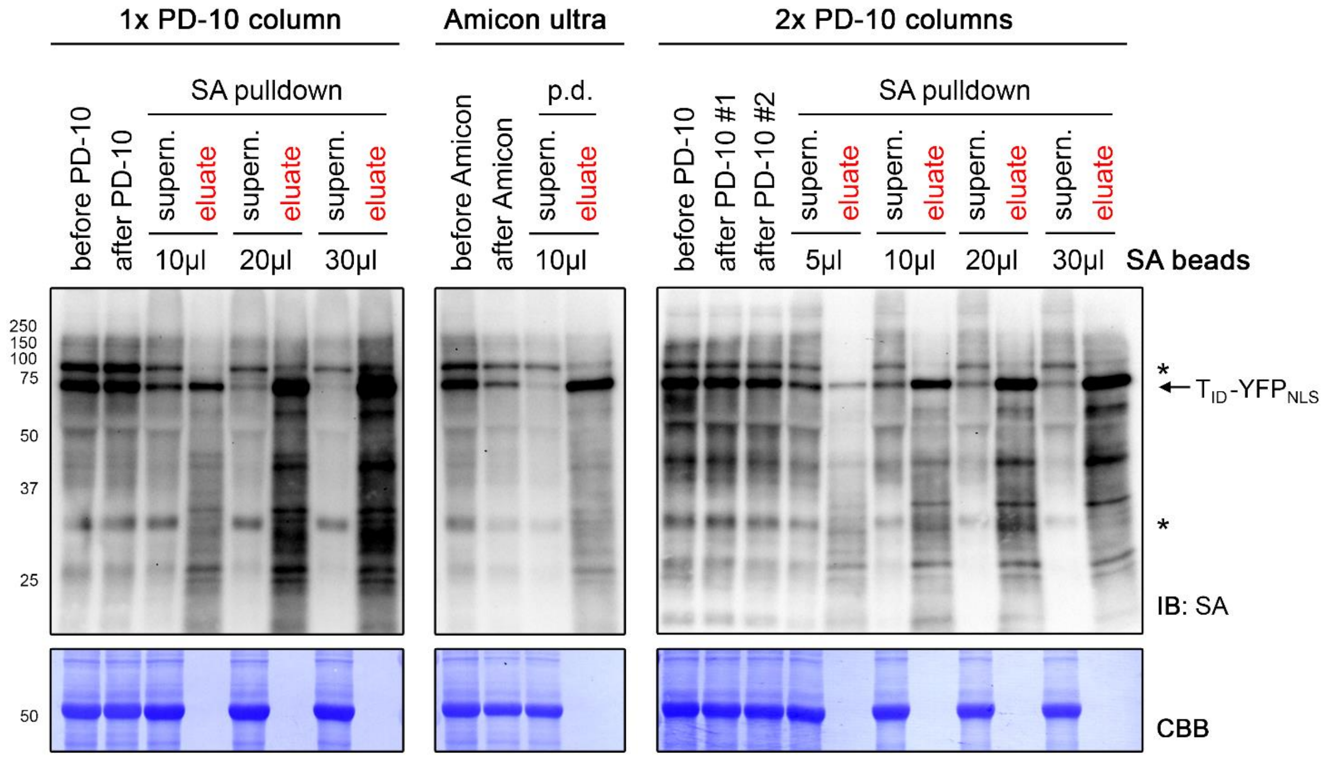


**Figure 4 – figure supplement 7: Comparison of different biotin depletion methods and bead concentrations for an effective pulldown of biotinylated proteins**

Five day old seedlings expressing the UBQ10pro::$T_{ID}$-$YFP_{NLS}$ construct were submerged in a 50 µM biotin solution for three hours and used for biotin depletion-affinity purification (AP) experiments. For biotin depletion, protein extracts were either filtered through one (1x) or two consecutive (2x) PD-10 desalting columns or concentrated repeatedly using an Amicon Ultra centrifugal filter. Samples were taken before and after this step to check for protein loss. From the biotin-depleted extracts, a volume corresponding to 1/5 of the amount used for the PL experiments shown in **Figures 4-6** was used for AP with the indicated amount of streptavidin (SA) beads. Samples of the supernatants after AP and of the proteins eluted from the beads were taken to determine the ratio of biotinylated proteins remaining in the protein extract and bound to the beads. All samples were analyzed by immunoblotting (IB) with SA-HRP. The amount of protein extracts and bead eluates loaded correspond to 0.05% and 0.5% of the total volumes, respectively. The Coomassie Brilliant Blue-stained membranes (CBB) are shown as loading controls for the IBs. Asterisks mark the positions of naturally biotinylated proteins. Biotin depletion with the Amicon Ultra Centrifugal filter, but not with PD-10 columns resulted in considerable protein loss. Increasing the amount of beads increased the AP efficiency.

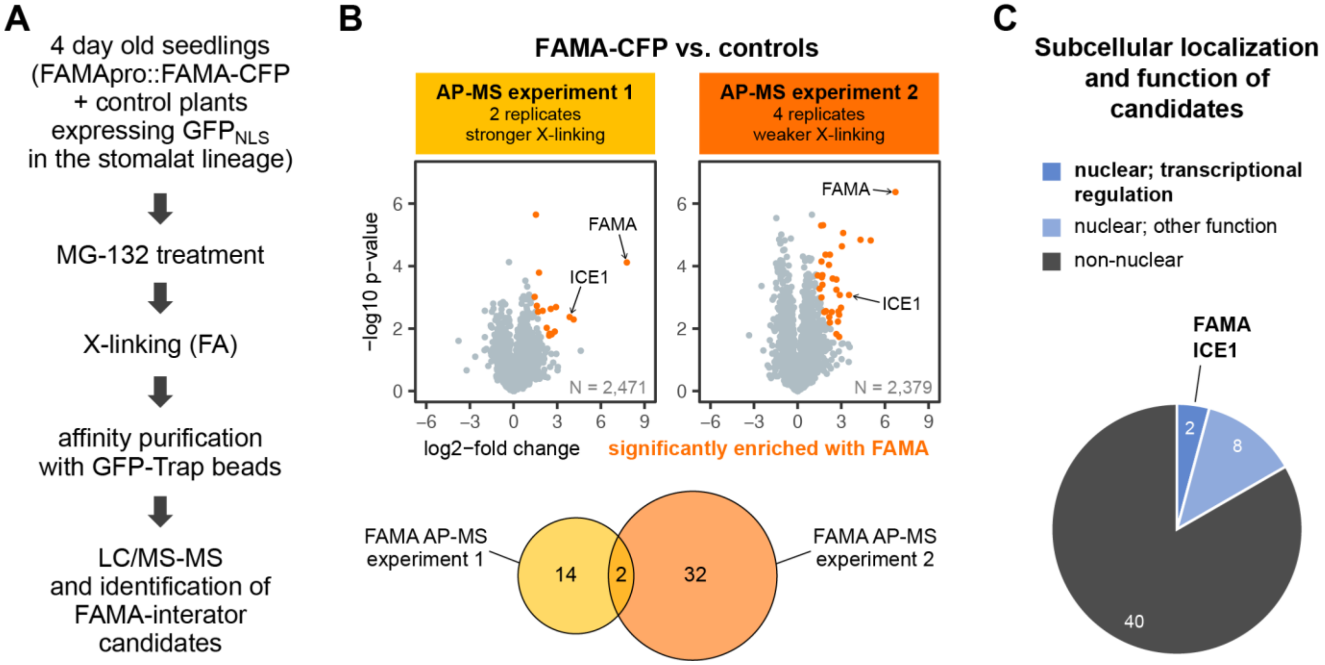


**Figure 5 – figure supplement 1: FAMA-CFP AP-MS experiments identified ICE1, but no novel transcriptional regulators as putative FAMA partners**

**(A) Simplified workflow of the AP-MS experiments.** Four day old seedlings of a FAMA-CFP line (FAMApro::FAMA-CFP in *fama-1*) and two lines expressing nuclear GFP under stomatal lineage-specific promoters (SPCHpro::GFP$_{NLS}$ and MUTEpro::GFP$_{NLS}$) were treated with MG-132 to stabilize proteins and with formaldehyde (FA) to cross-link (X-link) proteins. X-linking was done with 0.25 % FA in experiment 1 and with 0.125 % in experiment 2. Proteins extracted from 15 g of plant material per sample were used for AP with GFP-Trap beads and purified proteins were identified by liquid chromatography coupled to tandem mass spectrometry (LC-MS/MS). Two biological replicates of each line were used in experiment 1, and four of the FAMA-CFP line and two of each control in experiment 2. **(B) Significantly enriched proteins.** Scatterplots show the log2-fold change and -log10 p-value from unpaired 2-sided t-tests between FAMA-CFP samples and the controls with a permutation-based FDR for multiple sample correction (cutoff: AP-MS 1: FDR = 0.2, S0 = 0.5; AP-MS 2: FDR = 0.01, S0 = 0.5; N = number of proteins used in each test). Significantly enriched proteins are shown in orange and FAMA and its heterodimerization partner ICE1 are indicated. The Venn diagram shows the overlap between the candidate proteins identified in the two experiments (inlcuding FAMA). Only FAMA and ICE1 were enriched in the PL experiment shown in **Figure 5**. Filtering and statistical analysis was done in Perseus. **(C) Subcellular and functional distribution of candidates.** Candidates were classified into nuclear and non-nuclear proteins based on localization predictions (SUBA4 consensus prediction) and experimentally determined localization data and nuclear localized proteins were further divided into such that are involved in transcriptional regulation (2 proteins, including FAMA) or have a different function (8 proteins). For a list of identified proteins and the localization predictions see source file Figure 5 – supplemental table 5.

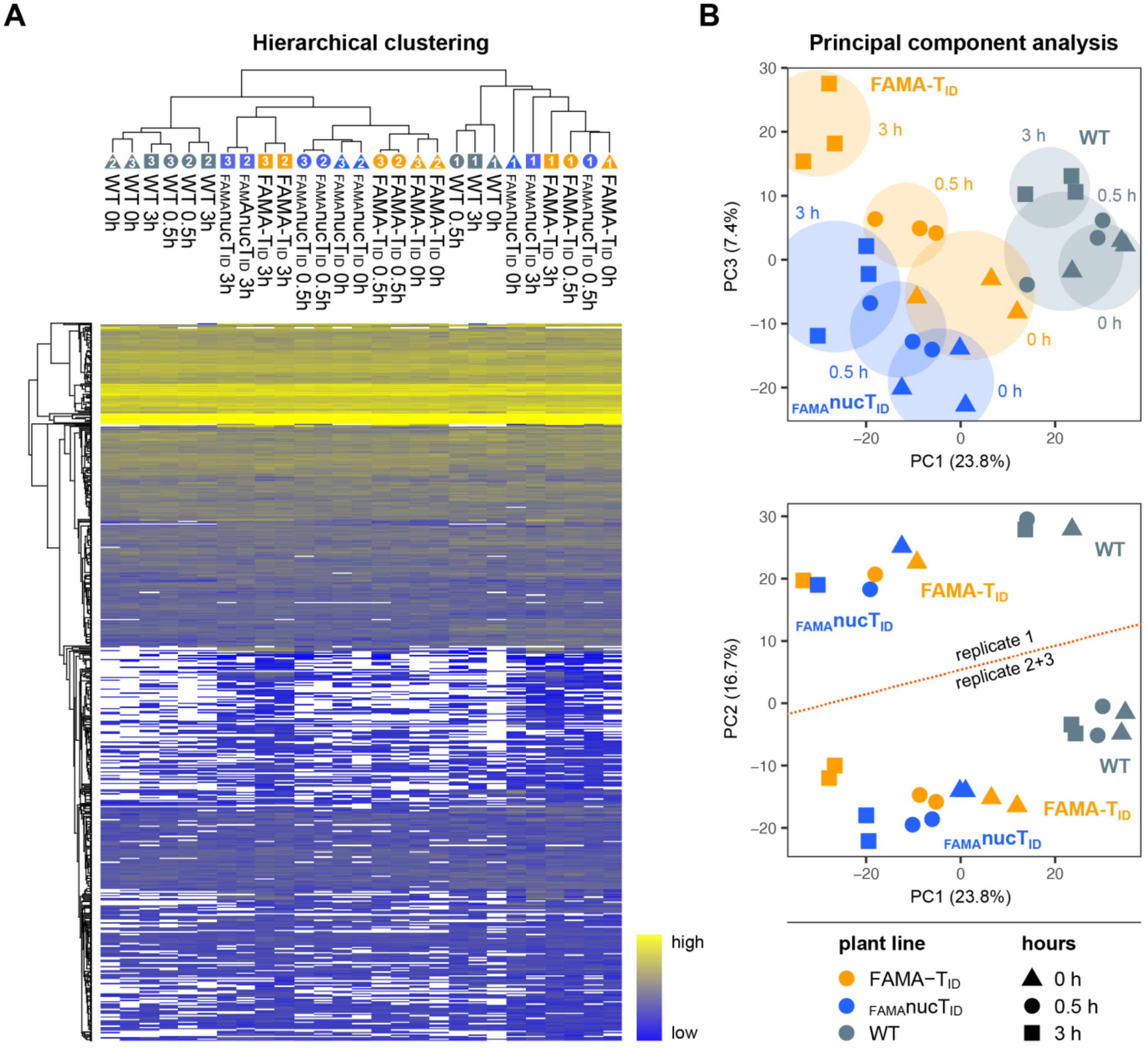


**Figure 5 – figure supplement 2: Clustering and PCA of samples for the 'FAMA complex' PL experiment**

**(A) Hierarchical clustering.** Proteins that were identified in all three replicates of at least one time point of one genotype were used for average linkage clustering with Euclidian distance. Rows and columns represent individual proteins and samples. In the sample dendrogram, genotypes, replicate numbers and time points are labeled with differently colored and numbered triangles, circles and square (same designations as in (B)). In the heatmap, proteins are colored according to their abundance (label free quantification value). White areas represent proteins not identified in a sample. **(B) Principal component analysis (PCA).** Missing value were imputed (from normal distribution) and the data matrix was used for PCA. Principal component 1 (PC1) was plotted against PC2 or 3. The PC1-PC3 plot (top) shows separation of the samples by genotype and time. Areas where samples cluster by genotypes and time point are shaded. A batch effect separating replicate 1 from replicates 2 and 3 is visible in PC2 (bottom) as well as in (A). Clustering and PCA analysis were done in Perseus using built-in functions.

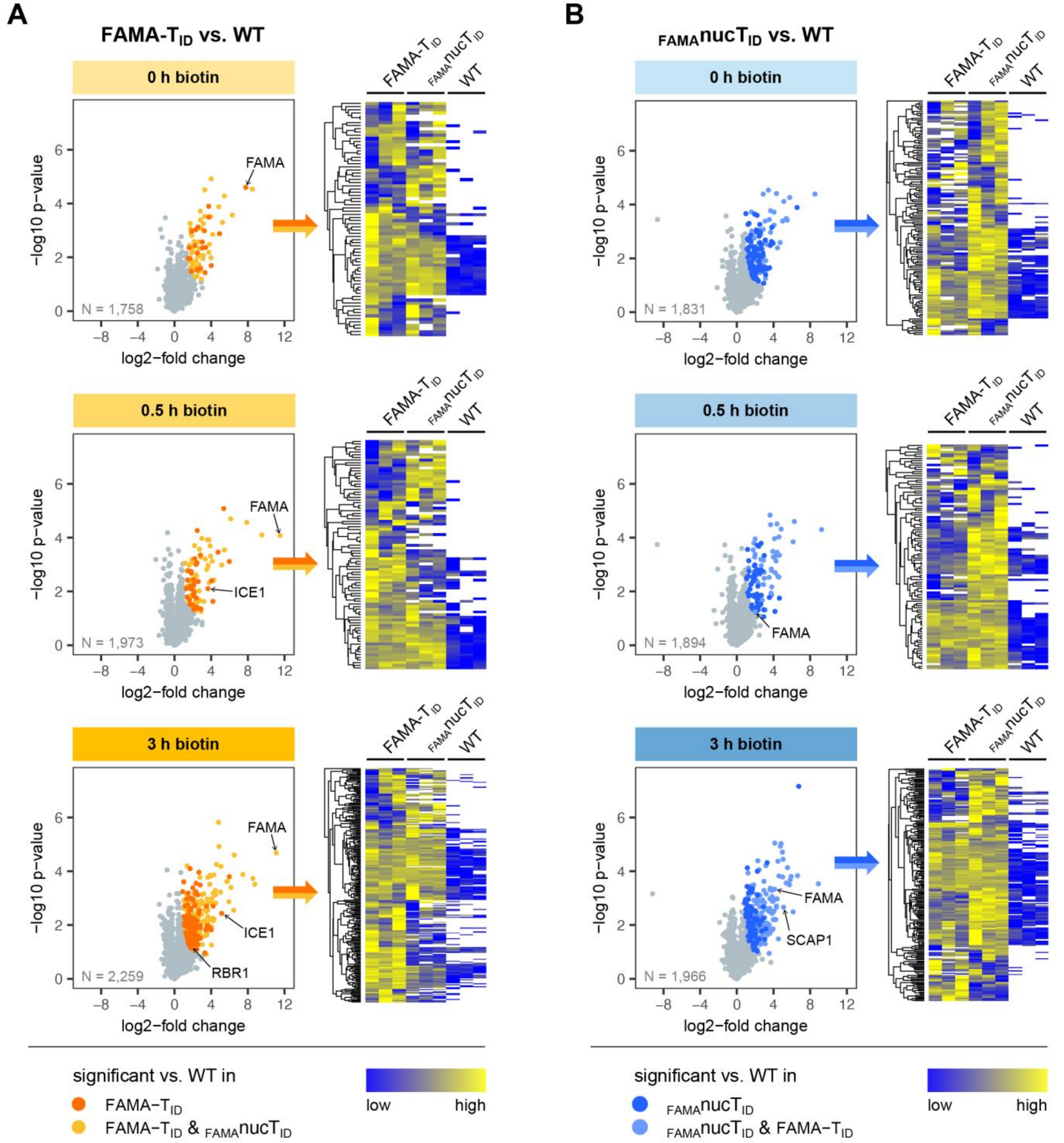


**Figure 5 – figure supplement 3: Significantly enriched proteins in the FAMA-T$_{ID}$ and $_{FAMA}$nucT$_{ID}$ lines**

For each time point, proteins that were significantly enriched in FAMA-T$_{ID}$ or $_{FAMA}$nucT$_{ID}$ compared to wild-type (WT) were determined by 2-sided t-tests with a permutation-based FDR for multiple sample correction (cutoff: FDR = 0.05, S0 = 0.5). Only proteins identified in all three biological replicates of either FAMA-T$_{ID}$ **(A)** or $_{FAMA}$nucT$_{ID}$ **(B)** at the indicated time point were used for these tests (N = number of proteins used in each test). Scatterplots on the **left** show log2-fold changes and -log10 p-values from t-tests (plots in (A) are also shown in Figure 5). Significantly enriched proteins are shown in orange/yellow (A) or dark/light blue (B) and increase with the duration of biotin treatment. Known FAMA interactors (ICE1 and RBR1) and nuclear proteins in guard cells (FAMA and SCAP1) are indicated. The heatmaps on the **right** of each scatter plot show average linkage clustering of significantly enriched proteins in this plot. Proteins are colored according to their abundance (Z-transformed label free quantification values). White areas represent proteins not identified in a sample.

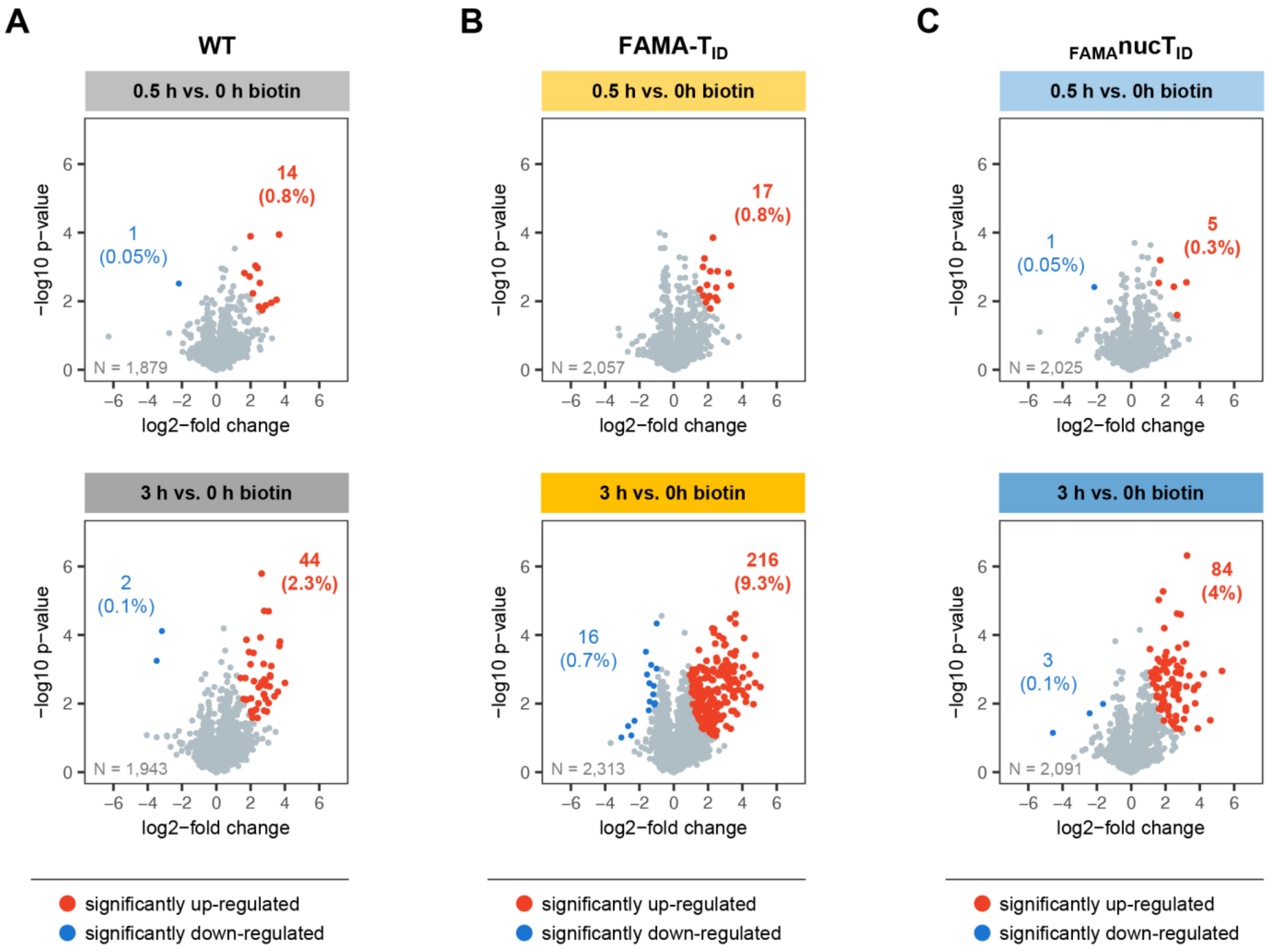


**Figure 5 – figure supplement 4: Increase of biotinylation in the PL samples over time**

Proteins that were significantly enriched in wild-type (WT), FAMA-$T_{ID}$ or $_{FAMA}$nuc$T_{ID}$ samples after 0.5 and after 3 hours of biotin treatment compared to untreated samples were determined by 2-sided t-tests with a permutation-based FDR for multiple sample correction (cutoff: FDR = 0.05, S0 = 0.5). Only proteins identified in all three biological replicates of wild-type (WT) **(A)**, FAMA-$T_{ID}$ **(B)** or $_{FAMA}$nuc$T_{ID}$ **(C)** at any of the two time points were used for the tests (N = number of proteins used in each test). Scatterplots show log2-fold changes and -log10 p-values from t-tests. Significantly up-and down-regulated proteins are shown in red and blue and their number and percentage of all analyzed proteins are given. Only few proteins are enriched in any of the genotypes after 0.5 hours of biotin treatment.

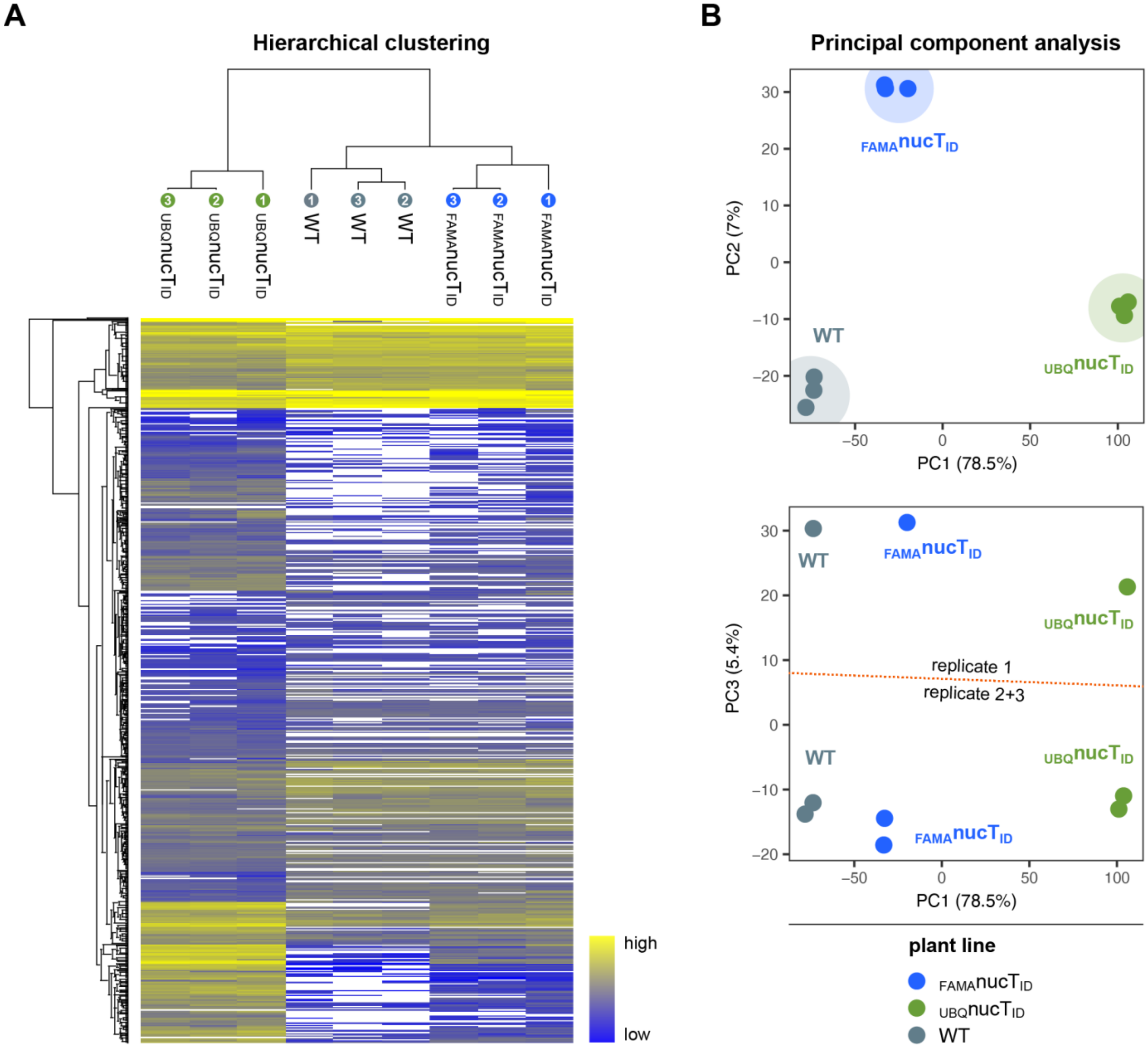


**Figure 6 – figure supplement 1: Clustering and PCA of samples for the 'nuclear proteome' PL experiment**

**(A) Hierarchical clustering.** Proteins that were identified in all three replicates of at least one genotype were used for average linkage clustering with Euclidian distance. Rows and columns represent individual proteins and samples. In the sample dendrogram, genotypes and replicate number are labeled with differently colored and numbered circles. In the heatmap, proteins are colored according to their abundance (label free quantification value). White areas represent proteins not identified in a sample. **(B) Principal component analysis (PCA).** Missing value were imputed (from normal distribution) and the data matrix was used for PCA. Principal component 1 (PC1) was plotted against PC2 or 3. The PC1-PC2 plot (top) shows clear separation of the samples by genotype. Areas where samples cluster are shaded. A batch effect separating replicate 1 from replicates 2 and 3 is visible in PC3 (bottom). Clustering and PCA analysis were done in Perseus using built-in functions.

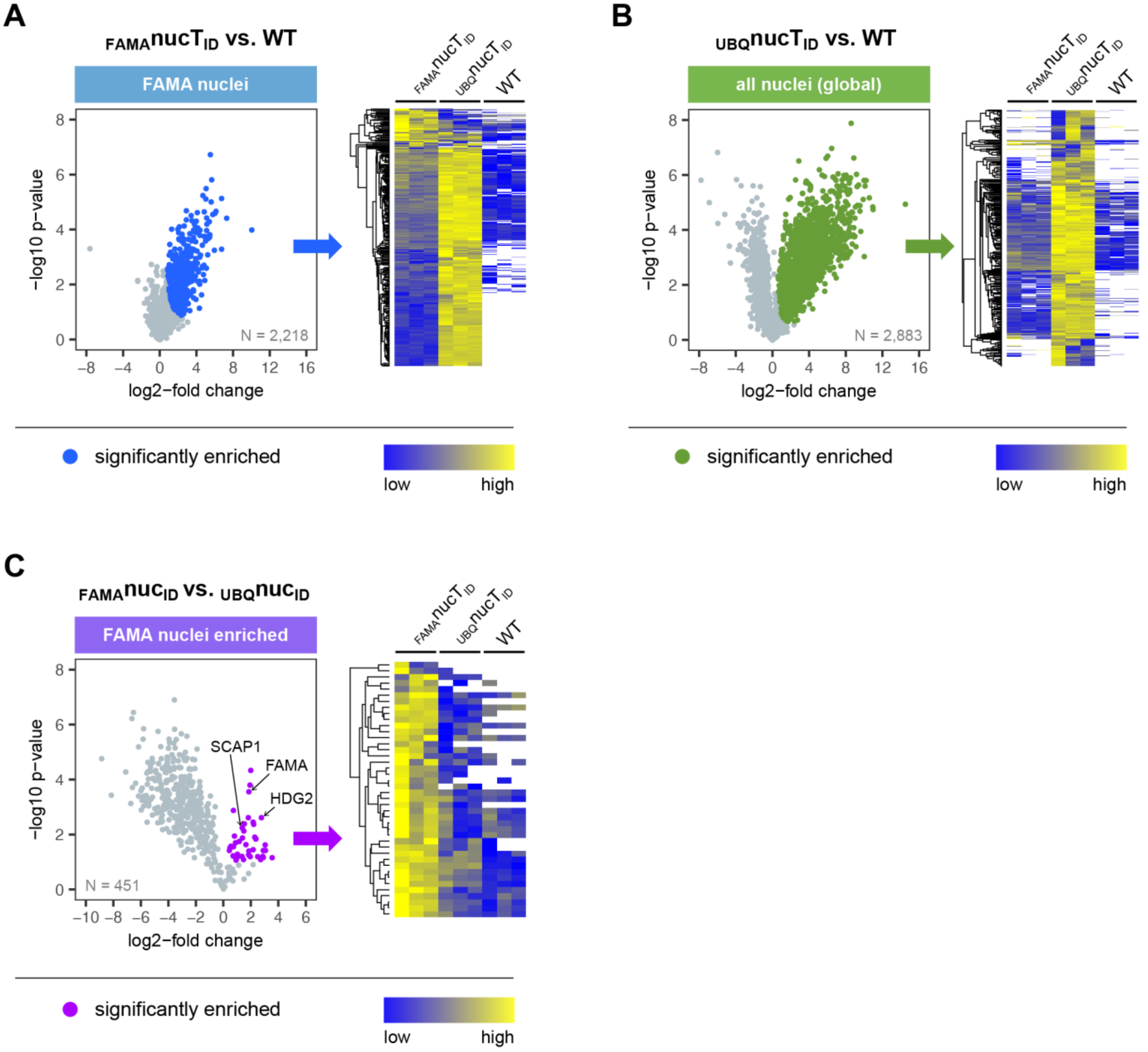


**Figure 6 – figure supplement 2: Significantly enriched proteins in FAMA-expressing and all nuclei**

Proteins that were significantly enriched in $_{FAMA}$nucT$_{ID}$ or $_{UBQ}$nucT$_{ID}$ compared to wild-type (WT) were determined by 2-sided t-tests with a permutation-based FDR for multiple sample correction (cutoff: FDR = 0.05, S0 = 0.5). Only proteins identified in all three biological replicates of $_{FAMA}$nucT$_{ID}$ **(A)** or $_{UBQ}$nucT$_{ID}$ **(B)** were used for these tests (N = number of proteins used in each test). Proteins that were enriched in FAMA nuclei compared to all nuclei **(C)** were determined by a 2-sided t-tests with an FDR of 0.01 (S0 = 0) using only proteins significantly enriched in $_{FAMA}$nucT$_{ID}$ compared to WT. Scatterplots on the **left** are from Figure 6 and show log2-fold changes and -log10 p-values from t-tests. Significantly enriched proteins are shown in blue (A), green (B) or purple (C). Known FAMA-nuclei enriched proteins (FAMA, SCAP1, HDG2) are indicated. The heatmaps on the **right** of each scatter plot show average linkage clustering of significantly enriched proteins in this plot. Proteins are colored according to their abundance (Z-transformed label free quantification values). White areas represent proteins not identified in a sample.

Enriched GO terms for the global nuclear proteome ($_{UBQ}nucT_{ID}$ enriched)

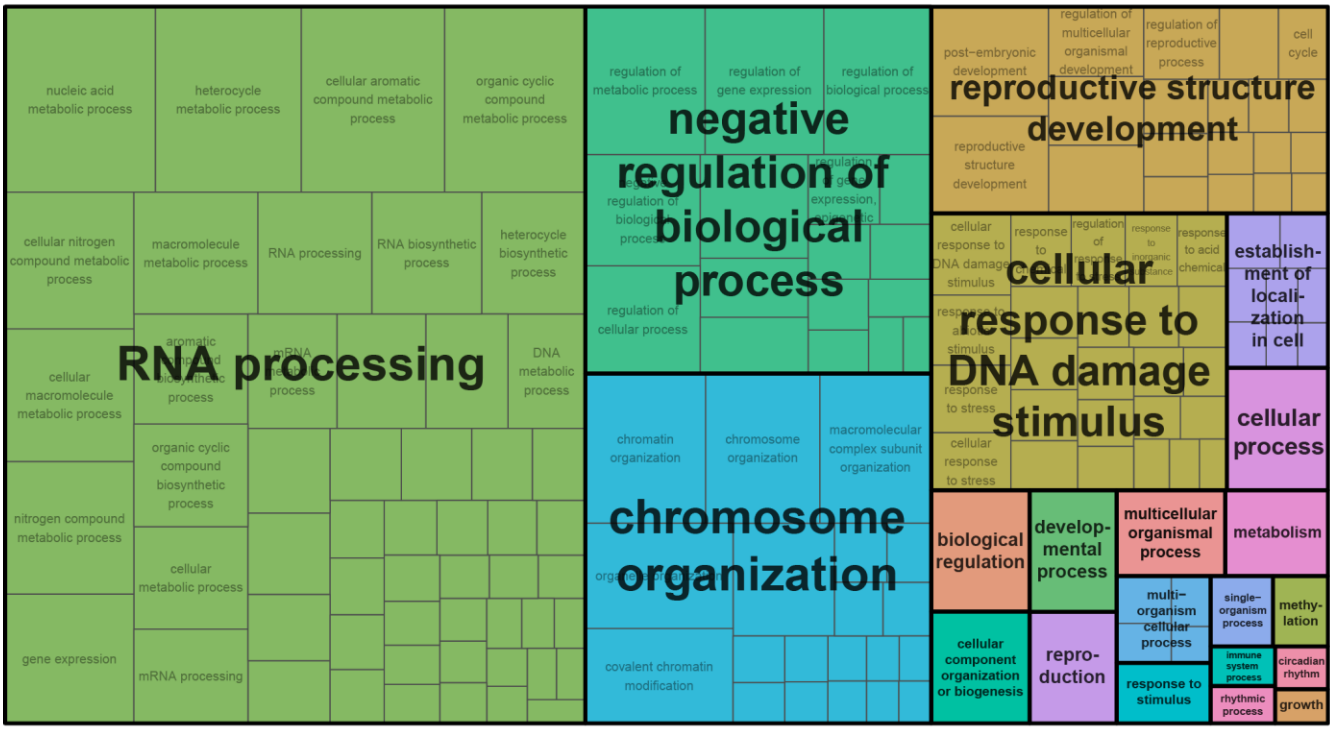


Enriched GO terms for the young guard cell nuclear proteome ($_{FAMA}nucT_{ID}$ enriched)

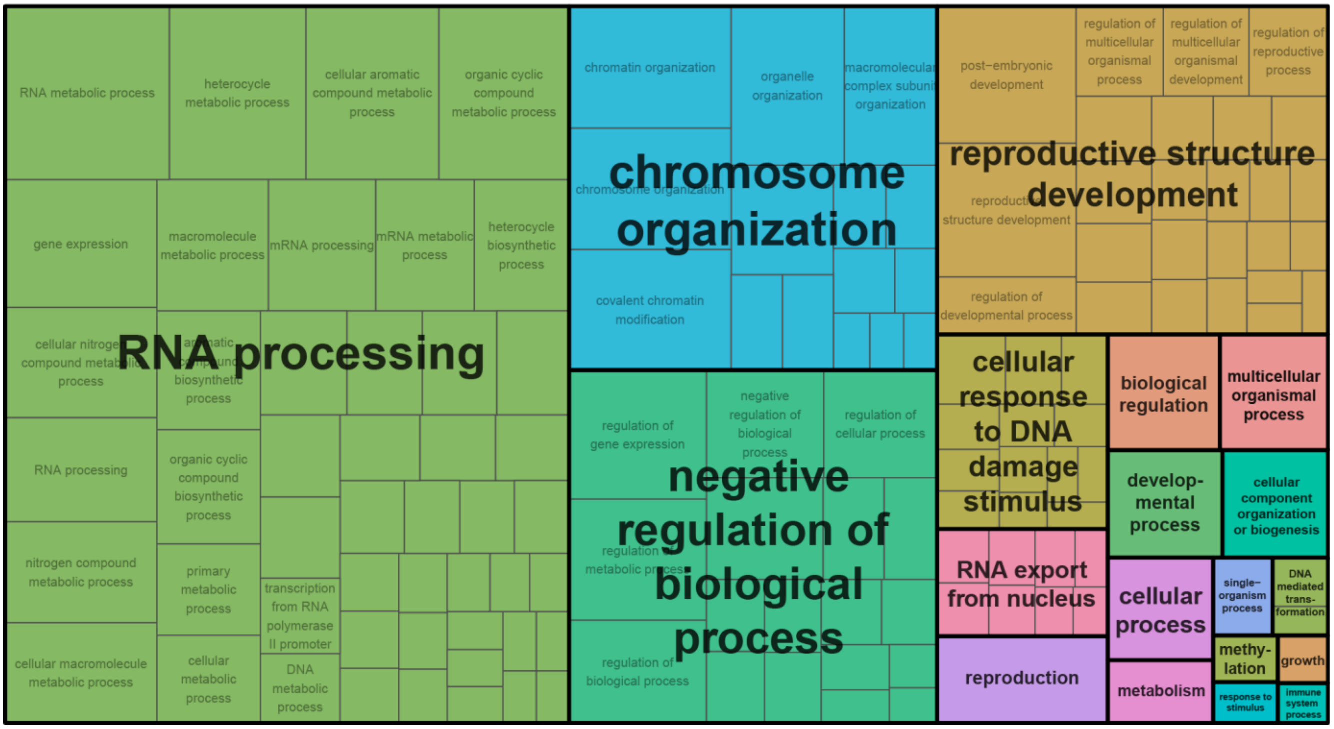


**Figure 6 – figure supplement 3: GO terms enriched in global and FAMA nuclear proteomes**

GO terms enriched for global and FAMA nuclear proteins (Figure 6) were determined with AgriGO v2 and visualized with REViGO. Shown is the TreeMap plot of enriched 'biological processes' in global (UBQ10 promoter, top) and FAMA (bottom) nuclei. Related GO-terms are clustered into same colored boxes and given summary titles.
